# Detecting polygenic adaptation in admixture graphs

**DOI:** 10.1101/146043

**Authors:** Fernando Racimo, Jeremy J. Berg, Joseph K. Pickrell

## Abstract

An open question in human evolution is the importance of polygenic adaptation: adaptive changes in the mean of a multifactorial trait due to shifts in allele frequencies across many loci. In recent years, several methods have been developed to detect polygenic adaptation using loci identified in genome-wide association studies (GWAS). Though powerful, these methods suffer from limited interpretability: they can detect which sets of populations have evidence for polygenic adaptation, but are unable to reveal where in the history of multiple populations these processes occurred. To address this, we created a method to detect polygenic adaptation in an admixture graph, which is a representation of the historical divergences and admixture events relating different populations through time. We developed a Markov chain Monte Carlo (MCMC) algorithm to infer branch-specific parameters reflecting the strength of selection in each branch of a graph. Additionally, we developed a set of summary statistics that are fast to compute and can indicate which branches are most likely to have experienced polygenic adaptation. We show via simulations that this method - which we call PolyGraph - has good power to detect polygenic adaptation, and applied it to human population genomic data from around the world. We also provide evidence that variants associated with several traits, including height, educational attainment, and self-reported unibrow, have been influenced by polygenic adaptation in different populations during human evolution.

## Introduction

There is much interest in identifying the individual genetic variants that have experienced natural selection during recent human evolution. Many popular methods tackle this problem by identifying alleles that have changed frequency faster than can be explained by genetic drift alone, and that can instead be explained by selective processes. These methods exploit patterns like haplotype homozygosity [1, 2, 3] and extreme population differentiation [4, 5, 6], and have yielded several important candidates for human adaptation: for example, *LCT* [7], *EDAR* [2], *EPAS1* [4] and the *FADS* region [8]. In order for these signals to be detectable at the level of an individual locus, the historical changes in allele frequency must have been large and rapid. Therefore, they can only be produced by alleles that confer a strong selective advantage.

With the advent of large-scale GWAS for a variety of measurable traits, however, it has now become possible to detect a more subtle mechanism of adaptation. If a trait is polygenic, positive selection may instead occur by concerted shifts at many loci that all contribute to the variation in a trait. Over short time scales, these shifts are expected to be small (but see [9] for polygenic dynamics under longer time scales). They are also expected to occur in consistent directions, such that alleles that increase the trait will systematically rise in frequency (if selection favors the increase of the trait) or fall in frequency (if selection operates in the opposite direction). None of the allele frequency changes need to be large on their own for the phenotypic change to be large. This process is called polygenic adaptation and may underlie major evolutionary processes in recent human history [10, 11, 12, 13].

A number of methods have been developed to detect polygenic adaptation using loci identified from GWAS. Turchin et al. [14] was the first such study. They developed a test for polygenic adaptation between two populations, and showed that there were systematic frequency differences at height-associated loci between northern and southern Europeans, which could not be explained by genetic drift alone. Berg and Coop [15, 16] developed a more general method to detect polygenic adaptation by testing for over-dispersion of mean genetic values among several populations, using a genome-wide population covariance matrix to predict how alleles should behave under neutrality. Robinson et al. [17] used theory by Ovaskainen et al. [18] to develop a similar population differentiation method to detect polygenic adaptation with GWAS. They also made use of the genome-wide covariance matrix, but, in contrast to Berg et al., their method is implemented in a Bayesian linear mixed model.

None of these methods require a detailed model of human history to detect polygenic adaptation. Their use of the genome-wide covariance matrix allows them to capture patterns of genetic drift among populations without having to infer their history. While this makes them quite powerful, it also means that they are not very useful at determining where and when polygenic adaptation took place in the past.

Here, we develop a method to detect polygenic adaptation that uses a more parameter-rich model of historical population structure: an admixture graph, which is a simplified representation of the history of divergences and admixture events among populations [19, 20]. An explicit model allows us to infer where particular bouts of polygenic adaptation took place in human history, so as to better understand how selection on trait-associated variants has occurred over past generations.

## Results

### Model

Assume we have measured genotypes at a SNP that influences a trait in a set of *M* populations. Let *d_m_* be the count of the derived allele in population *m* and let 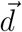 be the vector across the M populations of each *d_m_* observation. Let *n_m_* be the total number of chromosomes observed in population *m* (together 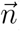). Assume we have an admixture graph *G* relating these populations, and that this graph consists of an accurate topology, as well as accurate branch lengths and admixture rates. Branch lengths are in units of drift, which are approximately equal to *t*/2*N_e_*, where, for each branch, *t* is the number of generations and *N_e_* is the effective population size, assuming *t* ≪ *N_e_* [21]. In practice, we can estimate such a graph from neutral genome-wide data, and we use the program *MixMapper* [22] when applying our method to real data below.

We wish to model the changes in frequency of the trait-associated allele over the graph *G*. At each node in the graph, we introduce a parameter that corresponds to the allele frequency of the variant at that node. Let *f_R_* be the derived allele frequency at the root of the graph, 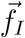 be the vector of allele frequencies at all the other internal nodes of the graph, and 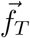 be the vector of allele frequencies at the tips of the graph.

The probability of the parameter values and the data can then be decomposed as follows:

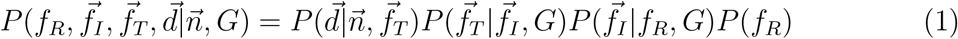

We now take each of terms above in turn. First, the probability of the observed counts is simply a product of binomial probabilities:

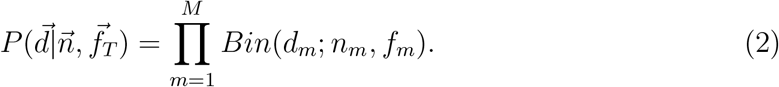

where *f_m_* is the element of 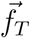 that corresponds to population *m*.

To get the probabilities of the changes in allele frequency across different nodes, consider a single branch of *G*. Assuming the branch is relatively short (such that the allele does not approach fixation or extinction in this time period) and there is no natural selection, we can use the Normal approximation to the Wright-Fisher diffusion [23, 24, 25] to model the allele frequency at the descendant node of the branch (*f_D_*) as a function of the allele frequency at the ancestral node (*f_A_*):

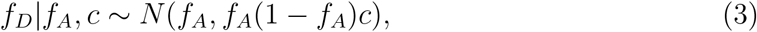

where *c* corresponds to the amount of drift that has occurred in the branch. In practice, we use a truncated Normal distribution with point masses at 0 and 1, to account for the possibility of fixation or extinction of the allele [26].

In our model there may have also been selection on the allele on the branch, such that it was pushed to either higher or lower frequency because of its influence on a trait. We can model the selected allele frequency by modifying the infinitesimal mean of the Wright-Fisher diffusion and approximating the diffusion with a Normal distribution that now includes some additional terms ([14, 26, 27], Bhérer et al. in prep.):

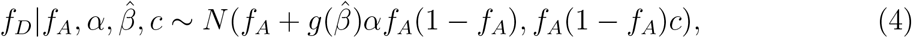

where 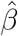 is the effect size estimate at that site (defined with respect to the derived allele), 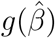 is some function that relates the effect size estimate to the selective pressure and *α* is our positive selection parameter, which is approximately equal to the product of the selection coefficient for the advantageous allele and the duration of the selective process [14, 28]. In practice, we will set 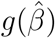 to be equal to the sign (+1 or −1) of 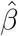 (Figure 2), so as to avoid giving too much weight to variants of strong effect. We will model selection only on SNPs that are associated with a trait in a particular GWAS.

**Figure 1:**
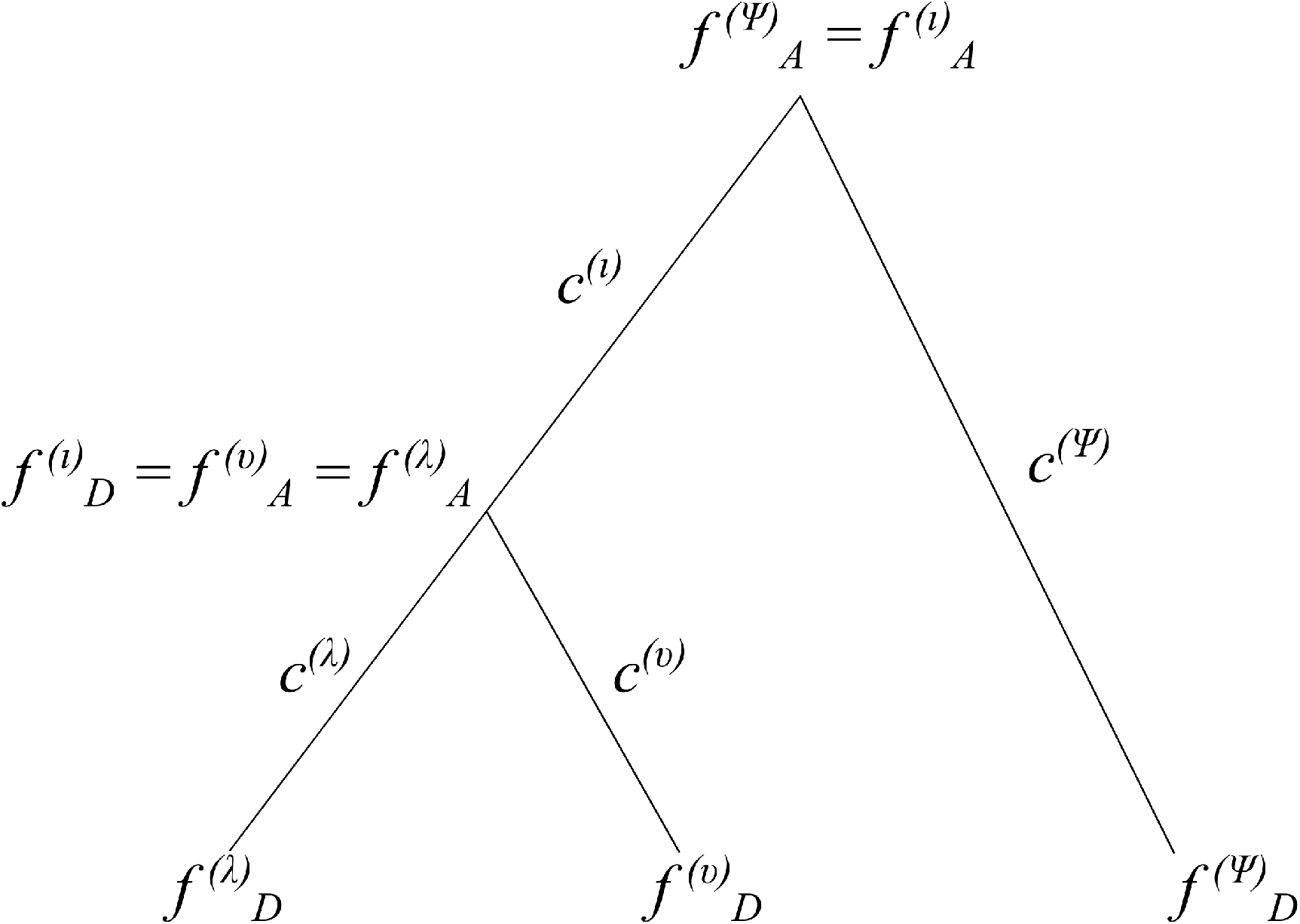
Schematic of drift values (*c*) and allele frequencies (*f*) for a 3-leaf population tree with no admixture.

**Figure 2:**
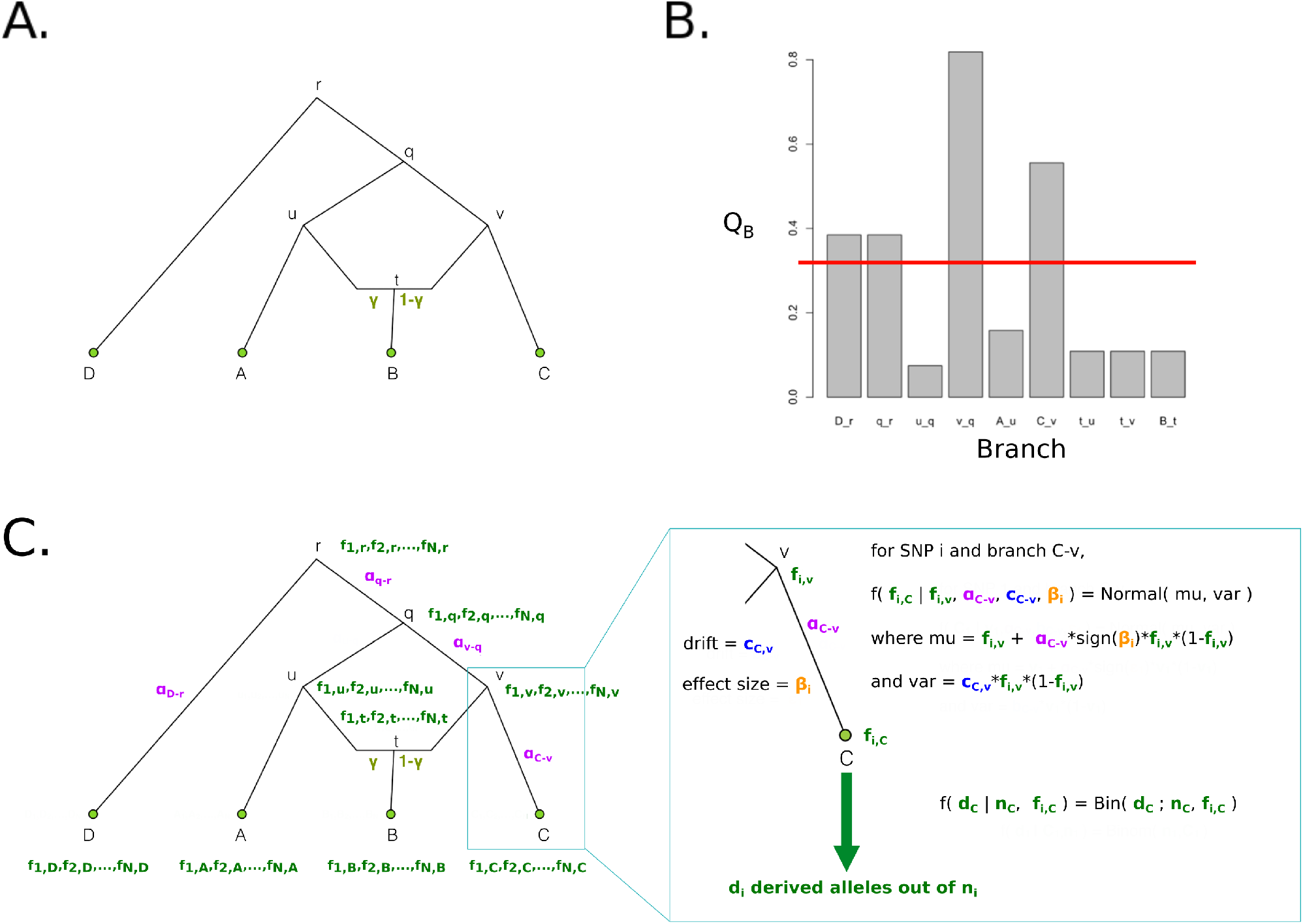
Schematic of PolyGraph estimation procedure for a 4-population graph with one admixture event. Panel A: The first step is the estimation of the admixture graph topology using neutral SNP data, via an admixture-graph fitting program like *MixMapper*. Panel B: Then, we use the *Q_B_* statistic to determine which branches to explore in the MCMC. Selection parameters whose corresponding branches have a *Q_B_* statistic that is smaller than a specific cutoff (red line) are set to a fixed value of 0 in the MCMC. Panel C: Model for MCMC sampling. The SNP frequencies in the nodes of the graph are shown in green, while the selection parameters for each candidate branch are shown in purple. For each SNP, the likelihood of each branch of the graph is a Normal distribution. To model the sampling of derived alleles in the leaves of the graph, we use a binomial distribution.

We can calculate the probability of these parameters at a particular site as the product of the Normal probability densities that correspond to the evolution of allele frequencies down each branch times a binomial probability density to account for sampling error. Let us denote the Normal density that corresponds to a particular branch *λ* as 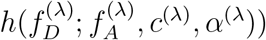, where we suppress notation of 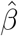 for clarity. Then, for example, the probability of a given pattern of allele frequencies and sample counts over a rooted 3-leaf tree with 4 branches *λ, ι, υ, ψ* (Figure 1) can be computed as follows:

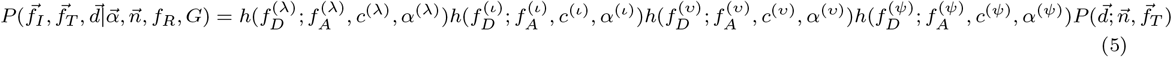

when the *α* parameters and the allele frequency at the root of the tree 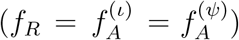 are known. Note that some of the symbols here correspond to the same allele frequencies. For example, if the *υ* branch is one of the immediate descendant branches of the *ι* branch, then 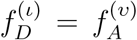. Assuming SNPs are unlinked, we can compute the probability of the allele frequency configurations at all N trait-associated SNPs as a product over the probabilities at each of the SNPs:

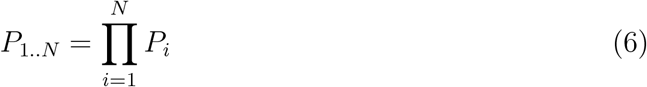

where *P_i_* is the probability of the parameters of interest at SNP *i* under our tree model.

More generally, we can also compute the probability of our parameters in an admixture graph, containing nodes with more than one parent. In that case, the probability of an allele frequency of an admixed node is a weighted sum of the probability paths corresponding to its two parents, where the weights are the admixture rates for each of the two contributions.

In practice, for a given SNP, we know neither the allele frequencies at the inner nodes 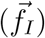 nodes, at the tip nodes 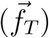 and at the root node (*f_R_*), nor the *α* parameters in each branch 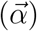. We want to obtain a posterior distribution of these parameters, given the data and the known graph: 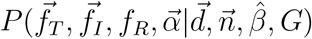. We aim to do this for all trait-associated SNPs. We therefore developed an MCMC sampler to transition between the states of these variables and estimate their posterior distribution (Figure 2).

We set the prior for the frequencies at the root *f_R_* to be a uniform distribution. For a given SNP i,

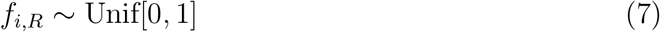

As there are many combinations of *α* parameters that generate almost equivalent likelihoods in a complex admixture graph (Figure S1), we use a “spike-and-slab” prior for the *α* parameters, so as to promote sparsity. For a given branch *j*,

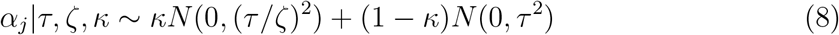

This is a mixture of two Normal distributions centered at 0: one of the distributions has a wide standard deviation (*τ*), while the other has a much narrower standard deviation, which is a fraction (1/*ζ*) of *τ*, and approximates a point mass at 0 [29, 30]. Here, *κ* is the mixture probability of drawing from the narrower Normal distribution, and we model it with a uniform hyperprior (see Materials and Methods). The idea behind this is that our assumed prior belief is that only a few of the branches in the admixture graph have experienced bouts of polygenic adaptation, so we reward *α* parameters that tend to stay in the neighborhood of 0 during the MCMC run.

### A statistic for prioritizing branches

We observed via simulations that different combinations of *α* parameters can produce very similar likelihood values. This causes the MCMC sampler to explore different combinations of *α* values in the same posterior run, when only one such combination is actually correct (Figure S1). The aforementioned spike-and-slab prior serves to partially ameliorate this problem, but we aimed to find a way to further encourage sparsity by reducing the possible number of candidate branches that are explored in the MCMC.

We therefore devised a set of summary statistics that can be computed before starting an MCMC run and are meant to detect branch-specific deviations from neutrality. Let **F** be the empirical population covariance matrix, which - under ideal neutral conditions - should be determined by the admixture graph connecting all the populations. Let 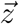 be the mean-centered vector of estimated mean genetic values for each of the M populations, computed from the *N* SNPs that are known to be associated with a trait. For a specific population *m*:

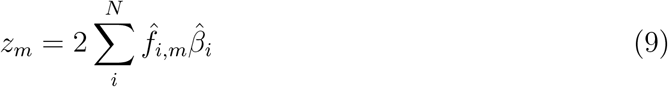

Here, 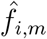 is the sample frequency of SNP *i* in a panel of population *m*, and *βi* is its effect size estimate. The vector *z* therefore contains the *z_m_* values for all *M* populations.

Furthermore, let us define the vector 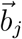 for a particular branch *j* to be equal to the contribution *w* of that branch to each of the leaves of the graph. For example, branch v-q in Figure 2 is a full ancestral branch to leaf C, a partial ancestral branch to leaf B (due to the admixture event), but not an ancestral branch to either A or D. Therefore, the vector 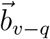 is equal to (*w_A_, w_B_, w_C_, w_D_*) = (0,1 – *γ*, 1, 0). By the same reasoning, 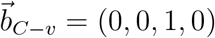 and 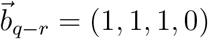.

Now, let:

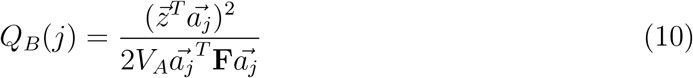

where 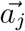 is a scaled version of 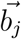 so that is has unit length. The numerator of this statistic is proportional to the squared covariance between the genetic values and the vector representing the contribution of drift down branch *j* to the overall pattern of among population divergence. The denominator gives the expectation of the numerator under the neutral model. This statistic reflects how much of the deviations from neutrality among the population mean genetic values for a trait is due to branch *j*. One can show (see Materials and Methods) that *Q_B_*(j) has a 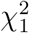 distribution under a null model of multivariate Normal drift, for any branch *j*, and excessively large values of *Q_B_*(j) therefore represent evidence suggesting non-neutral evolution down branch *j*.

We used this summary statistic (computed for each branch) to prioritize which branches to explore in our MCMC. We applied a prior point mass of zero to all branches whose corresponding *Q_B_* statistic were smaller than a particular cutoff. The choice of cutoff was based on simulations (see Materials and Methods). The MCMC only produces posterior samples for branches that pass this cutoff and are therefore highly deviated from their expectation under neutrality. We also update the *α* parameters of these latter branches in our MCMC with a frequency proportional to their *Q_B_* values, in a similar fashion to [31].

### Implementation

We implemented both the MCMC and the *Q_B_* statistic computation in a program that we call PolyGraph. We use the R packages *admixturegraph* [32] and *igraph* [33] to visualize and manipulate various aspects of a graph. The R scripts to run PolyGraph can be downloaded here: https://github.com/FerRacimo/PolyGraph

### Simulations

To assess the performance of our method, we simulated different demographic scenarios, under a Wright-Fisher binomial sampling model (see Materials and Methods). We first simulated a simple three-leaf tree with four branches, in which the sampled panels in the leaves were each composed of 100 diploid individuals (Figure 3.A). We tested scenarios of different branch lengths: each of the branches was simulated to be either of length 0.02 or of length 0.05. We also tested different types of branch under selection (either a terminal branch or an internal branch). We additionally tested a four-leaf admixture graph with one admixture event (Figure 3.C) in these same scenarios. For comparison, the amount of genetic drift between Spanish and French human populations is 0.016 and the amount of drift between French and Han Chinese human populations is 0.22 [34]. The latter is approximately equal to the drift separating populations A and C in our 4-population graph, when each branch has length equal to 0.05.

**Figure 3:**
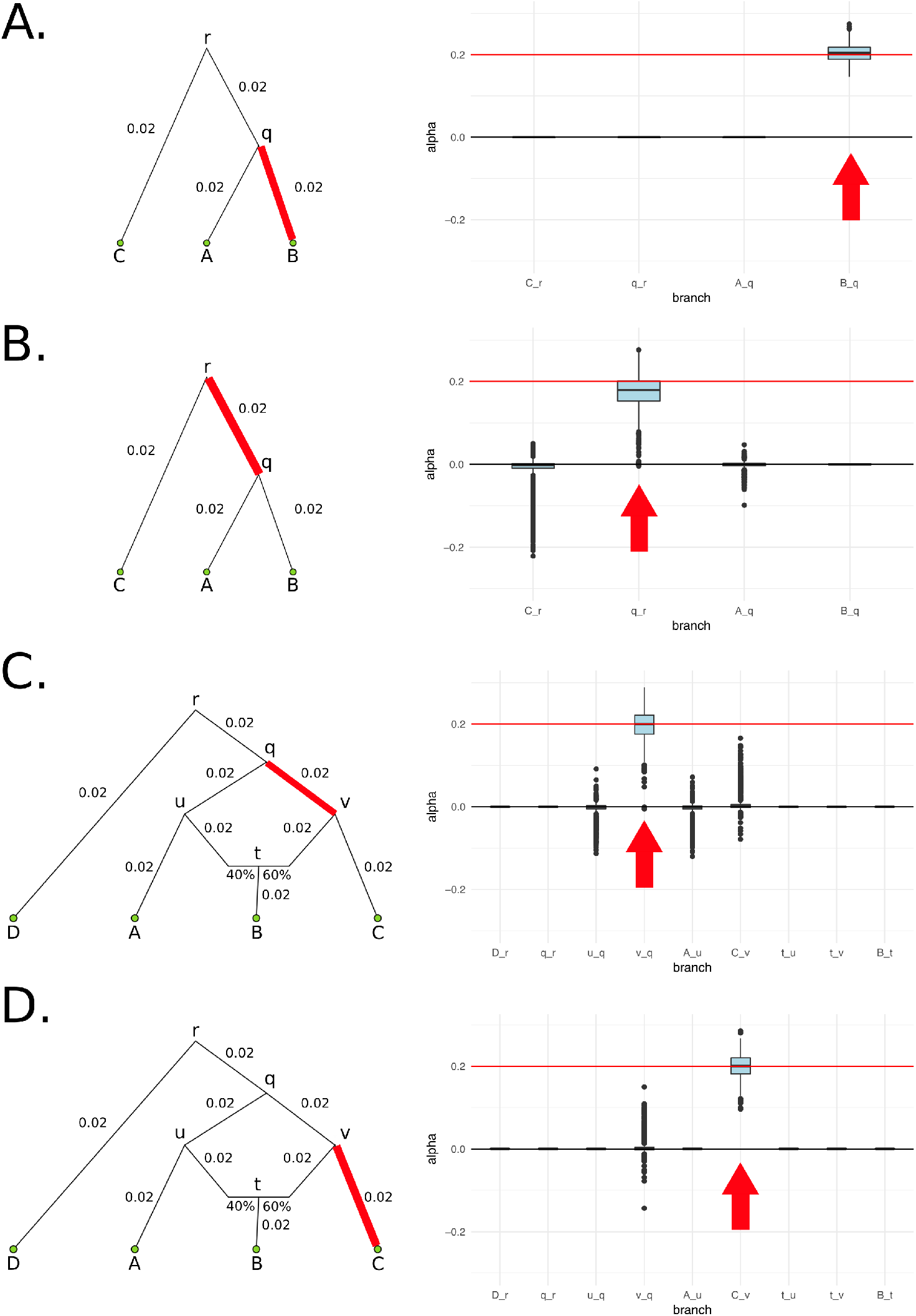
We simulated 400 SNPs affecting a trait under polygenic adaptation, and then used our MCMC to obtain posterior distributions of the *α* parameters for each branch. The red arrows denote the selected branch. The red line in the box-plots denotes the simulated value of *α* (in this case, 0.2). The lower, middle and upper hinges denote the 25th, 50th and 75th percentiles, respectively. The upper whisker extends to the highest value that is within 1.5 * IQR of the upper hinge, where IQR is the inter-quartile range. The lower whisker extends to the lowest value within 1.5 * IQR of the lower hinge. Data beyond the whiskers are plotted as points. The numbers in each graph denote the drift lengths and admixture proportions. A) Three-leaf tree with selection in a terminal branch (B-q). B) Three-leaf tree with selection in an internal branch (q-r). C) Four-population admixture graph with selection in an internal branch (v-q). D) Four-population admixture graph with selection in a terminal branch (C-v).

For all simulation scenarios, we tested five 400-SNPs replicates. We ran the MCMC four times on each simulation to check that it was behaving consistently. We observed that the runs for each simulation were very similar to each other, so we only show one run for each simulation.

When the *α* parameter is large (*α* = 0.2), the MCMC performed very well (Figure 3). For a tree (Figures S2 and S3) or a graph (Figures S4 and S5) with small branch lengths (0.02), the branch simulated to be under selection was included as a potential candidate branch in the MCMC in all simulations, indicating that the QB cutoff was not overly stringent. PolyGraph then consistently converged on the appropriate joint distribution of selection parameters. When the branches were simulated to be longer (0.05), the MCMC performed well, but, in a few simulations, it produced positive estimates for *α* parameters in neutral branches or failed to find evidence for selection in any branch (Figures S6 to S9). This occurred more often when the *α* parameter was simulated to be smaller (*α* = 0.1), but again, this was less of a problem with short-branch graphs (Figures S10 to S13) than with long-branch graphs (Figures S14 to S17). In general, we conclude that the method performs best when selective pressures are strong and/or exerted over long time periods (i.e. large *α*), and when drift parameters are small. We also observe that using a non-sparse prior (i.e. setting *κ* to 0) leads to strong mis-estimation of parameters when selection is concentrated on a single branch (Figure S18).

We were concerned about false positive estimates of selection when the graph is misspecified. To assess this, we simulated a graph like the one shown in Figure 3.C but with no selection. We first run PolyGraph while correctly specifying the topology and the branch lengths (of length equal to 0.02) as input (Figure S19), and observed that all posterior *α* estimates are tightly centered at 0, as expected. Then, we simulated a graph with the same topology but with each branch having length equal to 0.03, while incorrectly under-estimating the length of each branch to still be equal to 0.02 (Figure S20). Finally, we simulated the same graph but with each branch having length equal to 0.04 while incorrectly under-estimating the branch lengths to all be equal to 0.02 (Figure S21). With increasingly stronger misspecification of the branch lengths, we observe that the behavior of some of the posterior estimates becomes more erratic. Visual inspection of the MCMC trace indicates that underestimation of the branch lengths makes the chain to become more “sticky”, causing some parameters to get stuck at incorrect areas of parameter space for long periods of time

We also simulated a neutral graph as in Figure 3.C but pretended that population A had not been sampled, and that the graph was (incorrectly) estimated to be a 3-population tree like the one in Figure 3.A. This topological misspecification slightly affected the inference of neutrality in one of the five simulations (Figure S22), and we do not discard the possibility of other incorrect types of topologies that could also generate wrong inferences. We therefore stress that the admixture graph - especially the branch lengths - relating the populations under study should be correctly estimated before running PolyGraph. We also advise to run the MCMC only when there is significant evidence for selection based on the *Q_X_* statistic [15], which does not need an admixture graph as input, as it uses the full covariance matrix to model the expected amount of drift separating each of the populations.

### Application to 1000 Genomes data

We tested our method on sets of associated variants from 43 GWAS on 42 different traits (Table S1; two of the GWAS are for age at menarche) that were previously assembled as part of a meta-analysis studying the genetic correlations between such traits [35]. The meta-analysis split the genome into approximately independent linkage disequilibrium blocks [36]. The blocks were computed using the European populations in Phase 1 of the 1000 Genomes Project, but we observe virtually no differences in genetic scores when using the East Asian blocks instead. For each block with a posterior probability > 90% of containing an association (obtained from *fgwas* [37]), the SNP with the maximum posterior probability of being the causative variant was extracted. These SNPs can be found on github (https://github.com/PickrellLab/gwas-pw-paper/tree/master/all_single), and we downloaded them to build our candidate trait-associated SNPs. We used the VCF data from the 1000 Genomes Project [38] and built admixture graphs using *MixMapper* [22]. We excluded SNPs for which the ancestral allele in the 1000 Genomes data was unknown or unsure (lower case in VCF file). Because *MixMapper* cannot distinguish between the two drift values corresponding to two admixing branches with a common child node and the drift value specific to the immediate descendant branch of the child node (Figure 2 in ref. [22]), we forced the drift in the two admixing branches to be equal to 0.001 and assigned the drift estimated by *MixMapper* to the branch immediately descending from the child node.

We started by fitting a 7-leaf tree without attempting to model any admixture events. The tree included diverse populations sampled across the world (Figure S23.A): Nigerian Esan (ESN), Sierra Leone Mende (MSL), Northern Europeans from Utah (CEU), Southern Europeans from Tuscany (TSI), Dai Chinese (CDX), Japanese (JPT) and Peruvians (PEL). We took trait-associated variants to be under polygenic adaptation if the P-value for the corresponding *Q_X_* [15] statistic (testing for overall selection among the populations) was < 0.05/*n*, where n is the number of assessed GWASs. Traits with associated variants that passed this criterion are shown in Table 1. To account for possible artifacts arising from the ascertainment scheme for each GWAS, we also generated 1,000 samples in which we randomly switched the sign of the estimated effect size for all trait-associated SNPs. This serves to preserve the genetic architecture of each trait, while removing the effect of selection. We then computed a second P-value of the observed *Q_X_* (*P_rand_*) by comparing it to these samples (Table 1).

**Table 1:** Trait-associated variants with Bonferroni-corrected significant evidence of being under polygenic adaptation in the 1000 Genomes data, using the *Q_X_* statistic: *P* < 0.05/n where *n* is the number of GWAS tested, assuming a *χ*^2^ distribution. We also computed P-values from 1,000 samples in which we randomly switched the sign of effect size estimates, to simulate neutrality while preserving the genetic architecture of the traits (*P_rand_*). Additionally, we computed P-values from an empirical null distribution produced using 1,000 samples, each containing SNPs that were frequency-matched to the trait-associated SNPs, using their allele frequency in CEU, to account for each GWAS’s ascertainment scheme (*P_emp_*). Trait-associated variants for which *P_rand_* < 0.05/n are denoted with **, and trait-associated variants for which *P_rand_* < 0.05 are denoted with *.

| Graph | Trait | $P$ | $P_{rand}$ | $P_{emp}$ |
| --- | --- | --- | --- | --- |
| 7-leaf tree | height** | $3.516 * 10^{-13}$ | $< 0.001$ | $< 0.001$ |
| | educational attainment** | $3.404 * 10^{-6}$ | $< 0.001$ | $< 0.001$ |
| | unibrow* | $9.115 * 10^{-14}$ | 0.012 | $< 0.001$ |
|  | male-pattern baldness | 0.000679 | 0.177 | 0.007 |
| 5-leaf graph | height** | $2.355 * 10^{-11}$ | $< 0.001$ | $< 0.001$ |
| | educational attainment** | $1.379 * 10^{-7}$ | $< 0.001$ | $< 0.001$ |
|  | schizophrenia* | 0.000608 | 0.01 | 0.007 |
| | unibrow | $2.846 * 10^{-11}$ | 0.056 | $< 0.001$ |
|  | male-pattern baldness | 0.000562 | 0.213 | 0.003 |

We ran our MCMC on these trait-associated variants and obtained posterior distributions for the *α* parameters with the strongest evidence for selection, prioritizing branches as explained above (Figure S24). The P-values of the *Q_B_* statistic (obtained from a 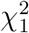 distribution) for each branch are shown in Table S2. We find strong evidence for selection on variants associated with height, educational attainment and self-reported unibrow, but little or no evidence for variants associated with male-pattern baldness or schizophrenia: even though these trait-associated variants are significant under the *Q_X_* and *Q_B_* frameworks, all their *α* parameters are approximately centered at 0. For height, we observe both selection for variants increasing height in the ancestral European branch and for variants decreasing height in the ancestral East Asian / Native American branch. However, this is only a consequence of the MCMC showing alternate strong support for selection in either one or the other branch at different points in the run, but only weak support for selection in both branches simultaneously (Figure S25), suggesting we are unable to discern which among these is the correct configuration.

To facilitate the visualization of posterior distributions for *α* parameters, we developed a new way to plot polygenic adaptation in a graph: a “poly-graph” (see Materials and Methods), which simultaneously depicts the structure of the studied graph and the marginal posterior mean of each *α* parameter, in the form of different colorings for each branch. We plotted poly-graphs for all traits that passed the significance criterion in the 7-leaf tree (Figure 4).

**Figure 4:**
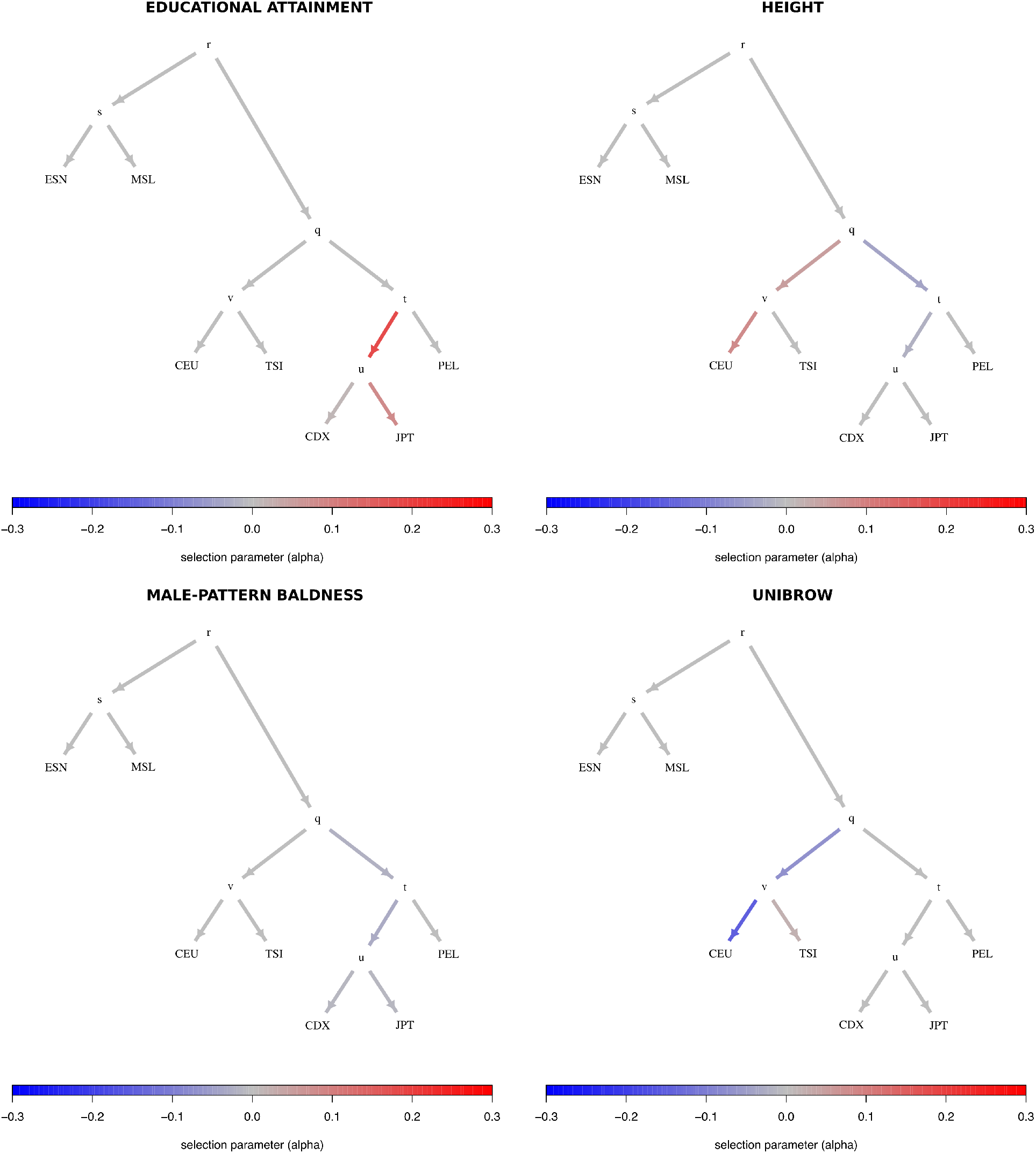
Poly-graphs for trait-associated variants that show significant evidence for polygenic adaptation in the 7-leaf tree built using 1000 Genomes allele frequency data. ESN = Nigerian Esan; MSL = Sierra Leone Mende; CEU = Northern Europeans from Utah; TSI = Southern Europeans from Tuscany; CDX = Dai Chinese; JPT = Japanese; PEL = Peruvians.

Given that PEL has European admixture, we also replaced PEL with an East Asian population (CHB) to verify that selection signals observed in the Eurasian branches were not dependent on poor modeling of PEL as a simple sister group to East Asians (Figure S26). Additionally, we tested an alternative set of panels, in which we kept PEL in the tree, but replaced the two European populations - CEU and TSI - by Finnish (FIN) and Iberians (IBS), and the two East Asian populations - JPT and CDX - by Han Chinese (CHB) and Southern Han (CHS) (Figure S27). The results from both alternative sets of panels are very similar to our original tree. We also find very similar results when using a Beta(2,2) prior for the root allele frequency in our MCMC (Figure S28), instead of the default Uniform[0,1] prior.

We then proceeded to explore a graph with an admixture event (Figure S23.B). This graph contained Yoruba (YRI), Colombians (CLM), CEU, CHB and PEL. We modeled CLM as resulting from an admixture event between CEU (76.55%) and PEL (23.45%), the latter of which is the panel with the highest amount of Native American ancestry in the 1000 Genomes Project [38]. Here, we recapitulated many of our previous findings from the 7-leaf tree (Table 1, Figure S29), like selection on variants associated with height and educational attainment. We list the P-values of the *Q_B_* statistic for each branch in Table S3. Poly-graphs of the 5-leaf admixture graph are shown in Figure 5.

**Figure 5:**
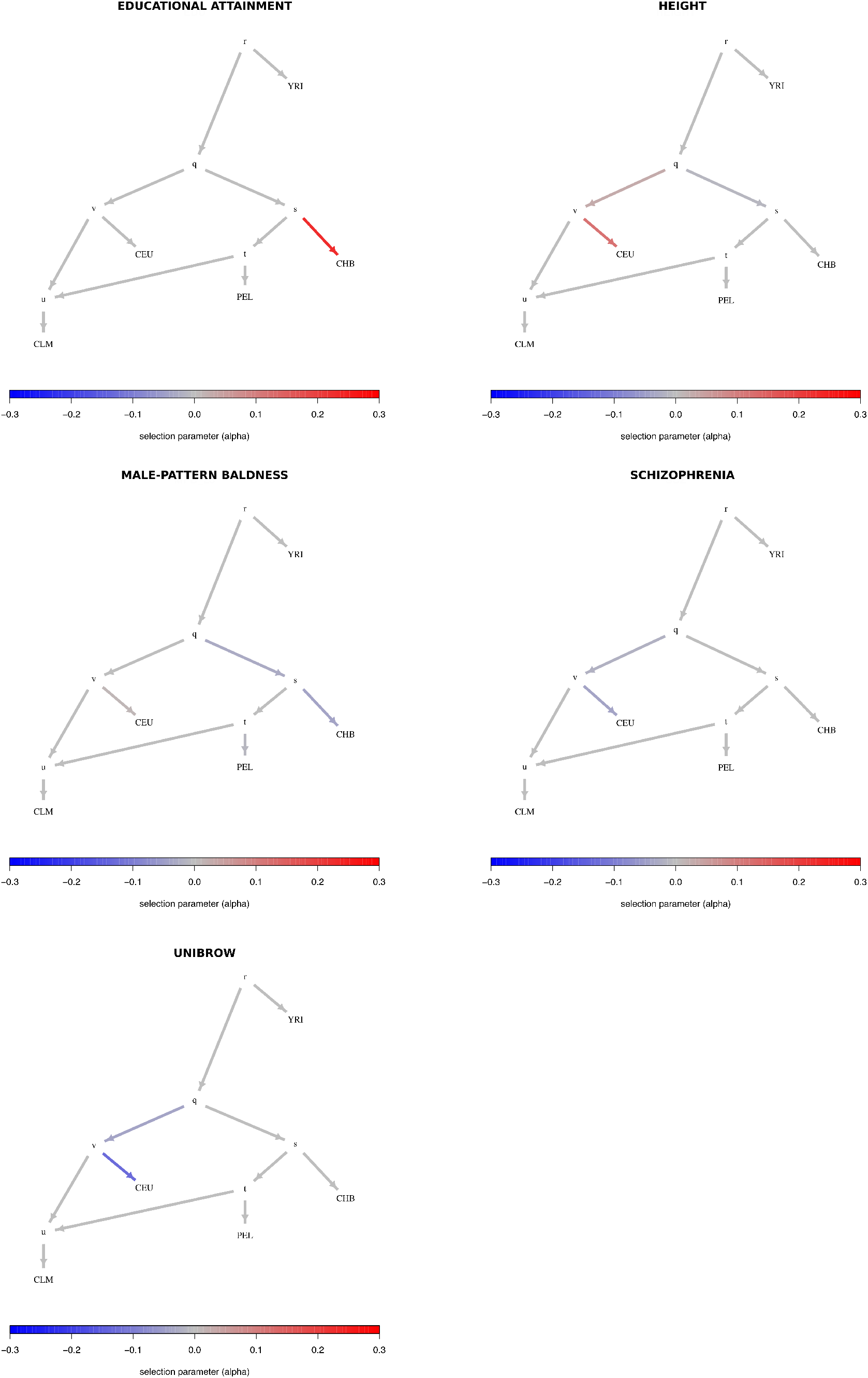
Poly-graphs for trait-associated variants that show significant evidence for polygenic adaptation in the 5-leaf admixture graph built using 1000 Genomes allele frequency data. CEU = Northern Europeans from Utah; TSI = Southern Europeans from Tuscany; PEL = Peruvians; CLM = Colombians; YRI = Yoruba; CHB = Han Chinese.

To make sure there were no artifacts due to GWAS ascertainment [15], we also generated an empirical null distribution produced using 1,000 samples, each containing SNPs that were frequency-matched to the trait-associated SNPs, using their allele frequency in CEU. We computed the *Q_X_* statistic for each of these samples, to obtain an empirical P-value (*P_emp_* in Table 1). We do not observe a value of *Q_X_* as high as the one observed in the real data, for either height, educational attainment or self-reported unibrow (Figure 6).

**Figure 6:**
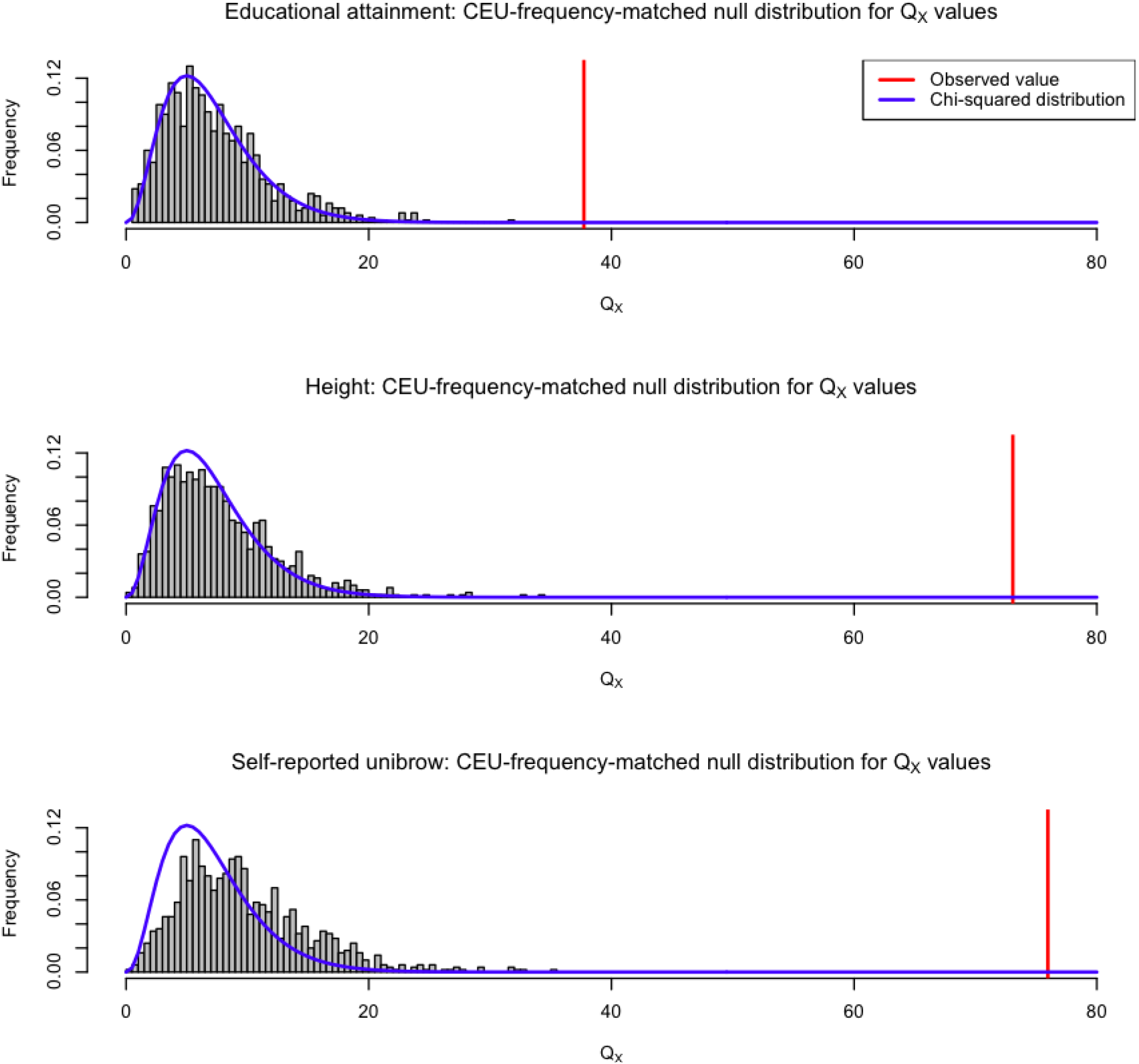
We generated an empirical null distribution by sampling SNPs from the genome that matched the CEU allele frequency of the SNPs associated with educational attainment, self-reported unibrow and height. We generated 1,000 samples this way, and computed the *Q_X_* statistic for each sample, using the population panels from Figure 4. The *Q_X_* value observed in the real data is depicted with a red line. We also plot the density of the corresponding *χ*^2^ distribution (blue line) for comparison.

To test how robust our results were to our modeling assumptions, we also performed a simpler two-tailed binomial sign test between every pair of 1000 Genomes panels. The assumption here is that - for every panel X and Y - we should observe roughly equal number of trait-increasing alleles at higher frequency in X than in Y as trait-decreasing alleles at higher frequency in X than in Y, under a model of neutrality with respect to the effect size sign [39]. This test only uses information about the sign of the effect estimates of each SNP, not their magnitudes, and does not use information about genome-wide drift parameters between each population. Thus, it is bound to have less power than the *Q_B_, Q_X_* or MCMC tests. The P-values for these pairwise binomial tests are shown in Tables S4 to S8 for all traits that were found to have significant evidence of selection using the Q*X* statistic. The top 10 most significant pairwise comparisons are shown in Tables S9 to S13. For ease of visualization, we also plotted, for each panel, the number of pairwise tests involving that panel that resulted in a P-value < 0.05 (Figures S30 to S34).

We were interested in verifying how sensitive different proportions of missing data (i.e. removal of SNPs) or erroneous effect size estimates would be to our three strongest signals of polygenic adaptation, on variants associated with height, educational attainment and unibrow. For this purpose, we focused on the comparison between CEU and CHB. First, we simulated different proportions of missing trait-associated SNPs - ranging from 5% to 95%, with step sizes of 5%. For each of 10,000 simulations under each missing data scenario, we assessed how often the polygenic score for unibrow and educational attainment in CHB was higher than the polygenic score for CEU, like we observe in the 1000 Genomes data. Height follows the opposite pattern (with CEU having a higher polygenic score than CHB), so in that case we assessed how often its polygenic score in CEU was higher than in CHB, across the 10,000 simulations for each scenario. Note that we built these scores using only the SNPs used in our selection tests. The results are in Figure S35. For example, we see that - even with 20% missing data - the polygenic scores for either of the 3 traits preserve the observed relationship of inequality between CEU and CHB almost 100% of the time. Finally, we simulated a situation in which some proportion of the signs of the effect size estimates were misassigned. We then assessed how often we could replicate the signal we see between CEU and CHB, but this time under different proportions of sign misassignment (Figure S36).

To understand how the signal of selection was distributed among our SNPs, we plotted the absolute value of the effect sizes of trait-associated SNPs for height, educational attainment and self-reported unibrow, as a function of the difference in frequency observed between CHB and CEU, polarized with respect to the trait-increasing allele in each SNP (Figure 7). We find that, in the case of self-reported unibrow, there are three variants of large effect with large frequency differences contributing to a higher polygenic score in CHB: rs3827760, rs16891982 and rs12916300. These SNPs are located in the genes *EDAR, SLC45A2* and *OCA2*/*HERC2*. These are genes involved in pigmentation and skin development, and all three have documented signatures of selective sweeps causing strong allele frequency differences between Europeans and East Asians [7, 3, 40, 2, 12]. After removing SNPs with large absolute effect size values (≥ 0.05), the P-value of the *Q_X_* statistic for these variants remains significant (*P* = 7.04 * 10^−5^). When looking at the other two sets (variants associated with height and educational attainment), the signal of selection is more uniformly distributed among the SNPs, with no strong outliers of large effect with large frequency differences (Figure 7).

**Figure 7:**
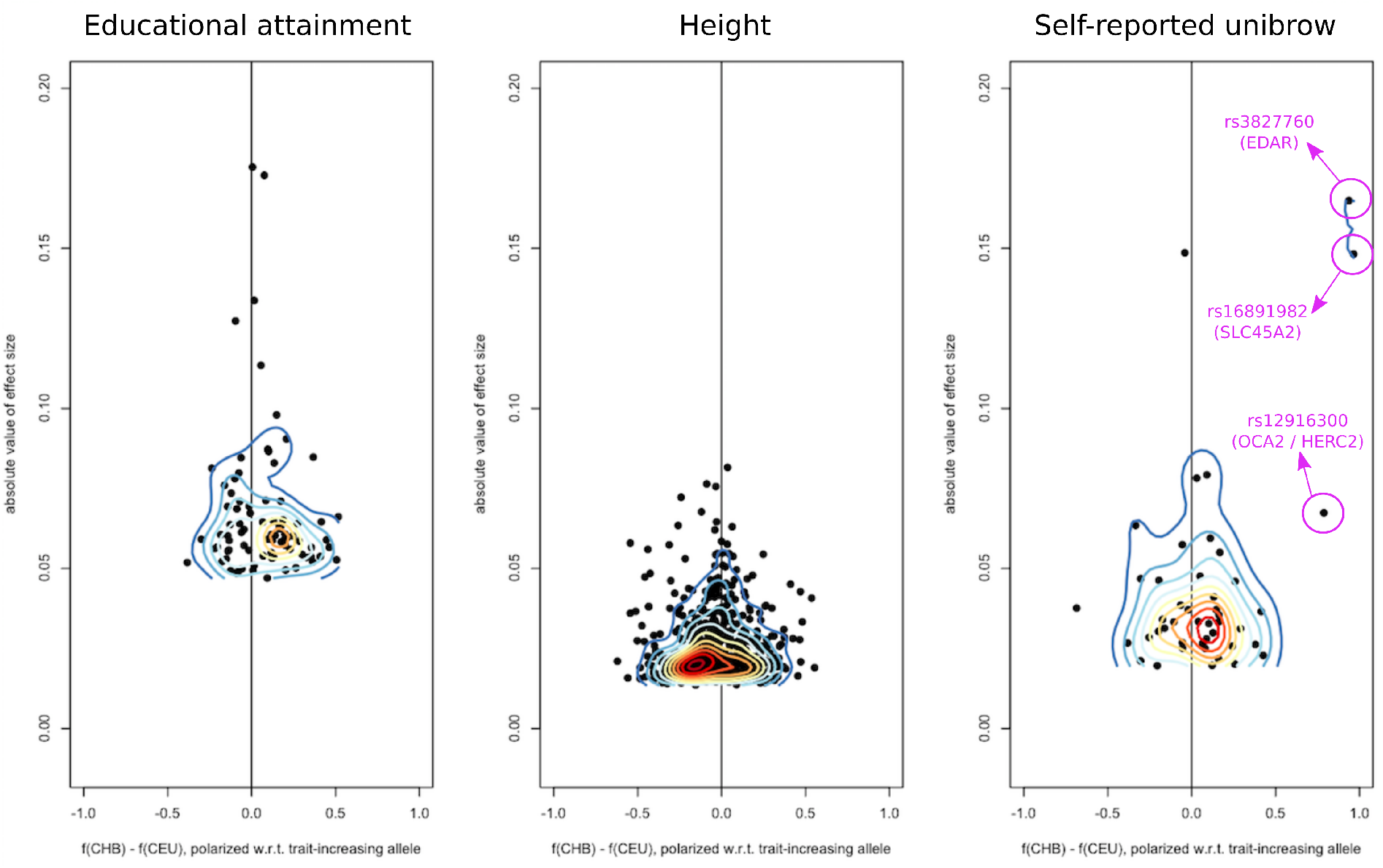
We plotted the absolute value of the effect sizes of trait-associated SNPs for height, educational attainment and self-reported unibrow, as a function of the difference in frequency observed between CHB and CEU. The panel frequencies are polarized with respect to the trait-increasing allele. We then overlaid a contour plot over each scatter-plot.

### Application to Lazaridis et al. (2014) data

We applied our method to a more broadly sampled SNP chip dataset containing present-day humans from 203 populations genotyped with the Human Origins array [19, 41]. This dataset was imputed using SHAPEIT [42] on the Michigan Imputation Server [43] with the 1000 Genomes Phase 3 data [38] as the reference panel (Bhérer et al. in prep.). We tested for polygenic adaptation in a 7-leaf admixture graph. This graph contains the panels Yoruba, Mandenka and Sardinian, along with the following 4 combinations of panels, which we built so as to have a large number of individuals per panel. The panel “Oceanian” contains the panels Papuan and Australian. The panel “EastAsian” contains the panels Cambodian, Mongola, Xibo, Daur, Hezhen, Oroqen, Naxi, Yi, Japanese, Han_NChina, Lahu, Miao, She, Han, Tujia and Dai. The panel “NativeAmerican” contains the panels Maya, Pima, Surui, Karitiana and Colombian. Finally, we modeled Europeans as a 2-way mixture of an ancestral component related to “NativeAmerican” and another component that split basally from the Eurasian tree and is a sister to Sardinians. This was the mixture fitted to Europeans by ref. [22], and provides a better fit to the data than modeling Europeans merely as a sister group to East Asians and Native Americans. Though we recognize that Europeans are better modeled as a 3- or 4-way mixture of ancestral components [41, 34, 44], it is hard to produce such a mixture without resorting to ancient DNA data (see Discussion). We tested 3 different versions of this graph, each containing three different sets of European populations (Figure S23.C) distinguished by how much “early European farmer” (EEF) ancestry they had (based on Figure 4 of ref. [41]). “EuropeA” (low EEF) contains the following panels: Estonian, Lithuanian, Scottish, Icelandic, Norwegian, Orcadian, Czech, English. “EuropeB” (medium EEF) contains Hungarian, Croatian, French, Basque, Spanish_North and French_South. Finally, “EuropeC” (high EEF) contains Bulgarian, Bergamo, Tuscan, Albanian, Greek and Spanish.

Trait-associated variants with significant evidence for polygenic adaptation are listed in Table 2 and the P-values of the *Q_B_* statistic for each branch are shown in Tables S14 to S16. With these data, we are able to recapitulate the adaptive increase in height-increasing variants in Europeans we had seen before, but only observe it in populations with medium or low EEF ancestry (Figures 8 and S37 to S41). This pattern is consistent with previous observations made using ancient DNA in Europeans [12]. We also recapitulate selection patterns on variants associated with other traits, like unibrow, educational attainment and male-pattern baldness, and observe evidence for polygenic adaptation in some additional trait-associated variants, like photic sneeze reflex (Table 2).

**Figure 8:**
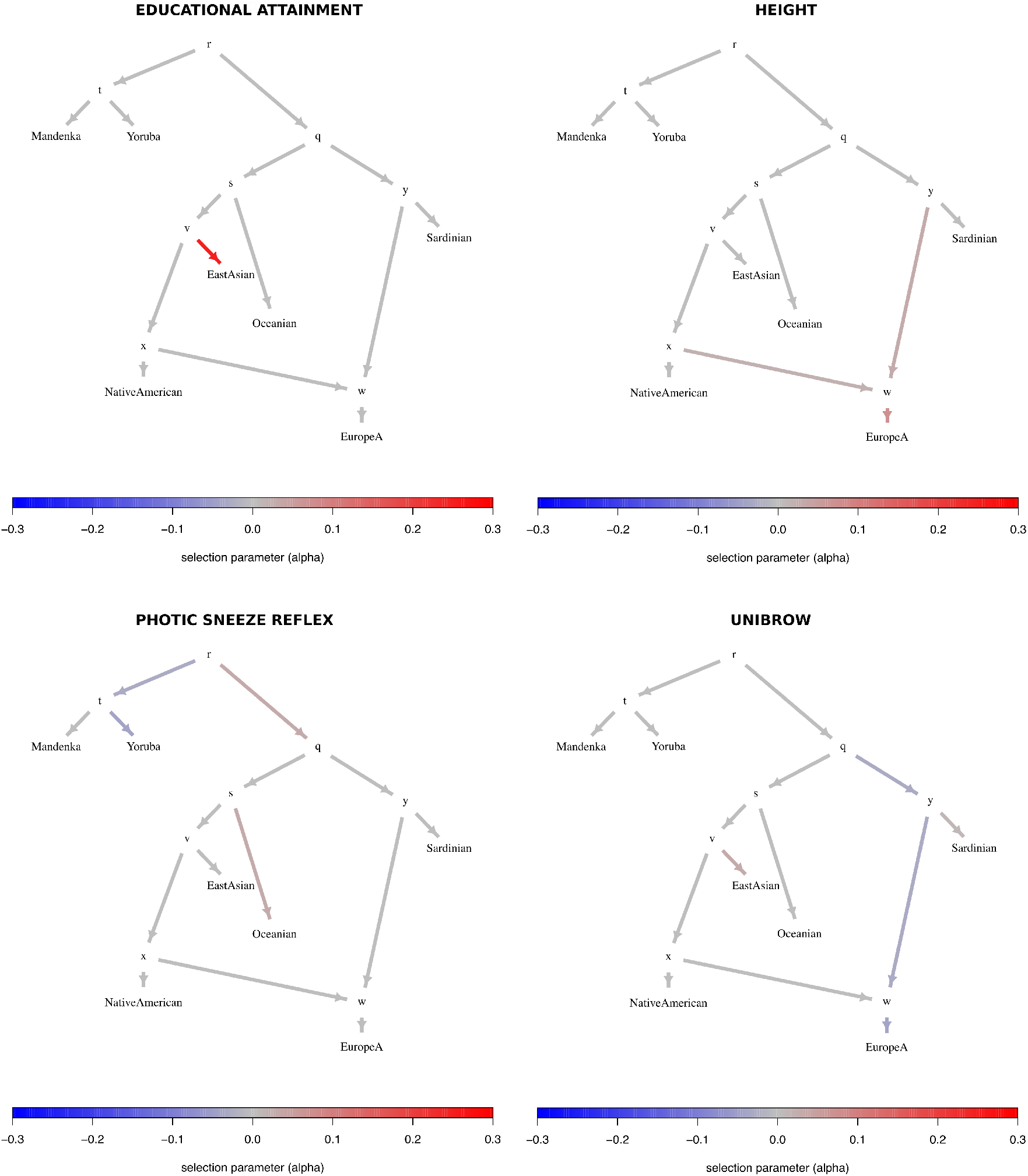
Poly-graphs for trait-associated variants that show significant evidence for polygenic adaptation in the 7-leaf admixture graph built using the Lazaridis et al. (2014) dataset and including the set of European populations with low EEF ancestry (“EuropeA”).

**Table 2:** Trait-associated variants with Bonferroni-corrected significant evidence of being under polygenic adaptation in the Lazaridis et al. (2014) dataset, using the *Q_X_* statistic: *P* < 0.05/*n* where *n* is the number of GWAS tested, assuming a *χ*^2^ distribution. We tested 3 different graphs with different sets of European panels, containing either low (EuropeA), medium (EuropeB) or high (EuropeC) EEF ancestry. We also computed P-values from 1,000 samples in which we randomly switched the sign of effect size estimates, to simulate neutrality while preserving the genetic architecture of the traits (*P_rand_*). Additionally, we computed P-values from an empirical null distribution produced using 1,000 samples, each containing SNPs that were frequency-matched to the trait-associated SNPs, using their allele frequency in Europeans, to account for each GWAS’s ascertainment scheme (*P_emp_*). Trait-associated variants for which *P_rand_* < 0.05/n are denoted with **, and trait-associated variants for which *P_rand_* < 0.05 are denoted with *.

| Graph | Trait | $P$ | $P_{rand}$ | $P_{emp}$ |
| --- | --- | --- | --- | --- |
| 7-leaf graph (w/EuropeA) | height** | $1.506 * 10^{-8}$ | $< 0.001$ | $< 0.001$ |
| | photic sneeze reflex** | 0.00087 | $< 0.001$ | 0.006 |
| | educational attainment** | $6.11 * 10^{-6}$ | 0.001 | 0.001 |
| | unibrow | $4.763 * 10^{-7}$ | 0.173 | $< 0.001$ |
| 7-leaf graph (w/EuropeB) | height** | $2.448 * 10^{-8}$ | $< 0.001$ | $< 0.001$ |
| | educational attainment** | $6.838 * 10^{-6}$ | $< 0.001$ | 0.001 |
| | age at voice drop** | 0.000953 | $< 0.001$ | 0.001 |
| | photic sneeze reflex** | 0.00086 | $< 0.001$ | 0.001 |
| | unibrow | $2.784 * 10^{-6}$ | 0.166 | $< 0.001$ |
| 7-leaf graph (w/EuropeC) | educational attainment** | $4.032 * 10^{-6}$ | $< 0.001$ | $< 0.001$ |
|  | photic sneeze reflex | 0.000653 | 0.004 | 0.004 |
| | unibrow | $2.957 * 10^{-6}$ | 0.199 | $< 0.001$ |

To check if there were systematic biases in ancestral/derived allele polarity relative to the direction of the effect size, we performed a two-tailed binomial test on each trait for which we found significant evidence of polygenic adaptation in the Lazaridis et al. or the 1000 Genomes dataset (Table S17). We find that only schizophrenia has a significant bias, showing an excess of derived alleles with negative effect sizes (P = 0.03511), though this is not significant after Bonferroni correction. We therefore caution that the evolution of these trait-associated variants may not be well-modeled by the multivariate Normal assumptions that we make to calculate the *Q_B_* statistic or when running the MCMC.

As before, to check the robustness of our results to our modeling assumptions, we show P-values for pairwise binomial sign tests involving each of the panels in the Lazaridis et al. (2014) dataset in Tables S18 to S22. The top 10 most significant pairwise comparisons are in Tables S23 to S26, with the exception of self-reported age at voice drop, in which all comparisons had P-values ≥ 0.25, due to the small number of associated SNPs. Figures S42 to S45 show, for each panel, the number of pairwise tests involving that panel that resulted in a P-value < 0.05.

### Replication using summary statistics from the UK Biobank

Given the potential contentiousness of our educational attainment signal, we aimed to determine whether the same global patterns were observed using summary statistics from a GWAS performed on an independent cohort. For this, we resorted to the GWAS set released by the Neale lab (http://www.nealelab.is/blog/2017/7/19/rapid-gwas-of-thousands-of-phenotypes-for-337000-samples-in-the-uk-biobank)and performed on the UK Biobank cohort [45]. Though we could not find “number of years of education” in the Neale lab GWAS, we found “college / university degree” and used this as a measure of educational attainment. We LD-partitioned the summary statistics in the same way as before, and then selected the SNP with the lowest P-value from each block. At *P* < 10^−9^, we obtain a similar number of SNPs (96) as we had for ref. [46] (86), and we find a 94% correlation between the population genetic scores made using the data from ref. [46] and the scores made using the UK Biobank (Figure S46), each standardized by their respective between-population standard deviations. At *P* < 10^−8^ and *P* < 10^−7^, the correlation is reduced to 56% and 86%, respectively. Using the 10^−9^ cutoff, we find a marginally significant overall *Q_X_* statistic when looking at the 7 populations from Figure 4 (*P* = 0.0398, *P_rand_* = 0.098, *P_emp_* = 0.076) and a significant *Q_B_* statistic in the ancestral East Asian branch (*P* = 0.013) and the terminal JPT branch (*P* = 0.002), but not in any of the other branches (*P* > 0.05). The *α* parameters of these two branches estimated from the PolyGraph MCMC are also positive (Figure S47), though their magnitude is not as large as the ones obtained from the summary statistics of ref. [46].

## Discussion

We have developed a method to infer polygenic adaptation on trait-associated variants in an admixture graph, so as to be able to pinpoint where in the history of a set of populations this type of selective processes took place. Our method requires GWAS data for a particular trait, allele frequency data for a set of populations, and a precomputed admixture graph that relates these populations with each other. Importantly, the method relies on the admixture graph as an accurate description of the ancestral genome-wide relationships among the populations under study. Potential users should be careful about correctly estimating branch lengths and ghost populations which are not included in the graph but may have substantial unaccounted ancestry contributions to the populations that are included. We used *MixMapper* [22] to infer the graph topology and branch lengths. Alternatively, one can also use other programs, like *qpGraph* [19] or *TreeMix* [20] to build graphs, though we caution that the estimated drift values of the branches in the output of these programs are scaled (in different ways) by the heterozygosity of ancestral nodes (see Supplementary Material of [22] for a way to properly obtain drift values from differences in allele frequencies between populations).

Running PolyGraph involves a two-step process, each of which is complementary to the other. The first step - the calculation of the *Q_B_* statistic - is fast and provides a preliminary way to assess which branch in a graph has significant evidence for polygenic adaptation. However, this statistic does not model the ancestral allele frequencies at each node of the graph. The second step - the MCMC - is slower, but provides posterior distributions for selection parameters under a more parameter-rich model of population history. In our pipeline, we use the first method as a filtering step, to avoid exploring selection parameters in the MCMC for those branches that have little evidence for selection, and encourage the MCMC to be sparse in its assignments of selection in the graph. We illustrate this point in Figure S48, where we show a side-by-side comparison of a poly-graph built using the posterior *α* estimates and a poly-graph built using *q_b_* - a signed version of the *Q_B_* statistic (see Materials and Methods).

In application to human populations, we detected signals of polygenic adaptation on sets of variants that have been identified to influence height, educational attainment, and self-reported unibrow. Selection on variants associated with height in Europeans has been previously reported elsewhere [15, 14, 17, 12] and our results are consistent with previous findings showing that height-increasing variants are at significantly and systematically higher frequencies in northern than in southern European populations. The signal for selection affecting variants associated with self-reported unibrow is also strong, but partly driven by a few variants of large effect with large frequency differences between populations, which have documented evidence for selective sweeps in genes involved in hair, skin and eye pigmentation, and skin development [3, 40, 7, 2, 12]. Additional trait-associated variants had inconsistent evidence across datasets and graph frameworks (like schizophrenia or male-pattern baldness) and/or are driven by differences in only a few SNPs of small effect (like age at voice drop), and so we do not discuss them.

We find preliminary evidence for polygenic adaptation in East Asian populations at variants that have been associated with educational attainment in European GWAS. This result is robust to the choice of population allele frequency data we used (1000 Genomes or Lazaridis et al. (2014) panels), to the choice of GWAS summary statistics (Figure S46), to GWAS ascertainment (Figure 6), and to our modeling assumptions, as we found a significant difference between East Asian and non-East-Asian populations even when performing a simple binomial sign test (Tables S4, S9, S19 and S23). However, we caution that this pends further verification via more GWAS on the same trait. Our modeling framework suggests that, if selection truly operated on these variants, it must have done so before or early in the process of divergence among East Asian populations - at least as far back as 5 thousand years ago [47, 48, 49, 50] - because the signal is common to different East Asian populations (Han Chinese, Dai Chinese, Japanese, Koreans, etc.). The signal seems only very weakly present in some Siberian populations - like the Even and Nganasan - and some Native American populations - like the Mixe and Pima, and not present at all in other Native American populations - like the Surui, Quechua and Karitiana. This is perhaps explained by the complex demographic make-up of Siberian and Native American populations, and their divergent history from East Asians [51, 52, 53].

Interpreting the educational attainment signal and the other signals we found requires awareness of a number of technical caveats, as well as several fundamental conceptual difficulties with the study of polygenic adaptation, some of which may ultimately prove intractable.

### What is the signal of polygenic adaptation?

Before discussing these difficulties, it is worth articulating exactly what a signal of polygenic adaptation consists of. Taking the height example as a case in point, the signal is that a set of genetic variants that have been identified as associated with increased height in a European GWAS are (as a class) at higher frequency in northern Europeans today than would be expected by genetic drift alone. Though this observation is consistent with the hypothesis that natural selection has operated on these variants, it does not necessarily imply that natural selection has operated directly on “height”, nor that observed height differences between northern Europeans and other populations are necessarily genetic and due to selection.

### Pleiotropy and phenotype definition

When looking at all our variant classes, we are necessarily limited by the traits that have been defined and studied by others, as we have grouped variants together based on these phenotypes. The use of previously established definitions makes it difficult to understand exactly why these variants may have been under selection in the past. This is perhaps best exemplified by the signal of polygenic adaptation for genetic variants associated with educational attainment.

Standardized schooling - and consequently, the concept of “educational attainment” - was only invented and implemented widely in the last few generations. It is obviously nonsensical to discuss its evolution over the past tens of thousands of years. Instead, it is likely that the set of variants for which we find evidence of selection was associated with some (unknown) phenotype(s) in the past. However, given that selection on these variants likely took place more than 5 thousand years ago, it may be difficult or impossible to identify what these were. A similar problem arises when thinking about the signal of polygenic adaptation on “unibrow”. This is a self-reported phenotype, and the genetic variants that have been identified may simply be associated with pigmentation (assuming people with certain hair and/or skin pigmentation phenotypes are more likely to notice they have hair between their eyes), or alternatively with some other (unmeasured) hair-related phenotype. It is also possible that direct sexual selection for absence or presence of unibrow as an attractive facial feature in certain cultures [54] may be the cause of this signal. Indeed, if a selective agent is cultural, but the culture has since changed, it may be impossible to determine what actually occurred. All these variants are also likely pleiotropic [55], which makes it even harder to determine which phenotypes were truly targeted by selection.

Perhaps, one could try to find the phenotypic gradients along which selection most likely operated [56] by modeling the evolution of trait-associated SNPs for multiple phenotypes together. However, it is also possible that genetic correlations among traits in the present are not good proxies for genetic correlations in the past.

### Relationship between polygenic scores and population mean phenotypes

Another fundamental limitation in interpreting all studies of polygenic adaptation (including this one) is that the connection between the distribution of allele frequencies today and any historical or geographic trends in phenotypes remains questionable. Indeed, though we have motivated this method as a way to identify adaptive shifts in the mean of a polygenic trait, it is a simple fact that massive changes in the mean values of many of the traits we consider have occurred by purely non-genetic environmental processes. For example, the mean height of men in the Netherlands increased from around 166 cm in the mid-1800s to currently over 180 cm [57], bringing the population from around the middle of the pack among European countries to the tallest one in the world. This likely occurred for environmental reasons, such as improvements in diet and health care [58, 59]. Likewise, the average educational attainment in Iceland and North America has increased dramatically over the past century, despite a slight estimated *decrease* in the frequencies of genetic variants associated with an increased value of the phenotype [60, 61, 62]. The somewhat paradoxical conclusion is that actual phenotypes can and do change across populations in directions that are uncorrelated to natural selection (which may in fact be a minor contributor to any such differences). It would be an understatement to say this poses challenges for the interpretation of the current study and others like it.

In fact, the trait-associated variants that we have used only explain a fraction of the narrow-sense heritability of their respective traits, even in the populations in which the association studies were performed. As we have only looked at variants that have high probability of association with a trait, this fraction is small in most cases. For example, the heritability for ‘educational attainment’ is estimated to be around 40%, and educational attainment itself is strongly determined by environmental factors [63]. The SNPs we used in this study (themselves a subset of all SNPs tested in the original GWAS [46]) explain only 1.05% of the total variance for this particular trait. All of the aforementioned traits are likely affected by a myriad of environmental and social variables, which might contribute to determine their ultimate expression in each human individual.

### Additional caveats

Beyond the above conceptual difficulties, there are a number of additional caveats with our approach to keep in mind. First, the effect sizes we have used derive from GWAS performed primarily on individuals of European ancestry. Thus, our tests can only detect if variants that have been found to be associated with a trait in European GWAS are significantly higher or lower in a particular (European or non-European) population, relative to what they should be under a pure drift model. This does not necessarily imply that populations for which we find evidence for selection have higher or lower average genetic values of such a trait than other populations. In fact, there is evidence to suggest that loci ascertained in European GWAS do not serve to make good predictors for traits in populations that are distantly related to Europeans [64]. One reason for this is that many or all of the traits we are studying are likely to be influenced in non-European populations by different variants from the ones that have been discovered in European GWAS. SNPs that may be strongly associated with a trait in a particular non-European population (like an African or East Asian panel) may have not reached genome-wide significance in a European GWAS, where those SNPs may not strongly affect the trait or may be at low frequencies. It is thus possible that there are variants associated with traits like educational attainment that occur at high frequencies in East Asians, but that are missing from our analysis, or that the effect sizes in trait-associated SNPs are different in non-European populations, in such a way that the average genetic values between these populations and Europeans are not significantly different. We also do not model dominance, epistasis or gene-by-environment interactions between our trait-associated variants and the diverse environments that human populations occupy, and any of these factors may further obscure the relationship between the patterns we observe and the actual underlying genetic contribution to phenotypes in these populations.

Second, we have assumed that all of the GWAS that we have used have properly accounted for population structure. If some of the trait-associated SNPs are in fact false positives caused by uncorrected structure, this could generate a false signal of polygenic adaptation. A future direction could be the incorporation of effect sizes that have been corrected for ancestry or population stratification [17, 65] and also effect sizes from GWAS performed on other populations [66, 67, 68], in order to assess the robustness of our empirical results across variants discovered in studies involving participants of different ancestries.

Third, we made the assumption that the admixture graph for the populations that we use as input is correct. If there are additional unmodeled aspects of the history of the populations, this could induce incorrect inference about the branch on which natural selection has occurred. We also recommend that the individuals in the population panels used as leaves in the graphs have roughly similar amounts of admixture. In other words, the method works best when admixture in the population was ancient enough for the admixed ancestry to have spread uniformly among members of the admixed panel. Otherwise, an admixture graph may not be the most appropriate way to model their evolution.

Finally, we have made an explicit assumption that our model should be sparse; i.e. that polygenic adaptation is rare. If in reality adaptation is common, the PolyGraph approach will necessarily only identify selection on a small number of branches.

### Future directions

A natural extension to the analyses we performed here would be to look at admixture graphs that include extinct populations or species, using ancient DNA [69]. For example, present-day Europeans are known to have resulted from admixture processes involving at least 4 ancestral populations [41, 44], and so modeling them as a sister group to East Asians or as a 2-way mixture between a Native American-related component and a basal Eurasian component may be overly simplistic. Incorporating ancient DNA would not require any additional theoretical work, as ancient populations can be naturally included as leaves in an admixture graph [41, 12]. Care should be taken, however, in making sure that the quality of the ancient DNA data at trait-associated SNPs is accounted for while inferring the number of ancestral and derived alleles, and that there is a sufficient number of ancient individuals per population to detect polygenic adaptation. One could envision either performing pseudo-haploid sampling [19, 34, 12] or using allele frequency estimators obtained from genotype likelihoods [70, 71], while accounting for errors characteristic of ancient DNA [72, 73]. When working with SNP capture data [19, 34], it may be necessary to perform imputation at the GWAS SNPs, if these were not originally covered in the SNP capture array. We aim to tackle these issues in a future study.

One concern when analyzing admixture graphs is identifiability. As we mentioned before, there are multiple configurations of the *α* parameters that may lead to almost identical likelihoods. The use of the spike-and-slab prior and the *Q_B_* filtering step serve to ameliorate this problem, assuming selection was sparse and only affected a few branches. An avenue of research could involve testing other types of models or constraints that may serve to better compare among different selection configurations, perhaps without having to reduce the space of possible candidate branches a priori, for example using reversible-jump MCMC for model selection [74].

In the future, it may be worth incorporating stabilizing selection into this method [55], or exploring tests of polygenic adaptation in the context of other types of demographic frameworks, like isolation-by-distance [75] or population structure [76] models. For example, one could envision settings in which trait-associated variants would be best modeled as expanding or contracting over a geographically extended area over time, in a way that is not explainable by genetic drift alone.

Lastly, we note that despite some clear methodological and conceptual differences, our method bears a close relationship to a number of methods for inferring changes in the rate of phenotypic evolution on species phylogenies over macroevolutionary timescales. Our use of the Normal model of drift as an approximation to the Wright-Fisher diffusion is closely analogous to the use of Brownian motion models in some phylogenetic methods [77, 78, 79, 80, 81]. It may also be worth exploring the relationship between Ornstein-Uhlenbeck models for phenotypic evolution on phylogenies [82, 83] and the aforementioned hypothetical extension of our method to include stabilizing selection, as the two processes are closely related [84, 55].

## Materials and Methods

### MCMC implementation

For the MCMC transition probabilities of the *α* parameters, we use a Normal distribution with constant variance. For the transition probabilities of the ancestral allele frequencies, we use a truncated Normal distribution with point masses at 0 and 1, with variance equal to a constant (input by the user) times *f_X_* (1 – *f_X_*) where *f_X_* is the frequency of an allele in its current state. This allows for the proposed transitions to be larger for SNPs at medium frequencies and smaller for SNPs at high or low frequencies. We apply a Hastings correction in the acceptance ratio to account for this asymmetric proposal distribution.

For all applications above, we run our MCMC sampler for 1 million steps with a burn-in period of 10,000, and obtain posterior samples every 1,000 steps. The variance of the transition probabilities of the ancestral nodes and the *α* parameters were chosen so that the acceptance rate was close to 23% for each set of parameters. For the spike-and-slab prior, we set *τ* to be equal to 0.1, and *ζ* to be equal to 25. The lower and upper boundaries of the uniform hyperprior for *κ* were set to be 60% and 80%, respectively, unless otherwise stated. We note that all of these parameters can be adjusted by the user as needed. To verify that the MCMC chain was mixing well, we also built auto-correlation plots of the alpha parameters (Figure S49 for the chain corresponding to simulation 1 of Figure S5).

### Derivation of *Q_B_* statistic

Let 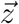 be the mean-centered vector of genetic values, **F** be the among-population genetic covariance matrix, and *V_A_* be the additive genetic variance of the ancestral population for a given character. We compute *V_A_* by taking the ancestral frequency for each SNP *i* to be the mean sample frequency over all populations 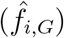:

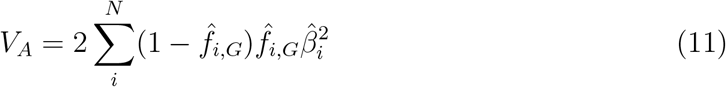

where 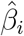 is the effect size estimate for the trait at SNP i. Following ref. [85, 15], if we are willing to assume that

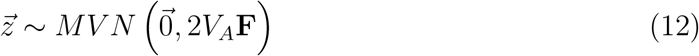

then by the definition of the multivariate normal distribution, 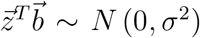 *for any choice* of 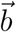 and

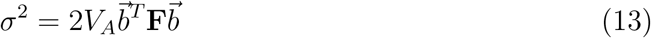

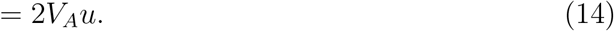

where we have set 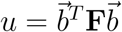 for notational convenience. It follows that

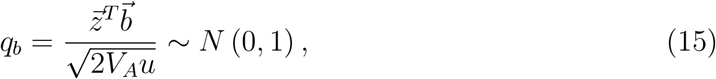

and this holds for all choices of 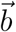. Therefore, the square of *q_b_* has a 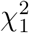 distribution under the null. Importantly, one is free to choose 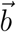 such that it represents a branch (j) in an admixture graph.

If j is a branch in an admixture graph, and we choose to scale 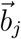 such that it has unit length:

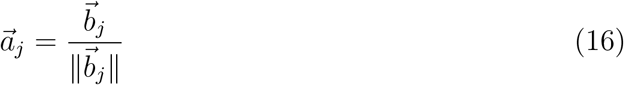

then

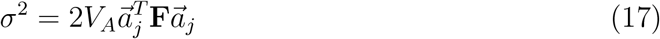

has an interpretation as the amount of among population additive genetic variance which we would expect to come about *because of* drift which occurs down branch j. In turn, 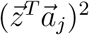 is the actual amount of variance observed along the axis consistent with that branch. The ratio of these two quantities is our statistic *Q_B_*(*j*) (with distribution 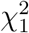) and it is therefore the appropriate test statistic to ask whether there is evidence to reject neutrality along a certain branch. Note that, by design, branches with the exact same child nodes have equal *Q_B_* statistics, as do branches at the root of the graph.

### Choosing a cutoff for *Q_B_*

We aimed to find a cutoff for *Q_B_* that would serve to minimize the number of candidate branches to be explored in the MCMC while at the same time trying to ensure that the true selected branches are included among these candidates. One choice would be to select the cutoff of a 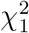 distribution that would correspond to a P-value of 0.05/*κ*, where *κ* is the number of branches tested. We find, however, that a constant value for this cutoff is not the most desirable choice as a way to prioritize branches for exploring the strength of selection in each of the branches in the MCMC, as graphs of different sizes (i.e. amounts of drift) result in quite different senstivity values, as well as number of candidate branches included, when *α* = 0.1 (Panels B and C in Figures S50 and S51 for two cases where each branch has drift length = 0.02 and Figures S52 and S53 for two cases where each branch has drift length = 0.05). We observe the same issues when simulating under *α* = 0.2 (Figures S54 to S57).

A more stable strategy across graphs of different sizes that also works better at minimizing the number of candidates is to choose the cutoff to be a fraction of the largest *Q_B_* statistic among all branches in the graph (Panels D and E in Figures S50 to S53). For all analyses below, we chose this to be 1/3 of the maximum *Q_B_* statistic. We note that this is a less conservative strategy than the fixed 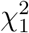 cutoff. However, it is important to remember that we do not aim to formally test for selection here, but merely obtain a set of likely candidate branches for the MCMC to explore downstream, some of which may end up producing a posterior mean estimate of *α* that is consistent with neutrality (i.e. *α* = 0).

### Simulations

For each simulated SNP, we sampled the root allele frequency (*f_R_*) from a Beta(2,2) distribution, to emulate the fact that, in the real data, variants in the leaf populations tend to be further away from the boundaries of fixation and extinction than under a uniform distribution. We evolved the SNPs throughout an admixture graph forwards in time using Wright-Fisher binomial sampling, and used binomial distributions to sample panel allele frequencies from the leaf populations. We set a constant effective population size *N_e_* of 10,000 and adjusted the number of generations in each branch, depending on how much drift we specified in each scenario. We used equation 2 from the Supplementary Note of ref. [14] to simulate selection. We set a constant time during which selection operates (*T_se1_*) to be equal to 100 generations, and adjusted the selection coefficient (s) to obtain the selection parameter specified in each scenario (*α* = *s* * *T_se1_*). If a branch had a larger number of generations than *T_sel_*, the selective phase was simulated to occur during the ancestral-most generations, followed by a neutrality period until reaching the end of the branch. The effect sizes of the SNPs were drawn from a Normal distribution with mean 0 and we simulated polygenic adaptation in a particular branch using only the sign of the effect size of each SNP. We also simulated an additional 10,000 SNPs that evolved neutrally under the same demography, so as to estimate the neutral population covariance matrix.

### Building admixture graphs

We used *MixMapper* (v1.02) [22] to build best-fitting scaffold trees, and then place putatively admixed populations as mixtures originating from different branches in these trees. We first pruned the 1000 Genomes data and the Human Origins imputed SNP data by sampling 1 out of every 100 SNPs before feeding it as input into *MixMapper*. To account for any residual linkage disequilibrium, we also performed 100 bootstrap replicates of the computed statistics, computed over 500 blocks along the genome.

The 7-leaf tree obtained using the 1000 Genomes data was the most additive fitted tree that contained ESN, MSL, CEU, TSI, CDX, JPT and PEL. To build the 5-leaf graph, we first fit the most additive tree containing CEU, CHB, YRI and PEL, and then attempted to fit CLM as a putative mixture of branches in the tree. The best-fitting combination was a mixture of the terminal branch leading to PEL and the terminal branch leading to CEU (residual norm = 1.91e-07). We also verified that the topologies we found in MixMapper were the same topologies as the ones inferred by *TreeMix* [20] (Figure S58). We note that adding one migration event to the 7-leaf tree makes PEL an admixed population, which is why we also verified our results were robust to replacing PEL by an East Asian population in the tree.

For building the 7-leaf graph using the Human Origins data, we again used *MixMapper* to fit the most additive tree, this time containing the following populations: Mandenka, Yoruba, Oceanian, East Asian, Native American and Sardinian. Then, we attempted to fit Europeans onto this tree as a putative mixture of branches in the tree, in an analogous way in which European populations were fitted in ref. [22]. The best-fitting combination was a mixture of the branch leading to Sardinians and the branch leading to Native Americans (residual norm = 6.5e-07, 1.08e-06 and 1.97e-06, when fitting EuropeA, EuropeB and EuropeC, respectively).

### Visualizing poly-graphs

In a poly-graph, the vertical component of a non-admixing branch is proportional to the amount of genetic drift that it experienced (calculated via *MixMapper*). The position of admixed nodes is determined based on the drift value of one randomly chosen parent branch. The colors indicate the marginal posterior mean estimate of the selection parameter for variants associated with the corresponding trait (with red indicating an increase in trait-increasing variant frequency, and blue indicating a decrease in trait-increasing variant frequency). We impose a minimum branch height (= 0.075) for clarity, as otherwise the selection parameters of some branches with very short drift lengths are impossible to visualize.

## Acknowledgments

We thank Claude Bhérer, Guy Sella, Molly Przeworski, Graham Coop, Joshua Schraiber, Simon Myers, Benjamin Peter and members of the Sella and Przeworski labs for helpful advice and discussions. We also thank an anonymous reviewer for valuable feedback, and Thomas Mailund for assistance with running the *admixturegraph* R package as well as helpful comments on the manuscript. Finally, we thank Mark Lipson for help with running *MixMapper*. This work was supported by NIGMS grant 1R01GM121372-01 to JKP.

---

**Table S1:**
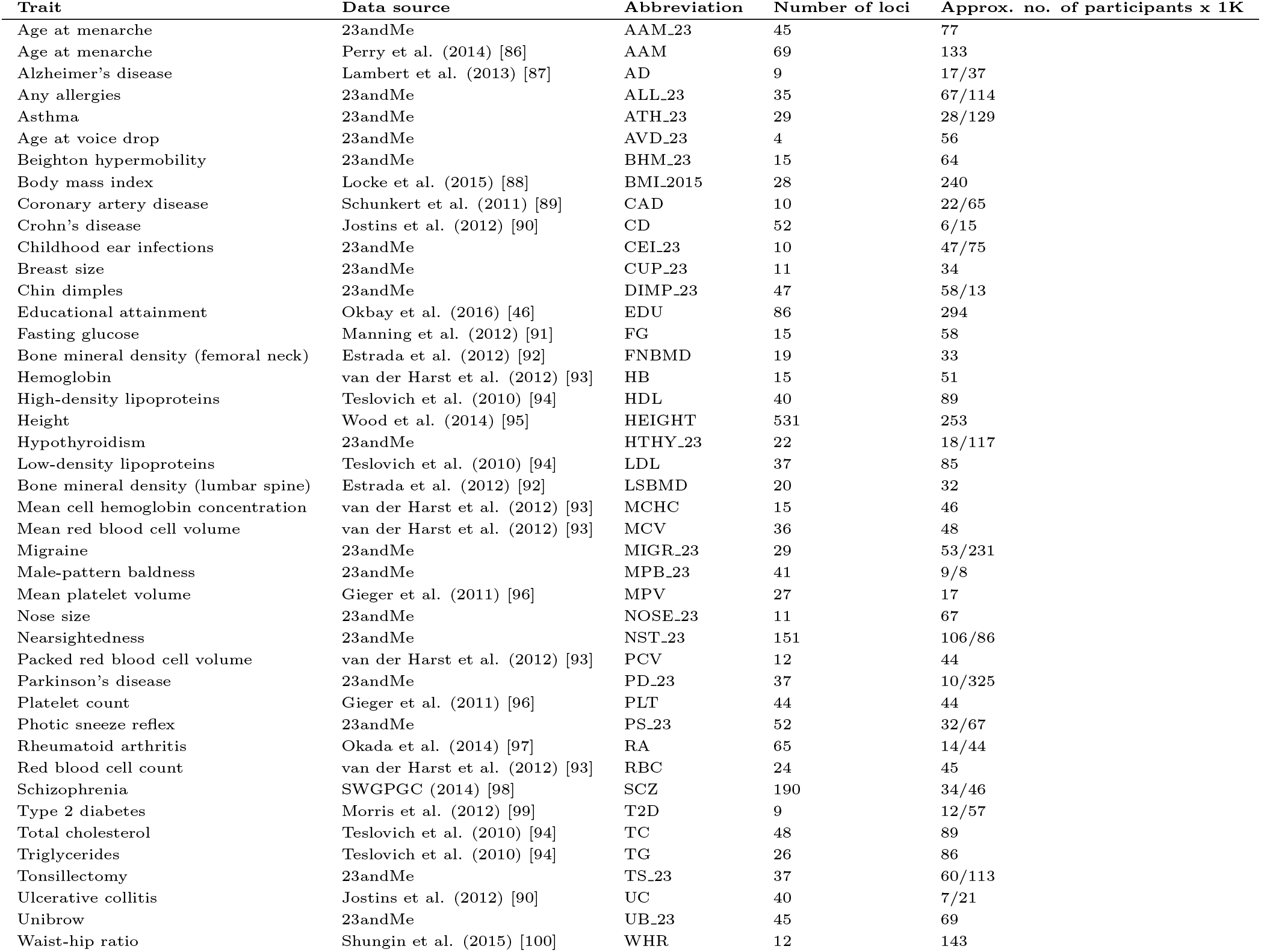
List of GWAS assembled by [35] and used to test for polygenic adaptation in this study. Under “Number of loci”, we list the number of autosomal SNPs that are significant for the trait in question, that overlap with the 1000 Genomes dataset and that have a confidently determined ancestral allele. Under “Approx. no. of participants x 1K”, we list two numbers corresponding to the approximate number of cases and controls (in thousands), if available. Otherwise, we list the approximate total number of participants, in thousands. SWGPGC = Schizophrenia Working Group of the Psychiatric Genomics Consortium.

**Table S2:** P-values from the *Q_B_* statistics of the 7-leaf population tree built using the 1000 Genomes data, for trait-associated variants with significant overall evidence of selection (P-value of *Q_X_* < 0.05/no. of GWAS tested).

| Branch (child-parent) | Educational attainment | Height | Male-pattern baldness | Unibrow |
| --- | --- | --- | --- | --- |
| q-r | 0.245 | 0.819 | 0.0247 | 0.24 |
| t-q | 0.0129 | 0.000191 | 5.87e-5 | 0.001 |
| u-t | 7.63e-06 | 0.000304 | 0.000155 | 0.00698 |
| s-r | 0.245 | 0.819 | 0.0247 | 0.24 |
| v-q | 0.147 | 5.14e-6 | 0.0777 | 6.08e-10 |
| MSL-s | 0.231 | 0.851 | 0.0216 | 0.301 |
| ESN-s | 0.292 | 0.799 | 0.0407 | 0.215 |
| TSI-v | 0.217 | 0.00284 | 0.217 | 7.51e-7 |
| CEU-v | 0.114 | 4.54e-9 | 0.0284 | 1.48e-12 |
| CDX-u | 0.000227 | 0.00105 | 0.000111 | 0.00177 |
| JPT-u | 1.59e-6 | 0.000362 | 0.000973 | 0.0482 |
| PEL-t | 0.0127 | 0.0632 | 0.0282 | 0.0118 |

**Table S3:** P-values from the *Q_B_* statistics of the 5-leaf population graph built using the 1000 Genomes data, for trait-associated variants with significant overall evidence of selection (P-value of *Q_X_* < 0.05/no. of GWAS tested).

| Branch (child-parent) | Educational attainment | Height | Male-pattern baldness | Schizophrenia | Unibrow |
| --- | --- | --- | --- | --- | --- |
| YRI-r | 0.629 | 0.752 | 0.0554 | 0.0747 | 0.282 |
| q-r | 0.629 | 0.752 | 0.0554 | 0.0747 | 0.282 |
| s-q | 0.0611 | 0.000286 | 2.82e-5 | 0.912 | 0.000106 |
| v-q | 0.107 | 7.11e-7 | 0.0178 | 0.013 | 1.717e-10 |
| CEU-v | 0.146 | 5.62e-10 | 0.0099 | 0.00154 | 1.183e-12 |
| CHB-s | 2.87e-7 | 0.00142 | 0.000127 | 0.447 | 0.0027 |
| t-s | 0.00516 | 0.0369 | 0.0248 | 0.309 | 0.00722 |
| u-t | 0.128 | 0.33 | 0.197 | 0.717 | 0.00169 |
| u-v | 0.128 | 0.33 | 0.197 | 0.717 | 0.00169 |
| PEL-t | 0.00719 | 0.0216 | 0.0123 | 0.315 | 0.00129 |
| CLM-u | 0.128 | 0.33 | 0.197 | 0.717 | 0.00169 |

Table S4: Results from pairwise two-tailed binomial sign tests using the 1000 Genomes data, for variants associated with educational attainment. Each cell contains the number of SNPs in which the two panels have different frequencies, the number of SNPs in which the column panel has a higher frequency of the trait-increasing allele than the row panel and the P-value from the binomial test.

Table S5: Results from pairwise two-tailed binomial sign tests using the 1000 Genomes data, for variants associated with height. Each cell contains the number of SNPs in which the two panels have different frequencies, the number of SNPs in which the column panel has a higher frequency of the trait-increasing allele than the row panel and the P-value from the binomial test.

Table S6: Results from pairwise two-tailed binomial sign tests using the 1000 Genomes data, for variants associated with self-reported male-pattern baldness. Each cell contains the number of SNPs in which the two panels have different frequencies, the number of SNPs in which the column panel has a higher frequency of the trait-increasing allele than the row panel and the P-value from the binomial test.

Table S7: Results from pairwise two-tailed binomial sign tests using the 1000 Genomes data, for variants associated with schizophrenia. Each cell contains the number of SNPs in which the two panels have different frequencies, the number of SNPs in which the column panel has a higher frequency of the trait-increasing allele than the row panel and the P-value from the binomial test.

Table S8: Results from pairwise two-tailed binomial sign tests using the 1000 Genomes data, for variants associated with self-reported unibrow. Each cell contains the number of SNPs in which the two panels have different frequencies, the number of SNPs in which the column panel has a higher frequency of the trait-increasing allele than the row panel and the P-value from the binomial test.

**Table S9:** Top 10 most significant comparisons from pairwise two-tailed binomial sign tests using the 1000 Genomes data, for variants associated with educational attainment. *N* = number of SNPs in which the two panels have different frequencies. *P*1 > *P*2 = number of SNPs in which Pop1 has a higher frequency of the trait-increasing allele than Pop2. Two-tailed Pval = P-value from the binomial test.

| Pop1 | Pop2 | N | P1 > P2 | Two-tailed Pval |
| --- | --- | --- | --- | --- |
| JPT | AMR | 86 | 63 | 1.88E-05 |
| CHB | AMR | 86 | 61 | 0.000130379 |
| KHV | PEL | 84 | 58 | 0.000627947 |
| JPT | GIH | 86 | 59 | 0.00073171 |
| JPT | TSI | 86 | 59 | 0.00073171 |
| PUR | CHB | 86 | 27 | 0.00073171 |
| PUR | JPT | 86 | 27 | 0.00073171 |
| MXL | CHB | 85 | 27 | 0.001016221 |
| MXL | CHS | 85 | 27 | 0.001016221 |
| JPT | MXL | 85 | 58 | 0.001016221 |

**Table S10:** Top 10 most significant comparisons from pairwise two-tailed binomial sign tests using the 1000 Genomes data, for variants associated with height. *N* = number of SNPs in which the two panels have different frequencies. *P*1 > *P*2 = number of SNPs in which Pop1 has a higher frequency of the trait-increasing allele than Pop2. Two-tailed Pval = P-value from the binomial test.

| Pop1 | Pop2 | N | P1 > P2 | Two-tailed Pval |
| --- | --- | --- | --- | --- |
| CLM | CEU | 531 | 194 | 5.68E-10 |
| AMR | CEU | 531 | 198 | 5.04E-09 |
| EUR | AMR | 531 | 329 | 3.95E-08 |
| CLM | GBR | 531 | 203 | 6.47E-08 |
| CEU | PEL | 531 | 327 | 1.05E-07 |
| AMR | GBR | 531 | 204 | 1.05E-07 |
| PUR | CEU | 531 | 205 | 1.70E-07 |
| EUR | CLM | 531 | 326 | 1.70E-07 |
| CEU | TSI | 531 | 325 | 2.73E-07 |
| CEU | IBS | 531 | 324 | 4.34E-07 |

**Table S11:** Top 10 most significant comparisons from pairwise two-tailed binomial sign tests using the 1000 Genomes data, for variants associated with self-reported male-pattern baldness. *N* = number of SNPs in which the two panels have different frequencies. *P*1 > *P*2 = number of SNPs in which Pop1 has a higher frequency of the trait-increasing allele than Pop2. Two-tailed Pval = P-value from the binomial test.

| Pop1 | Pop2 | N | P1 > P2 | Two-tailed Pval |
| --- | --- | --- | --- | --- |
| AFR | KHV | 40 | 29 | 0.006426576 |
| CDX | ITU | 40 | 11 | 0.006426576 |
| ESN | KHV | 38 | 27 | 0.013852965 |
| PJL | SAS | 40 | 28 | 0.016589003 |
| AFR | CHS | 40 | 28 | 0.016589003 |
| KHV | ASW | 39 | 12 | 0.023702702 |
| CHS | STU | 39 | 12 | 0.023702702 |
| CDX | STU | 39 | 12 | 0.023702702 |
| CHS | EAS | 36 | 11 | 0.02881672 |
| KHV | LWK | 38 | 12 | 0.03355244 |

**Table S12:** Top 10 most significant comparisons from pairwise two-tailed binomial sign tests using the 1000 Genomes data, for variants associated with schizophrenia. *N* = number of SNPs in which the two panels have different frequencies. *P*1 > *P*2 = number of SNPs in which Pop1 has a higher frequency of the trait-increasing allele than Pop2. Two-tailed Pval = P-value from the binomial test.

| Pop1 | Pop2 | N | P1 > P2 | Two-tailed Pval |
| --- | --- | --- | --- | --- |
| PJL | FIN | 189 | 118 | 0.000775545 |
| ITU | GBR | 189 | 116 | 0.002165972 |
| YRI | LWK | 173 | 107 | 0.002262171 |
| FIN | MSL | 190 | 74 | 0.002837056 |
| ESN | GBR | 190 | 116 | 0.002837056 |
| ASW | GWD | 185 | 72 | 0.003165809 |
| FIN | STU | 189 | 74 | 0.00350814 |
| ESN | ASW | 184 | 112 | 0.003917461 |
| AFR | GBR | 190 | 115 | 0.004537135 |
| AFR | FIN | 190 | 115 | 0.004537135 |

**Table S13:** Top 10 most significant comparisons from pairwise two-tailed binomial sign tests using the 1000 Genomes data, for variants associated with self-reported unibrow. *N* = number of SNPs in which the two panels have different frequencies. *P*1 > *P*2 = number of SNPs in which Pop1 has a higher frequency of the trait-increasing allele than Pop2. Two-tailed Pval = P-value from the binomial test.

| Pop1 | Pop2 | N | P1 > P2 | Two-tailed Pval |
| --- | --- | --- | --- | --- |
| ESN | LWK | 37 | 28 | 0.002563208 |
| EUR | ASW | 45 | 13 | 0.006608823 |
| CEU | GIH | 45 | 13 | 0.006608823 |
| SAS | CEU | 45 | 32 | 0.006608823 |
| CLM | CEU | 45 | 32 | 0.006608823 |
| EUR | CEU | 45 | 32 | 0.006608823 |
| CEU | IBS | 44 | 13 | 0.009559879 |
| CEU | BEB | 45 | 14 | 0.01609436 |
| FIN | IBS | 45 | 14 | 0.01609436 |
| ESN | GWD | 37 | 26 | 0.020073852 |

**Table S14:**
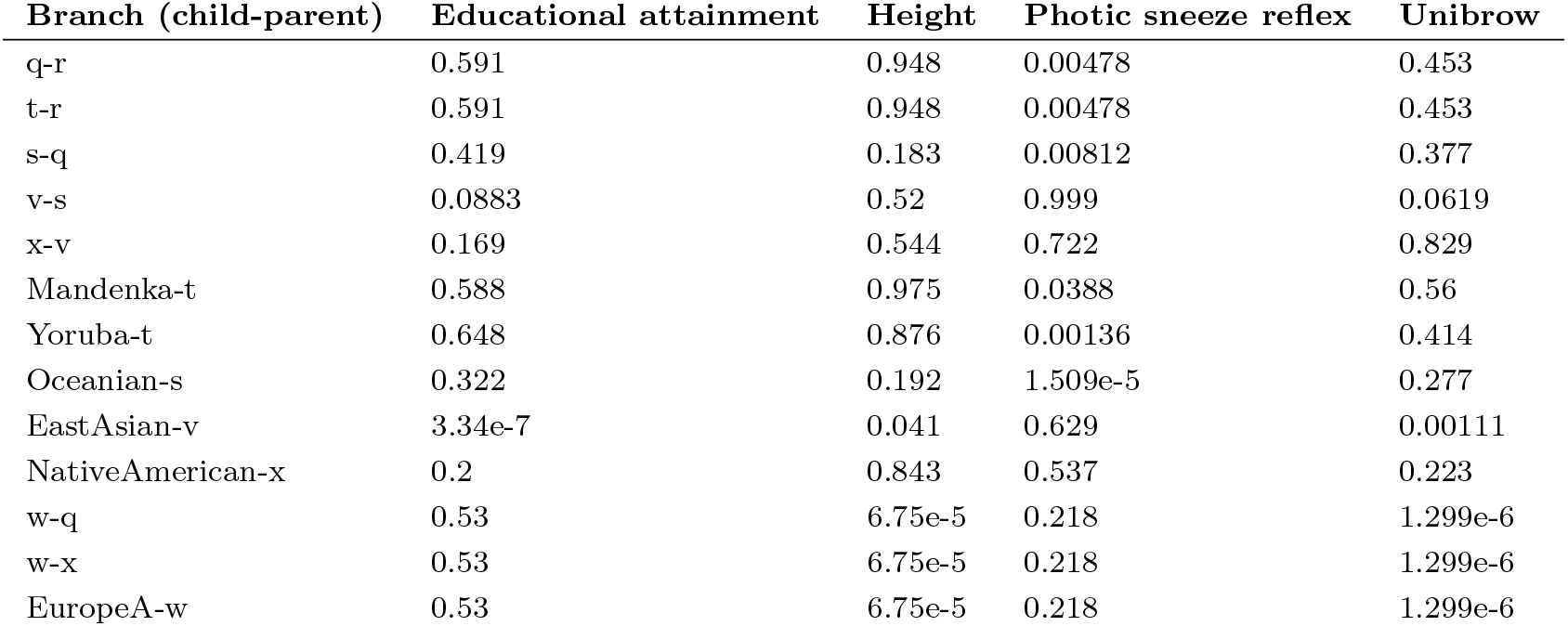
P-values from the *Q_B_* statistics of the 7-leaf population graph built using the Lazaridis et al. (2014) data, with the “EuropeA” panel, for traits with significant overall evidence of selection (P-value of *Q_X_* < 0.05/no. of GWAS tested).

**Table S15:** P-values from the *Q_B_* statistics of the 7-leaf population graph built using the Lazaridis et al. (2014) data, with the “EuropeB” panel, for traits with significant overall evidence of selection (P-value of *Q_X_* < 0.05/no. of GWAS tested).

| Branch (child-parent) | Age at voice drop | Educational attainment | Height | Photoc sneeze reflex | Unibrow |
| --- | --- | --- | --- | --- | --- |
| q-r | 0.935 | 0.648 | 0.965 | 0.00525 | 0.297 |
| t-r | 0.935 | 0.648 | 0.965 | 0.00525 | 0.297 |
| s-q | 0.562 | 0.385 | 0.123 | 0.0325 | 0.0682 |
| v-s | 0.114 | 0.0777 | 0.325 | 0.71 | 0.00514 |
| x-v | 0.0772 | 0.222 | 0.942 | 0.477 | 0.211 |
| y-q | 0.361 | 0.6321 | 0.0309 | 0.169 | 4.64e-5 |
| Mandenka-t | 0.938 | 0.641 | 0.965 | 0.0755 | 0.388 |
| Yoruba-t | 0.94 | 0.698 | 0.898 | 0.149 | 0.275 |
| Oceanian-s | 0.00221 | 0.362 | 0.217 | 5.72e-5 | 0.478 |
| EastAsian-v | 0.517 | 7.06e-7 | 0.0498 | 0.81 | 0.000278 |
| NativeAmerican-x | 0.0748 | 0.234 | 0.861 | 0.438 | 0.127 |
| Sardinian-y | 0.068 | 0.666 | 0.769 | 0.144 | 0.00134 |
| w-y | 0.757 | 0.631 | 2.75e-5 | 0.265 | 3.11e-6 |
| w-x | 0.757 | 0.631 | 2.75e-5 | 0.265 | 3.11e-6 |
| EuropeB-w | 0.757 | 0.631 | 2.75e-5 | 0.265 | 3.11e-6 |

**Table S16:** P-values from the *Q_B_* statistics of the 7-leaf population graph built using the Lazaridis et al. (2014) data, with the “EuropeC” panel, for traits with significant overall evidence of selection (P-value of *Q_X_* < 0.05/no. of GWAS tested).

| Branch (child-parent) | Educational attainment | Photoc sneeze reflex | Unibrow |
| --- | --- | --- | --- |
| q-r | 0.648 | 0.00525 | 0.297 |
| t-r | 0.648 | 0.00525 | 0.297 |
| s-q | 0.385 | 0.0325 | 0.0682 |
| v-s | 0.0777 | 0.71 | 0.00514 |
| x-v | 0.222 | 0.477 | 0.211 |
| y-q | 0.632 | 0.169 | 4.64e-5 |
| Mandenka-t | 0.641 | 0.0367 | 0.388 |
| Yoruba-t | 0.698 | 0.00159 | 0.275 |
| Oceanian-s | 0.362 | 5.72e-5 | 0.478 |
| EastAsian-v | 7.06e-7 | 0.81 | 0.000278 |
| NativeAmerican-x | 0.234 | 0.438 | 0.127 |
| Sardinian-y | 0.666 | 0.144 | 0.00134 |
| w-y | 0.631 | 0.265 | 3.11e-6 |
| w-x | 0.631 | 0.265 | 3.11e-6 |
| EuropeB-w | 0.631 | 0.265 | 3.11e-6 |

**Table S17:**
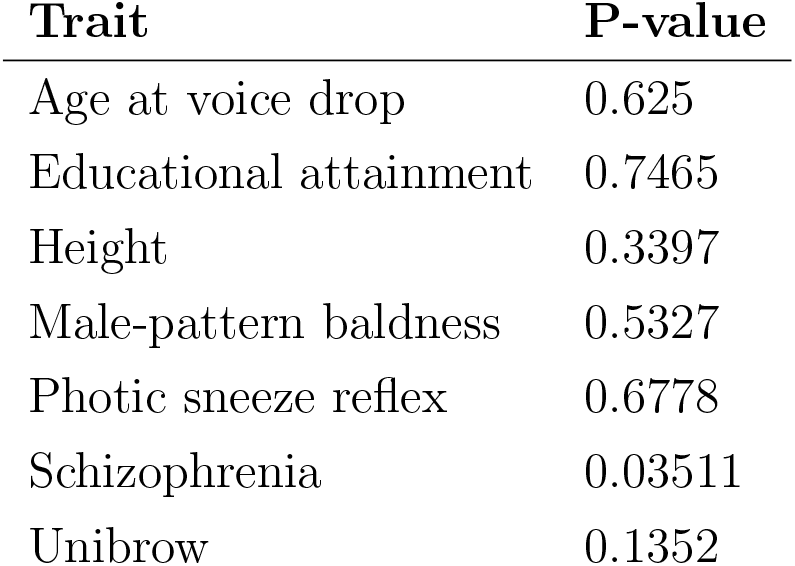
P-values from two-tailed binomial test performed to check if there were systematic biases in ancestral/derived allele polarity relative to the direction of the effect size, for all traits for which we found significant evidence of polygenic adaptation.

Table S18: Results from pairwise two-tailed binomial sign tests using the Lazaridis et al. (2014) data, for variants associated with self-reported age at voice drop. Each cell contains the number of SNPs in which the two panels have different frequencies, the number of SNPs in which the column panel has a higher frequency of the trait-increasing allele than the row panel and the P-value from the binomial test.

Table S19: Results from pairwise two-tailed binomial sign tests using the Lazaridis et al. (2014) data, for variants associated with educational attainment. Each cell contains the number of SNPs in which the two panels have different frequencies, the number of SNPs in which the column panel has a higher frequency of the trait-increasing allele than the row panel and the P-value from the binomial test.

Table S20: Results from pairwise two-tailed binomial sign tests using the Lazaridis et al. (2014) data, for variants associated with height. Each cell contains the number of SNPs in which the two panels have different frequencies, the number of SNPs in which the column panel has a higher frequency of the trait-increasing allele than the row panel and the P-value from the binomial test.

Table S21: Results from pairwise two-tailed binomial sign tests using the Lazaridis et al. (2014) data, for variants associated with self-reported photic sneeze reflex. Each cell contains the number of SNPs in which the two panels have different frequencies, the number of SNPs in which the column panel has a higher frequency of the trait-increasing allele than the row panel and the P-value from the binomial test.

Table S22: Results from pairwise two-tailed binomial sign tests using the Lazaridis et al. (2014) data, for variants associated with self-reported unibrow. Each cell contains the number of SNPs in which the two panels have different frequencies, the number of SNPs in which the column panel has a higher frequency of the trait-increasing allele than the row panel and the P-value from the binomial test.

**Table S23:** Top 10 most significant comparisons from pairwise two-tailed binomial sign tests using the Lazaridis et al. (2014) data, for variants associated with educational attainment. *N* = number of SNPs in which the two panels have different frequencies. *P*1 > *P*2 = number of SNPs in which Pop1 has a higher frequency of the trait-increasing allele than Pop2. Two-tailed Pval = P-value from the binomial test.

| Pop1 | Pop2 | N | P1 > P2 | Two-tailed Pval |
| --- | --- | --- | --- | --- |
| Yakut | Iranian_Jew | 71 | 54 | 1.25E-05 |
| Korean | Finnish | 70 | 53 | 1.92E-05 |
| Yi | Jordanian | 70 | 53 | 1.92E-05 |
| Jordanian | Han | 71 | 18 | 3.88E-05 |
| Mongola | Finnish | 71 | 53 | 3.88E-05 |
| Korean | Surui | 59 | 45 | 6.53E-05 |
| Yi | Iranian | 72 | 53 | 7.56E-05 |
| Japanese | Jordanian | 72 | 53 | 7.56E-05 |
| Tuscan | Korean | 72 | 19 | 7.56E-05 |
| Japanese | Kalash | 72 | 53 | 7.56E-05 |

**Table S24:**
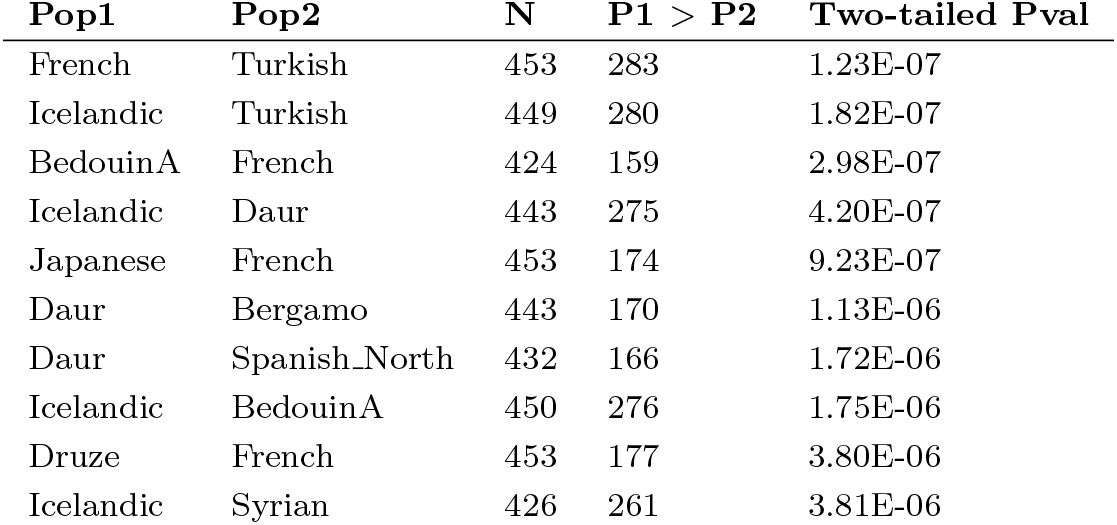
Top 10 most significant comparisons from pairwise two-tailed binomial sign tests using the Lazaridis et al. (2014) data, for variants associated with height. *N* = number of SNPs in which the two panels have different frequencies. *P*1 > *P*2 = number of SNPs in which Pop1 has a higher frequency of the trait-increasing allele than Pop2. Two-tailed Pval = P-value from the binomial test.

**Table S25:** Top 10 most significant comparisons from pairwise two-tailed binomial sign tests using the Lazaridis et al. (2014) data, for variants associated with self-reported photic sneeze reflex. *N* = number of SNPs in which the two panels have different frequencies. *P*1 > *P*2 = number of SNPs in which Pop1 has a higher frequency of the trait-increasing allele than Pop2. Two-tailed Pval = P-value from the binomial test.

| Pop1 | Pop2 | N | P1 > P2 | Two-tailed Pval |
| --- | --- | --- | --- | --- |
| Papuan | BantuSA | 39 | 32 | 7.03E-05 |
| Daur | Yoruba | 40 | 32 | 0.000182166 |
| Papuan | Biaka | 40 | 32 | 0.000182166 |
| Mende | She | 39 | 8 | 0.000294077 |
| Papuan | BantuKenya | 40 | 31 | 0.000679548 |
| Mende | Turkmen | 40 | 9 | 0.000679548 |
| Mende | Uzbek | 40 | 9 | 0.000679548 |
| Esan | Korean | 40 | 9 | 0.000679548 |
| Mende | Datog | 40 | 9 | 0.000679548 |
| Brahui | Turkish | 42 | 10 | 0.000940674 |

**Table S26:** Top 10 most significant comparisons from pairwise two-tailed binomial sign tests using the Lazaridis et al. (2014) data, for variants associated with self-reported unibrow. *N* = number of SNPs in which the two panels have different frequencies. *P*1 > *P*2 = number of SNPs in which Pop1 has a higher frequency of the trait-increasing allele than Pop2. Two-tailed Pval = P-value from the binomial test.

| **Pop1** | **Pop2** | **N** | **P1 > P2** | **Two-tailed Pval** |
| --- | --- | --- | --- | --- |
| Finnish | Kyrgyz | 33 | 7 | 0.001318727 |
| Bougainville | Finnish | 33 | 26 | 0.001318727 |
| Yemen | Bulgarian | 30 | 24 | 0.001430906 |
| Kusunda | Uzbek | 30 | 24 | 0.001430906 |
| Kusunda | Mordovian | 27 | 22 | 0.00151372 |
| Basque | Maltese | 32 | 7 | 0.002102402 |
| English | Yemen | 32 | 7 | 0.002102402 |
| Czech | Atayal | 32 | 7 | 0.002102402 |
| Yi | Nogai | 32 | 25 | 0.002102402 |
| Bougainville | Russian | 32 | 25 | 0.002102402 |

**Figure S1:**
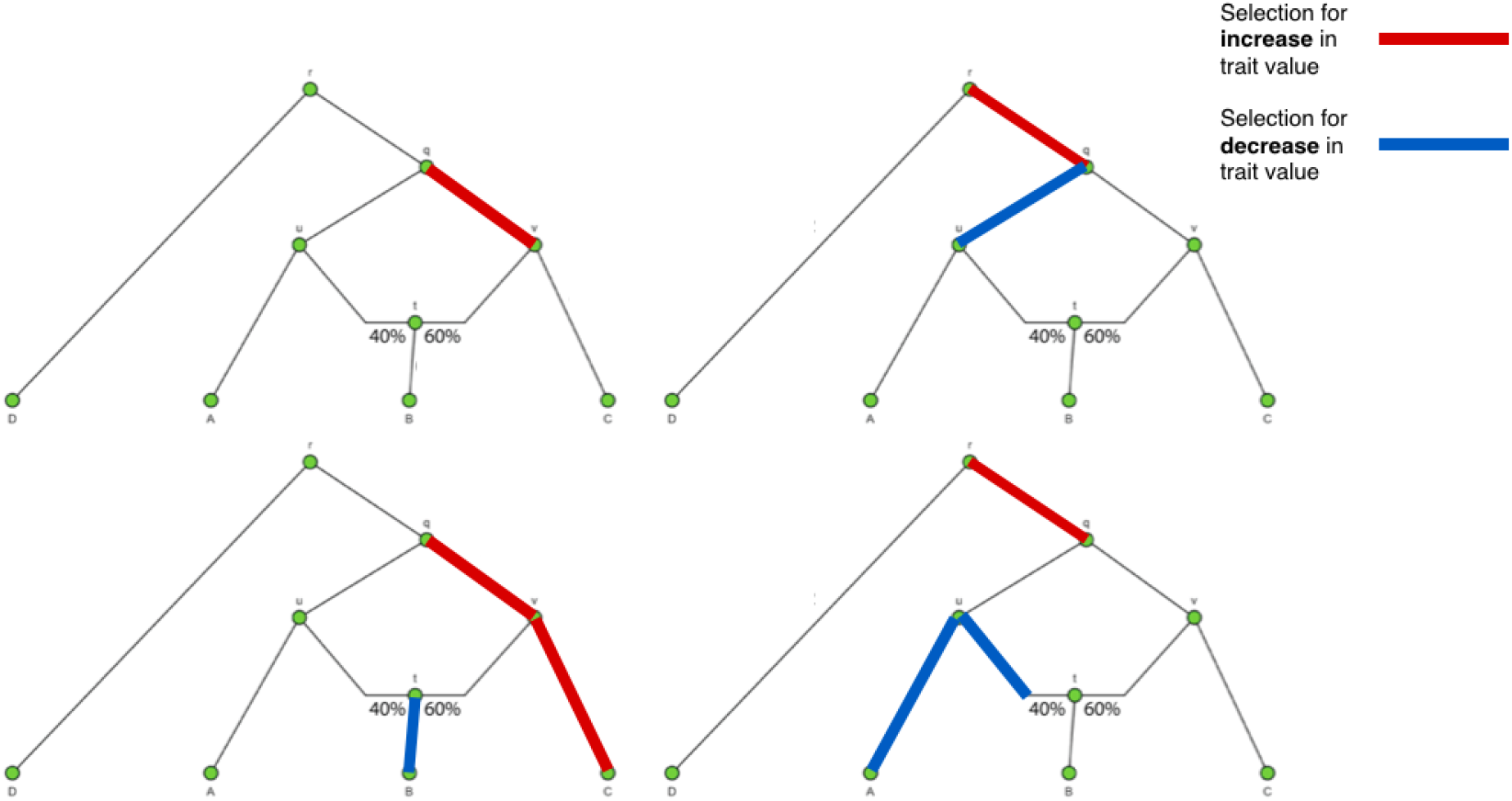
Four example scenarios in which different combinations of *α* parameters can produce very similar likelihood values.

**Figure S2:**
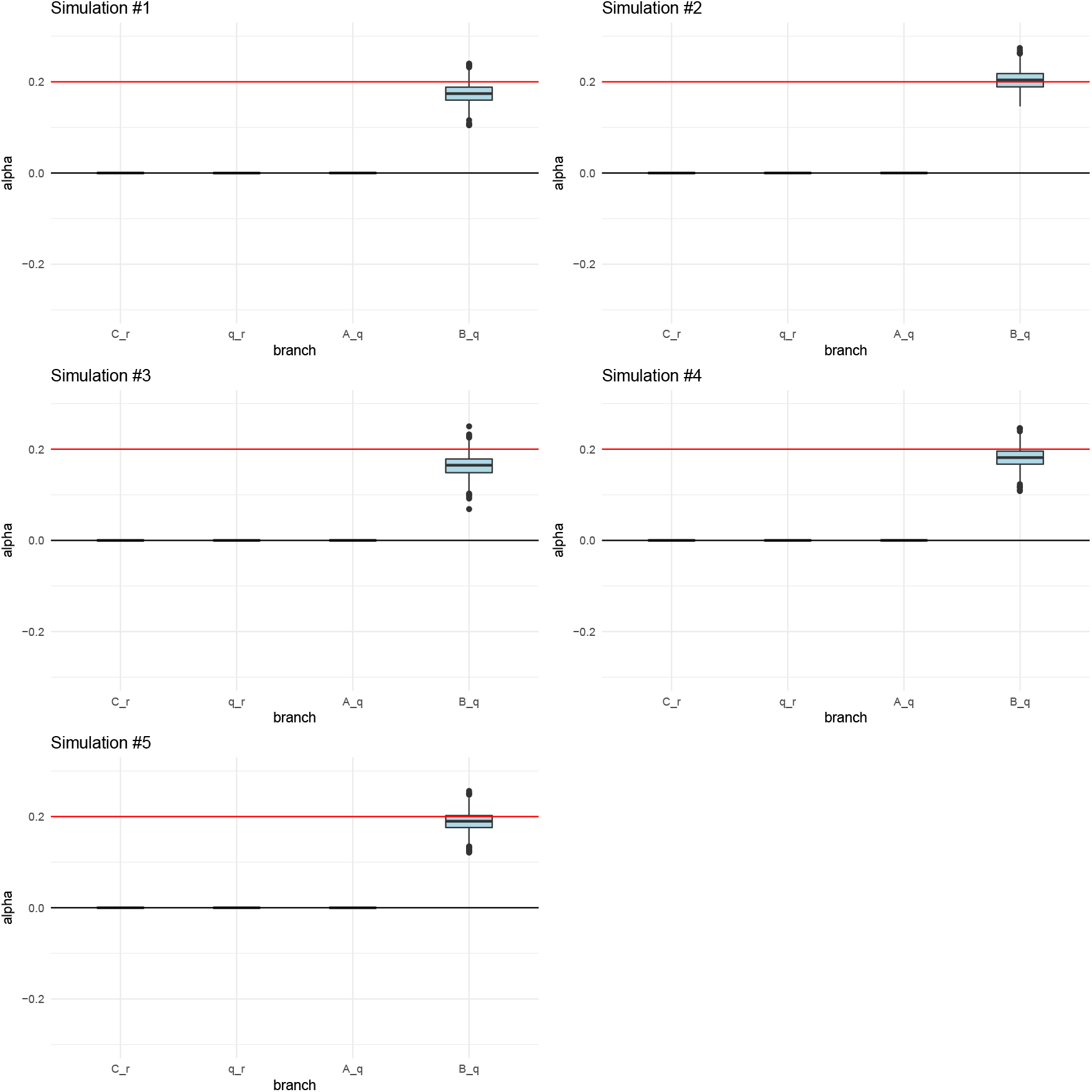
Posterior distributions of *α* parameters for five 400-SNPs simulations of a 3-leaf tree with small branch drift parameters (0.02) and *α* = 0.2 (red line) simulated in a terminal branch (B-q). The lower, middle and upper hinges denote the 25th, 50th and 75th percentiles, respectively. The upper whisker extends to the highest value that is within 1.5 * IQR of the upper hinge, where IQR is the inter-quartile range. The lower whisker extends to the lowest value within 1.5 * IQR of the lower hinge. Data beyond the whiskers are plotted as points.

**Figure S3:**
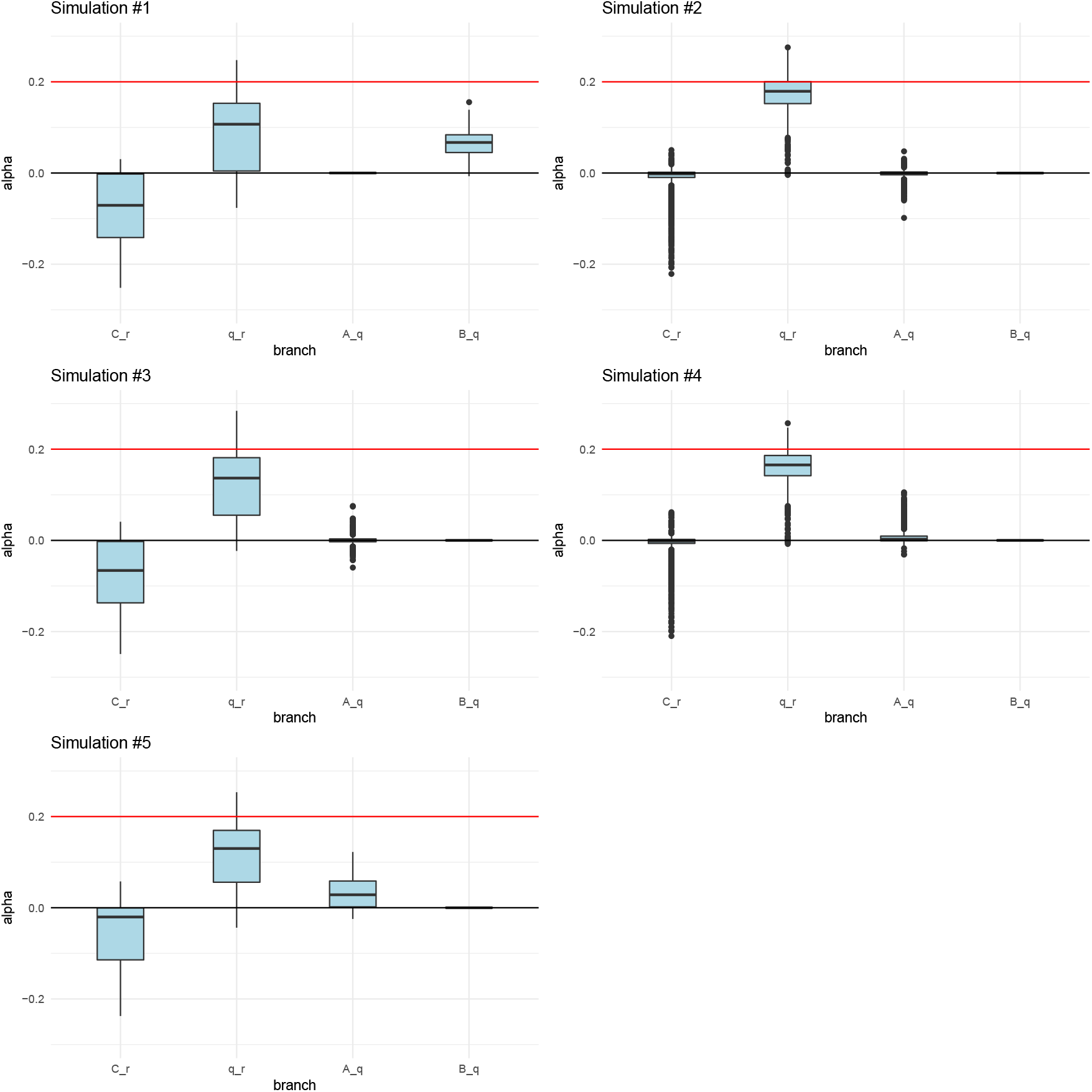
Posterior distributions of α parameters for five 400-SNP simulations of a 3-leaf tree with small branch drift parameters (0.02) and *α* = 0.2 (red line) simulated in an interior branch (q-r). The lower, middle and upper hinges denote the 25th, 50th and 75th percentiles, respectively. The upper whisker extends to the highest value that is within 1.5 * IQR of the upper hinge, where IQR is the inter-quartile range. The lower whisker extends to the lowest value within 1.5 * IQR of the lower hinge. Data beyond the whiskers are plotted as points.

**Figure S4:**
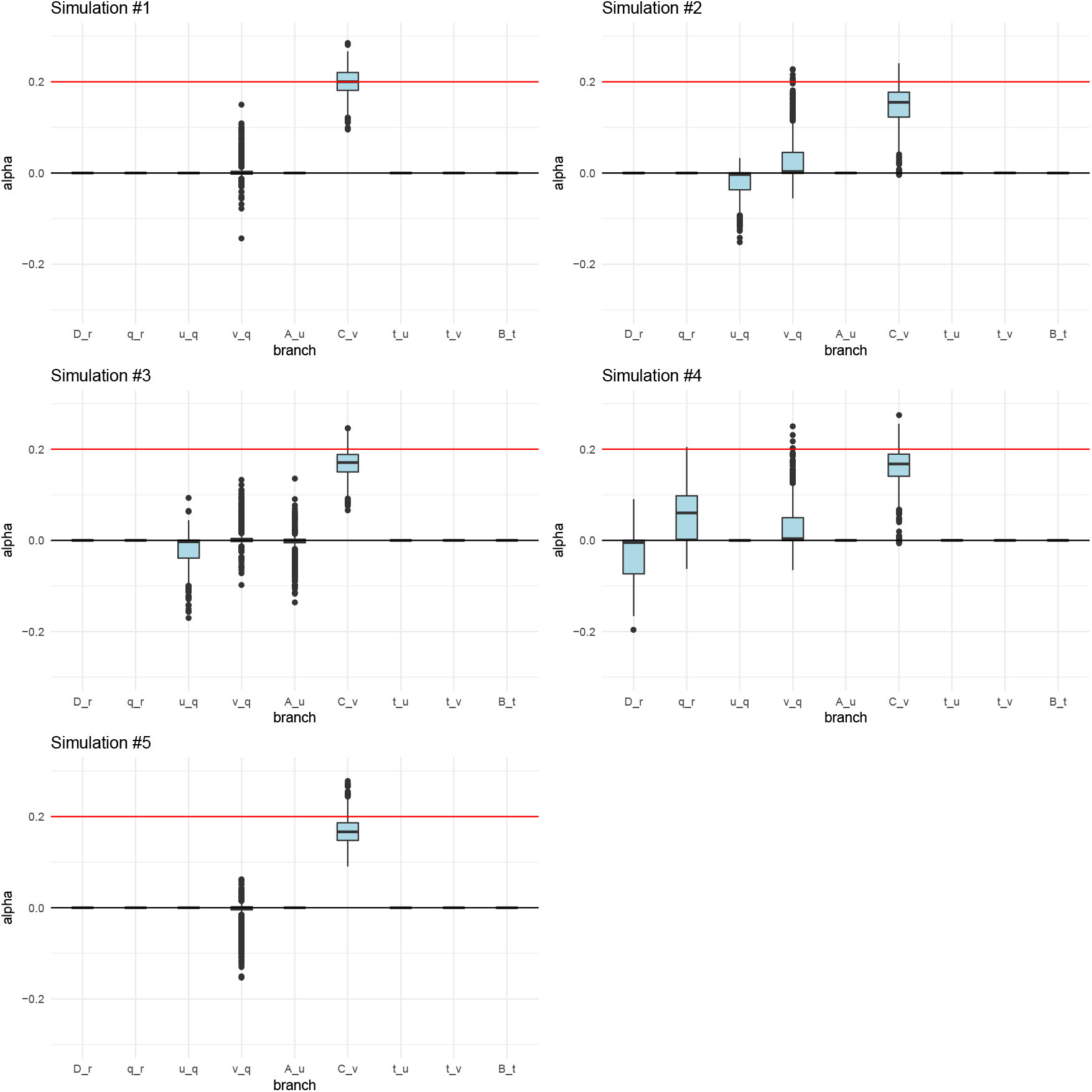
Posterior distributions of *α* parameters for five 400-SNPs simulations of a 4-leaf admixture graph with small branch drift parameters (0.02) and *α* = 0.2 (red line) simulated in a terminal branch (C-v). The upper and lower hinges denote the 25th and 75th percentiles. The lower, middle and upper hinges denote the 25th, 50th and 75th percentiles, respectively. The upper whisker extends to the highest value that is within 1.5 * IQR of the upper hinge, where IQR is the inter-quartile range. The lower whisker extends to the lowest value within 1.5 * IQR of the lower hinge. Data beyond the whiskers are plotted as points.

**Figure S5:**
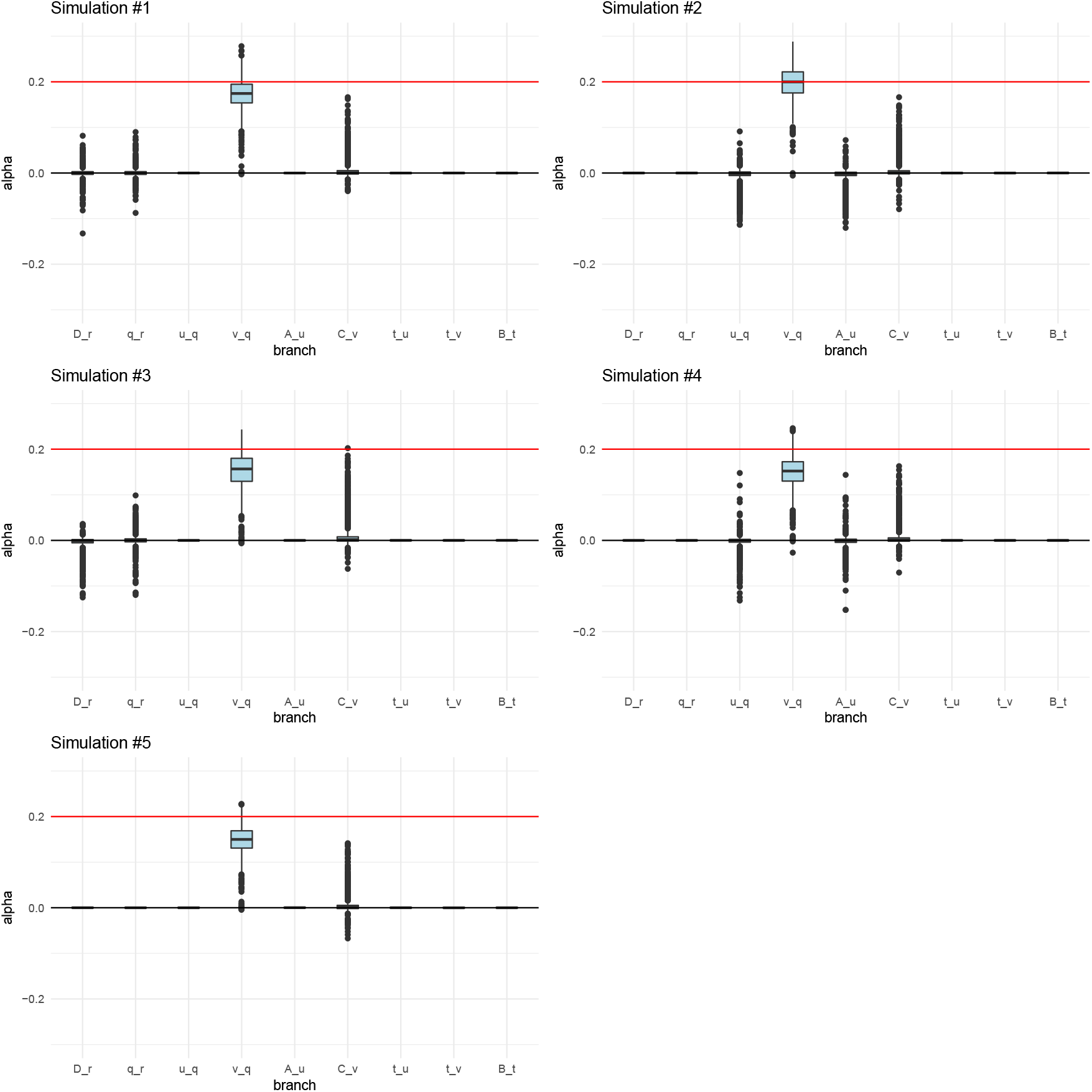
Posterior distributions of *α* parameters for five 400-SNPs simulations of a 4-leaf admixture graph with small branch drift parameters (0.02) and *α* = 0.2 (red line) simulated in an interior branch (v-q). The lower, middle and upper hinges denote the 25th, 50th and 75th percentiles, respectively. The upper whisker extends to the highest value that is within 1.5 * IQR of the upper hinge, where IQR is the inter-quartile range. The lower whisker extends to the lowest value within 1.5 * IQR of the lower hinge. Data beyond the whiskers are plotted as points.

**Figure S6:**
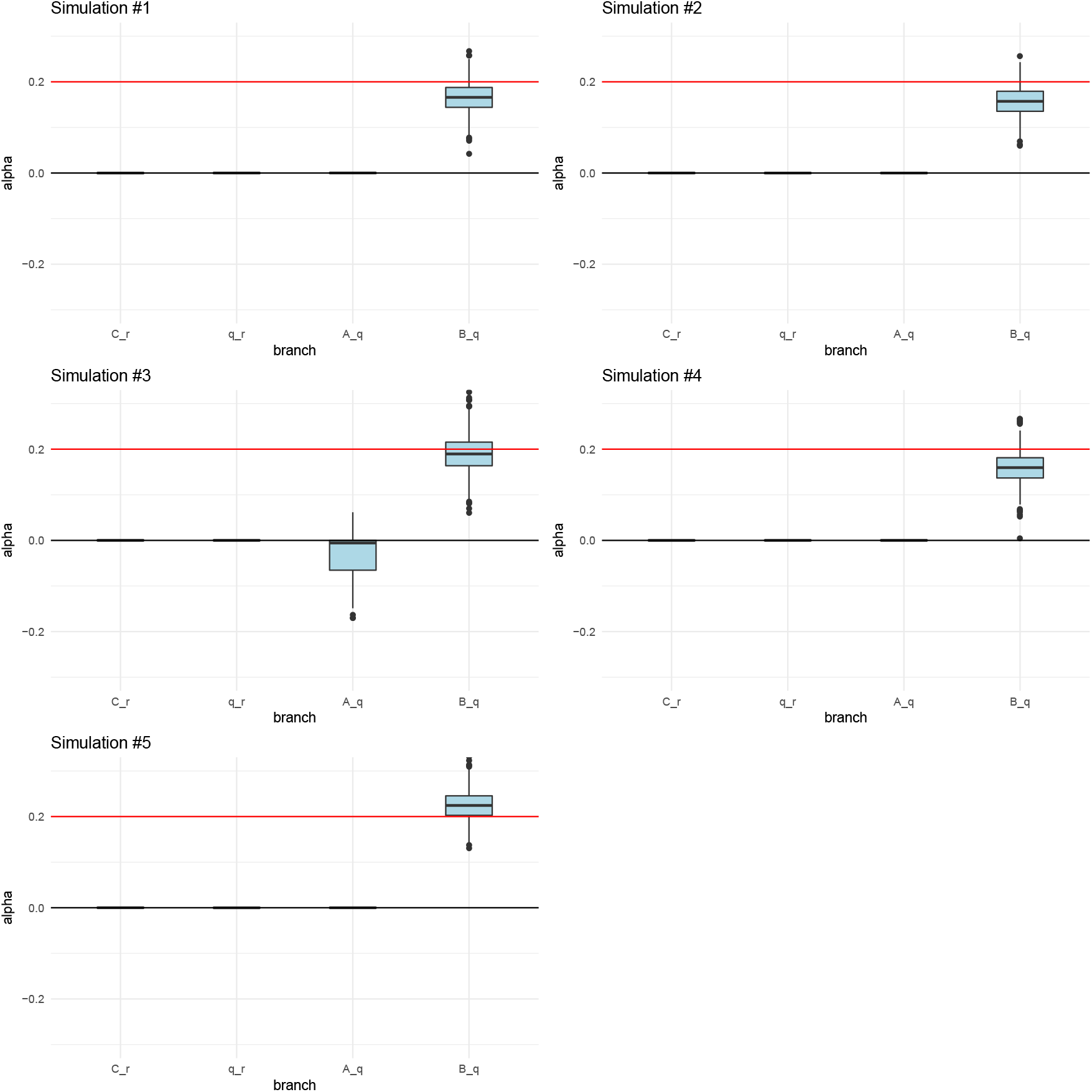
Posterior distributions of *α* parameters for five 400-SNPs simulations of a 3-leaf tree with large branch drift parameters (0.05) and *α* = 0.2 (red line) simulated in a terminal branch (B-q). The lower, middle and upper hinges denote the 25th, 50th and 75th percentiles, respectively. The upper whisker extends to the highest value that is within 1.5 * IQR of the upper hinge, where IQR is the inter-quartile range. The lower whisker extends to the lowest value within 1.5 * IQR of the lower hinge. Data beyond the whiskers are plotted as points.

**Figure S7:**
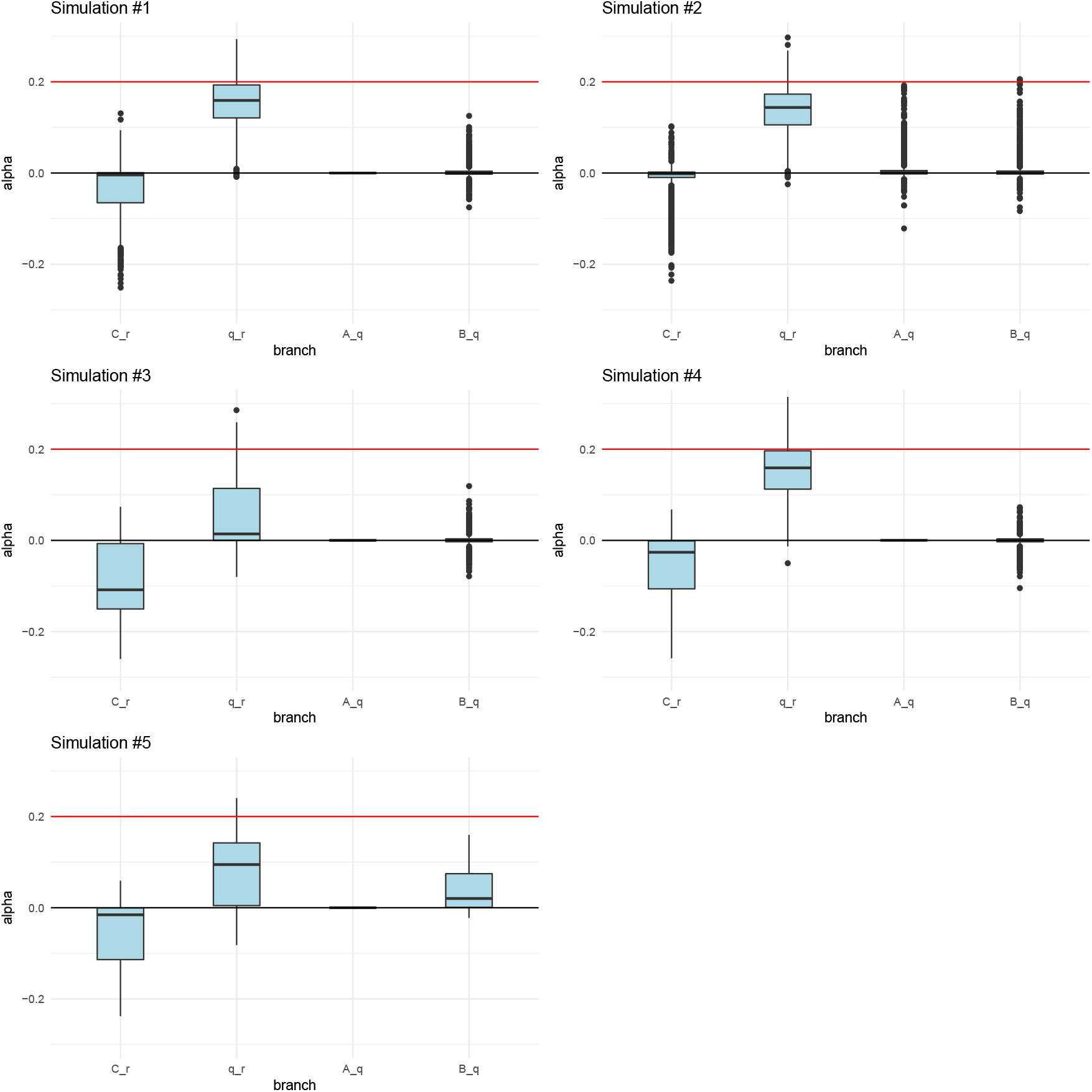
Posterior distributions of *α* parameters for five 400-SNPs simulations of a 3-leaf tree with large branch drift parameters (0.05) and *α* = 0.2 (red line) simulated in an interior branch (q-r). The lower, middle and upper hinges denote the 25th, 50th and 75th percentiles, respectively. The upper whisker extends to the highest value that is within 1.5 * IQR of the upper hinge, where IQR is the inter-quartile range. The lower whisker extends to the lowest value within 1.5 * IQR of the lower hinge. Data beyond the whiskers are plotted as points.

**Figure S8:**
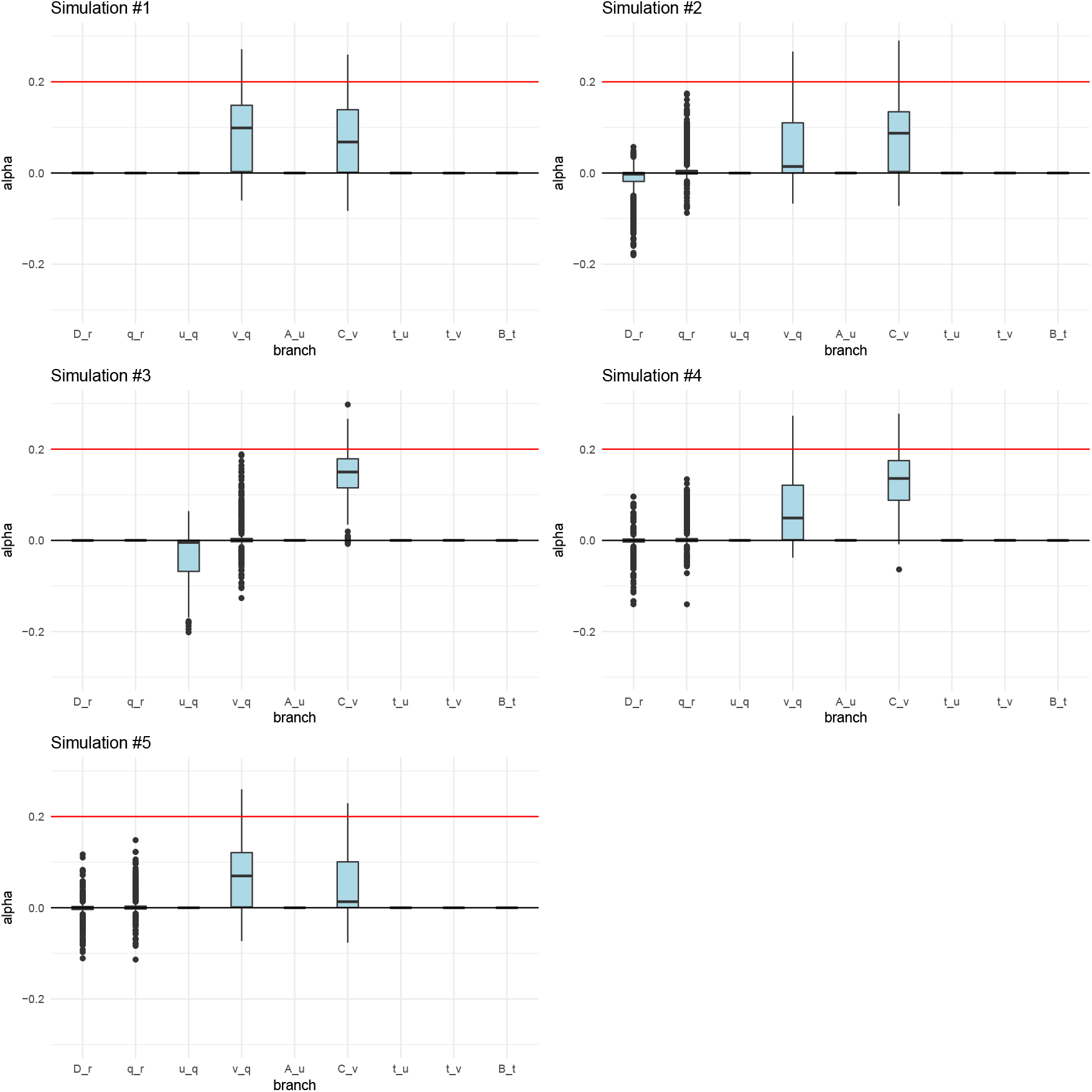
Posterior distributions of *α* parameters for five 400-SNPs simulations of a 4-leaf admixture graph with large branch drift parameters (0.05) and *α* = 0.2 (red line) simulated in a terminal branch (C-v). The lower, middle and upper hinges denote the 25th, 50th and 75th percentiles, respectively. The upper whisker extends to the highest value that is within 1.5 * IQR of the upper hinge, where IQR is the inter-quartile range. The lower whisker extends to the lowest value within 1.5 * IQR of the lower hinge. Data beyond the whiskers are plotted as points.

**Figure S9:**
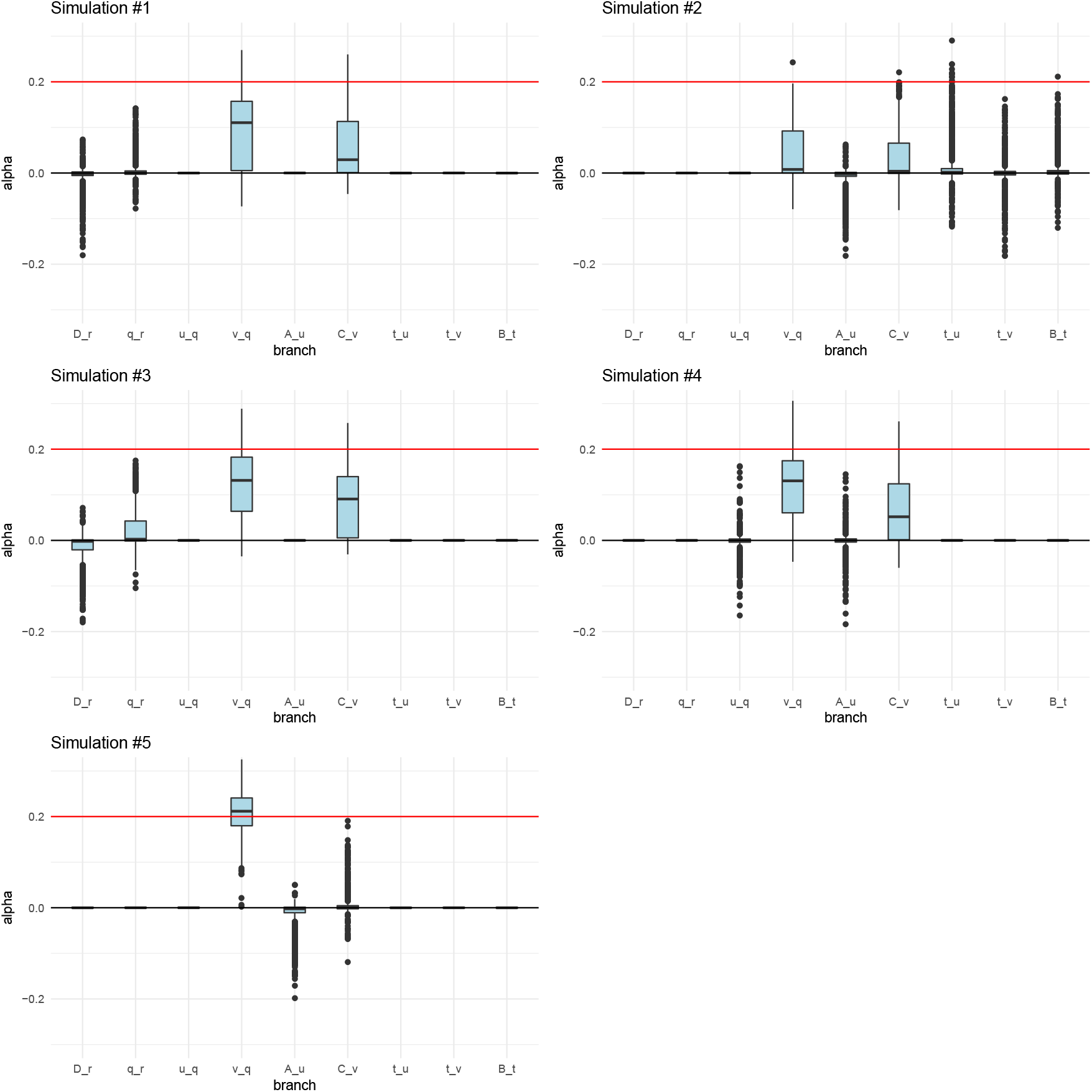
Posterior distributions of *α* parameters for five 400-SNPs simulations of a 4-leaf admixture graph with large branch drift parameters (0.05) and *α* = 0.2 (red line) simulated in an interior branch (v-q). The lower, middle and upper hinges denote the 25th, 50th and 75th percentiles, respectively. The upper whisker extends to the highest value that is within 1.5 * IQR of the upper hinge, where IQR is the inter-quartile range. The lower whisker extends to the lowest value within 1.5 * IQR of the lower hinge. Data beyond the whiskers are plotted as points.

**Figure S10:**
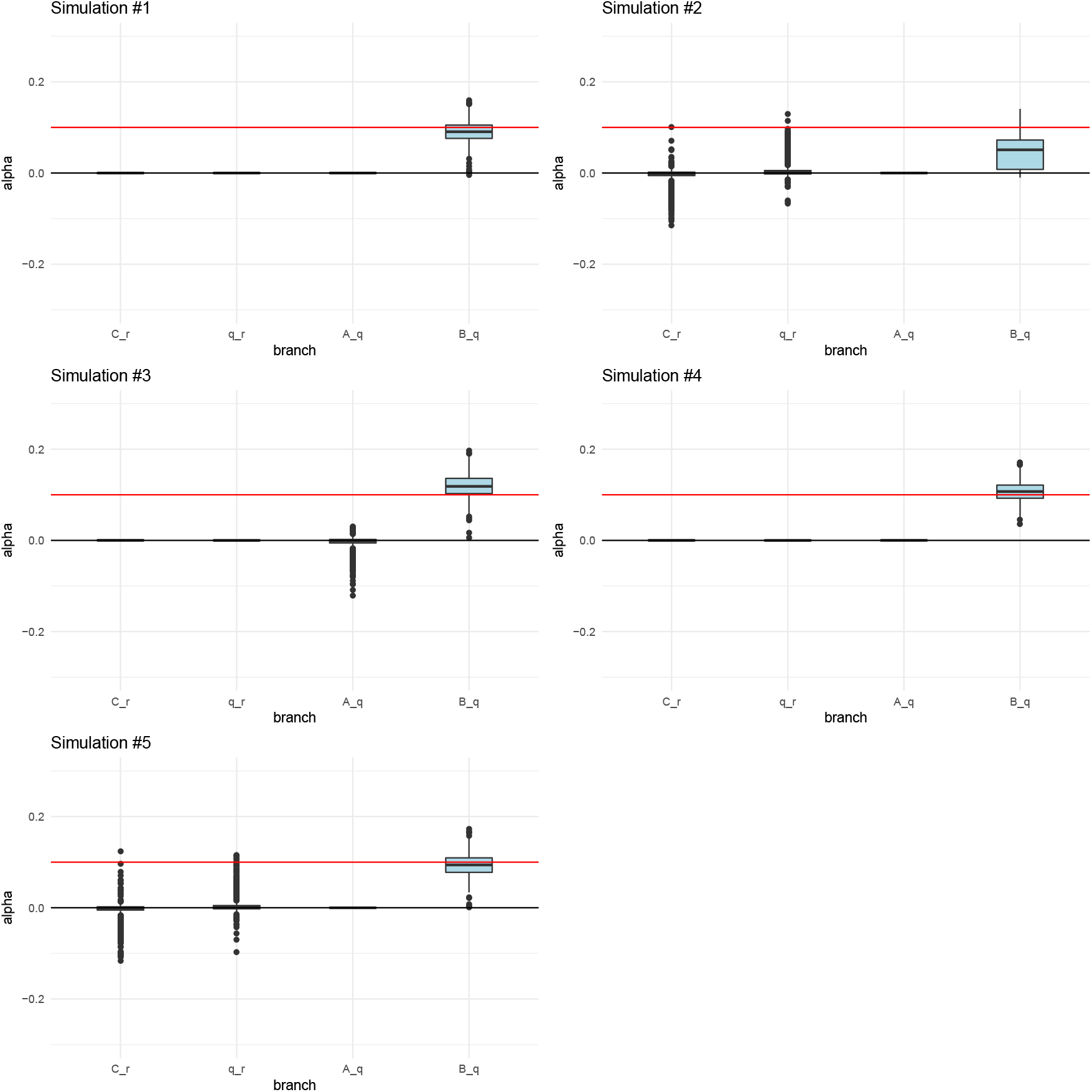
Posterior distributions of *α* parameters for five 400-SNPs simulations of a 3-leaf tree with small branch drift parameters (0.02) and *α* = 0.1 (red line) simulated in a terminal branch (B-q). The lower, middle and upper hinges denote the 25th, 50th and 75th percentiles, respectively. The upper whisker extends to the highest value that is within 1.5 * IQR of the upper hinge, where IQR is the inter-quartile range. The lower whisker extends to the lowest value within 1.5 * IQR of the lower hinge. Data beyond the whiskers are plotted as points.

**Figure S11:**
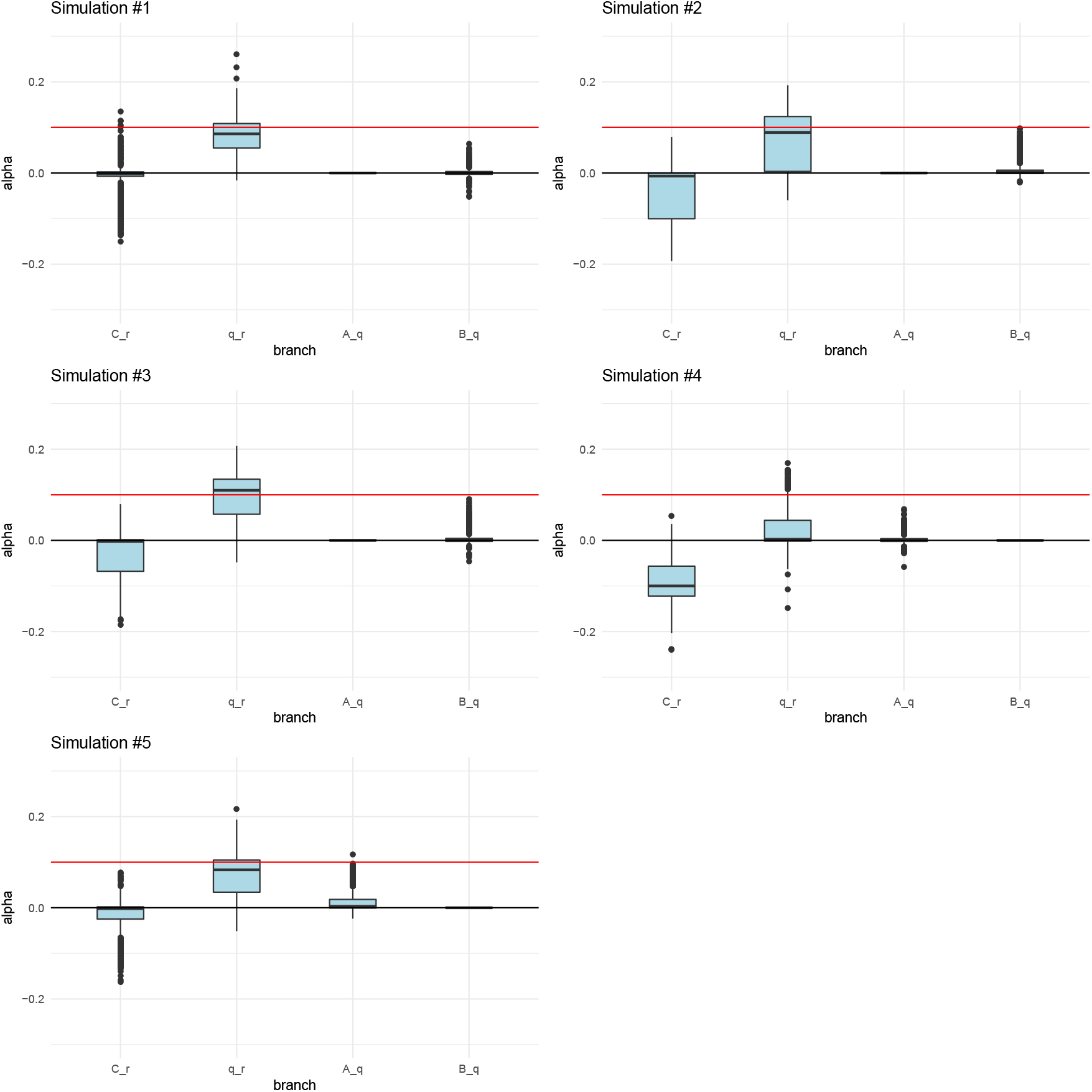
Posterior distributions of *α* parameters for five 400-SNPs simulations of a 3-leaf tree with small branch drift parameters (0.02) and *α* = 0.1 (red line) simulated in an interior branch (q-r). The lower, middle and upper hinges denote the 25th, 50th and 75th percentiles, respectively. The upper whisker extends to the highest value that is within 1.5 * IQR of the upper hinge, where IQR is the inter-quartile range. The lower whisker extends to the lowest value within 1.5 * IQR of the lower hinge. Data beyond the whiskers are plotted as points.

**Figure S12:**
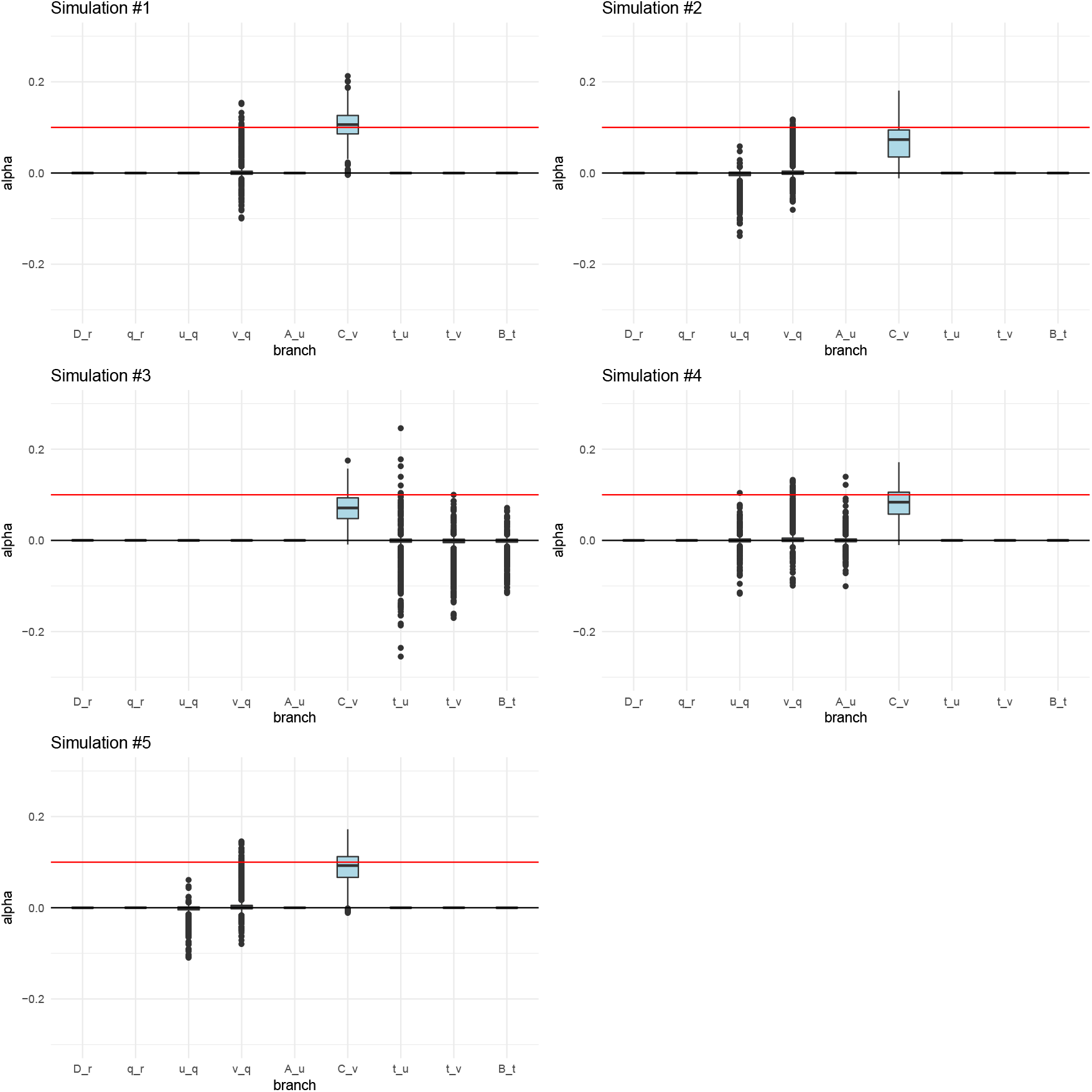
Posterior distributions of *α* parameters for five 400-SNPs simulations of a 4-leaf admixture graph with small branch drift parameters (0.02) and *α* = 0.1 (red line) simulated in a terminal branch (C-v). The lower, middle and upper hinges denote the 25th, 50th and 75th percentiles, respectively. The upper whisker extends to the highest value that is within 1.5 * IQR of the upper hinge, where IQR is the inter-quartile range. The lower whisker extends to the lowest value within 1.5 * IQR of the lower hinge. Data beyond the whiskers are plotted as points.

**Figure S13:**
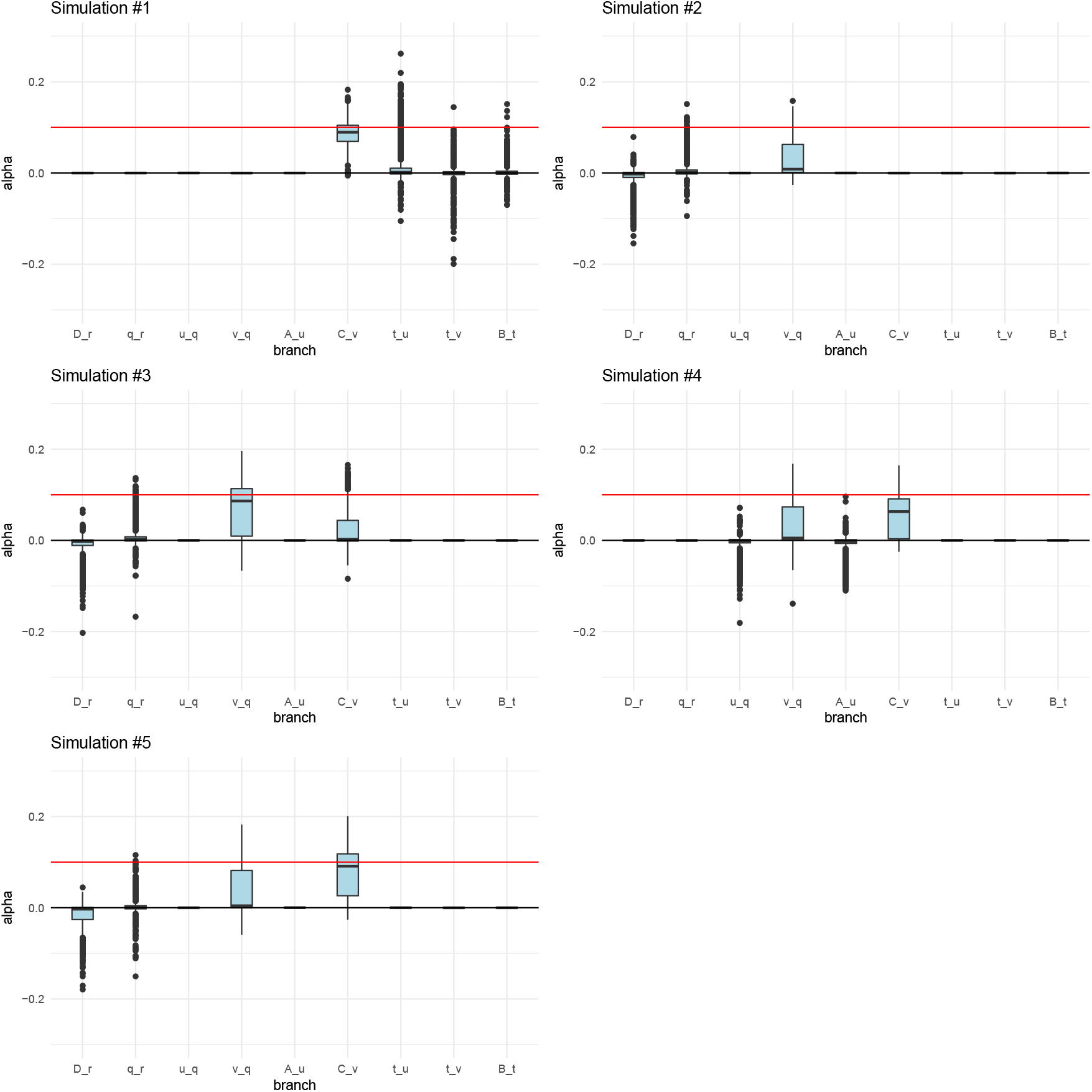
Posterior distributions of *α* parameters for five 400-SNPs simulations of a 4-leaf admixture graph with small branch drift parameters (0.02) and *α* = 0.1 (red line) simulated in an interior branch (v-q). The lower, middle and upper hinges denote the 25th, 50th and 75th percentiles, respectively. The upper whisker extends to the highest value that is within 1.5 * IQR of the upper hinge, where IQR is the inter-quartile range. The lower whisker extends to the lowest value within 1.5 * IQR of the lower hinge. Data beyond the whiskers are plotted as points.

**Figure S14:**
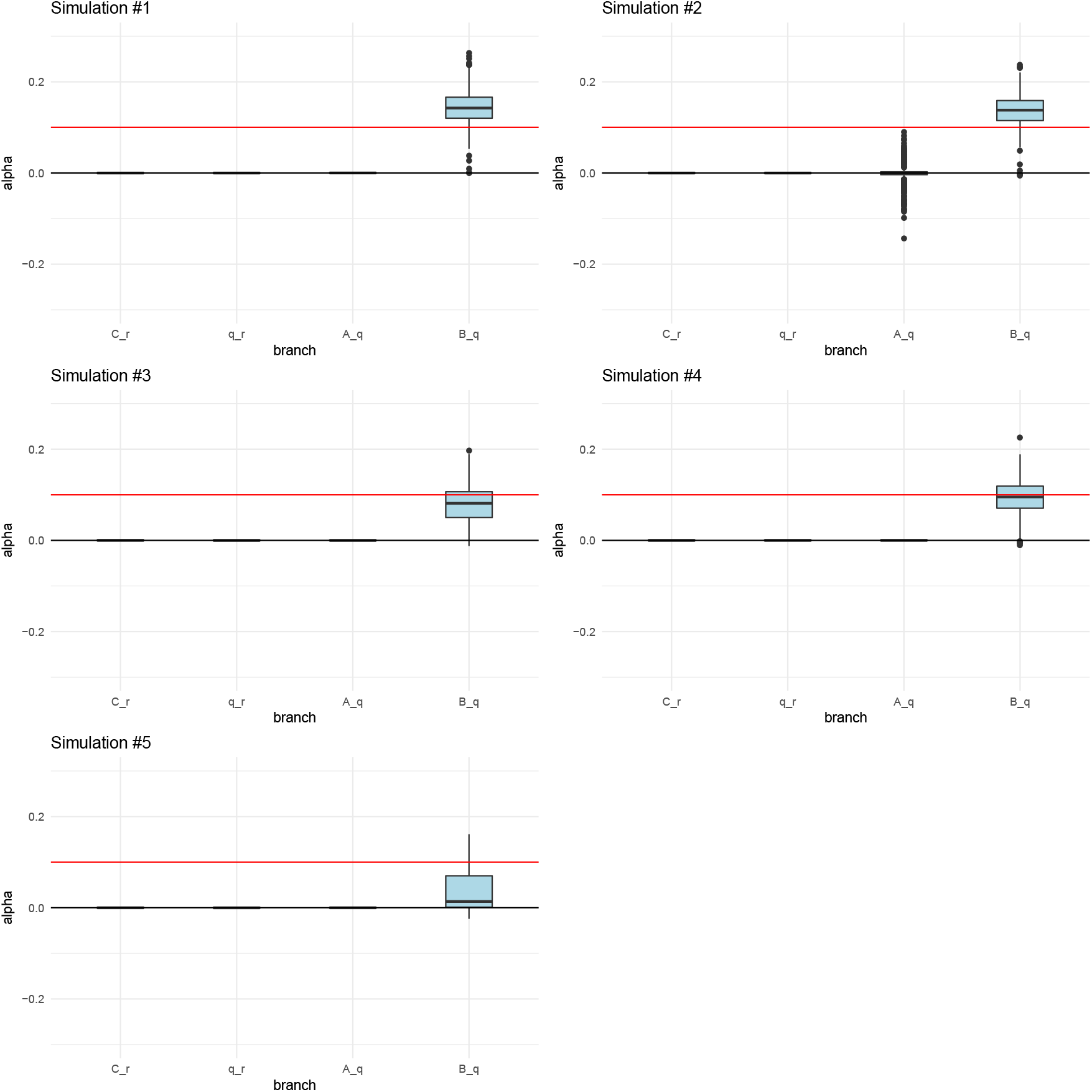
Posterior distributions of *α* parameters for five 400-SNPs simulations of a 3-leaf tree with large branch drift parameters (0.05) and *α* = 0.1 (red line) simulated in a terminal branch (B-q). The lower, middle and upper hinges denote the 25th, 50th and 75th percentiles, respectively. The upper whisker extends to the highest value that is within 1.5 * IQR of the upper hinge, where IQR is the inter-quartile range. The lower whisker extends to the lowest value within 1.5 * IQR of the lower hinge. Data beyond the whiskers are plotted as points.

**Figure S15:**
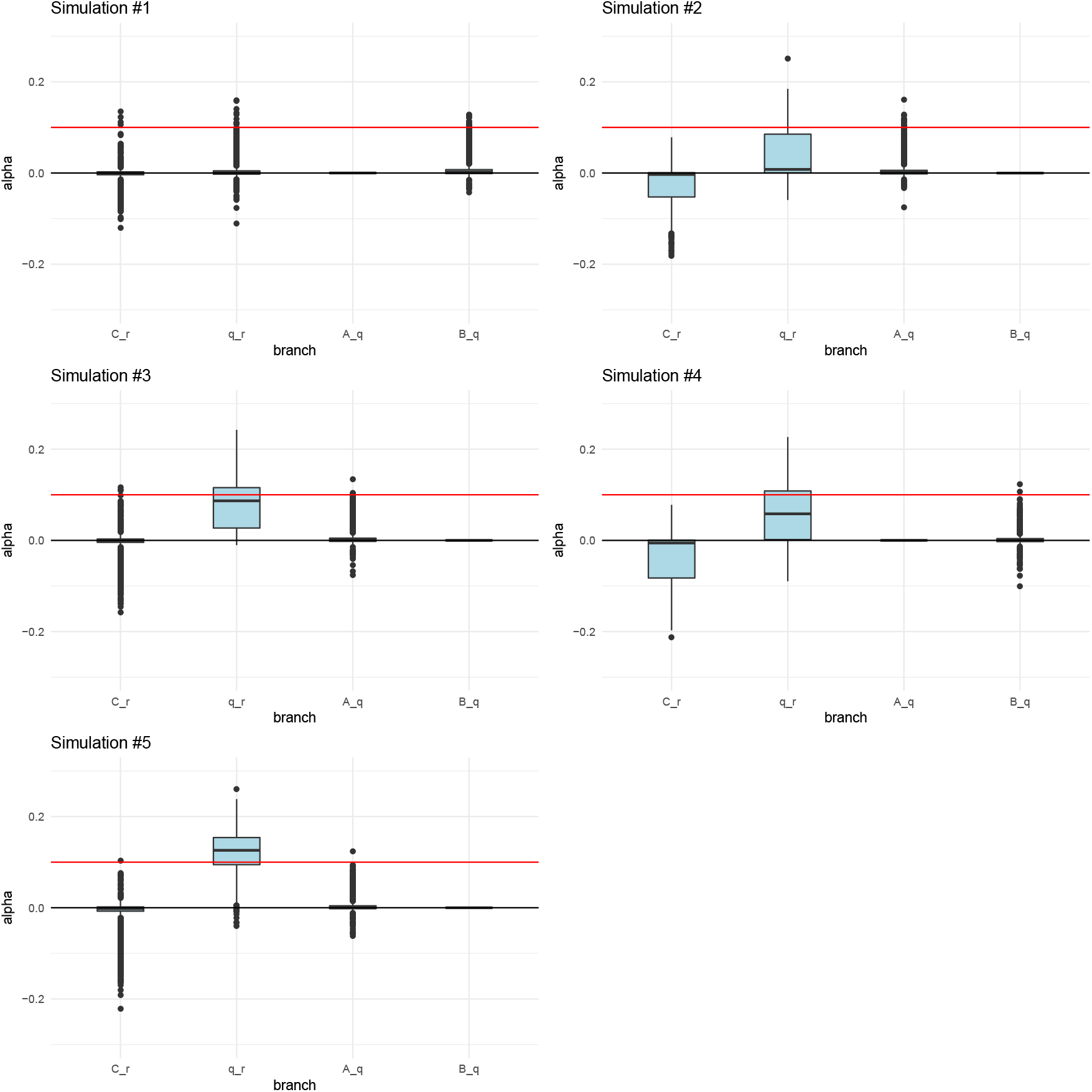
Posterior distributions of *α* parameters for five 400-SNPs simulations of a 3-leaf tree with large branch drift parameters (0.05) and *α* = 0.1 (red line) simulated in an interior branch (q-r). The lower, middle and upper hinges denote the 25th, 50th and 75th percentiles, respectively. The upper whisker extends to the highest value that is within 1.5 * IQR of the upper hinge, where IQR is the inter-quartile range. The lower whisker extends to the lowest value within 1.5 * IQR of the lower hinge. Data beyond the whiskers are plotted as points.

**Figure S16:**
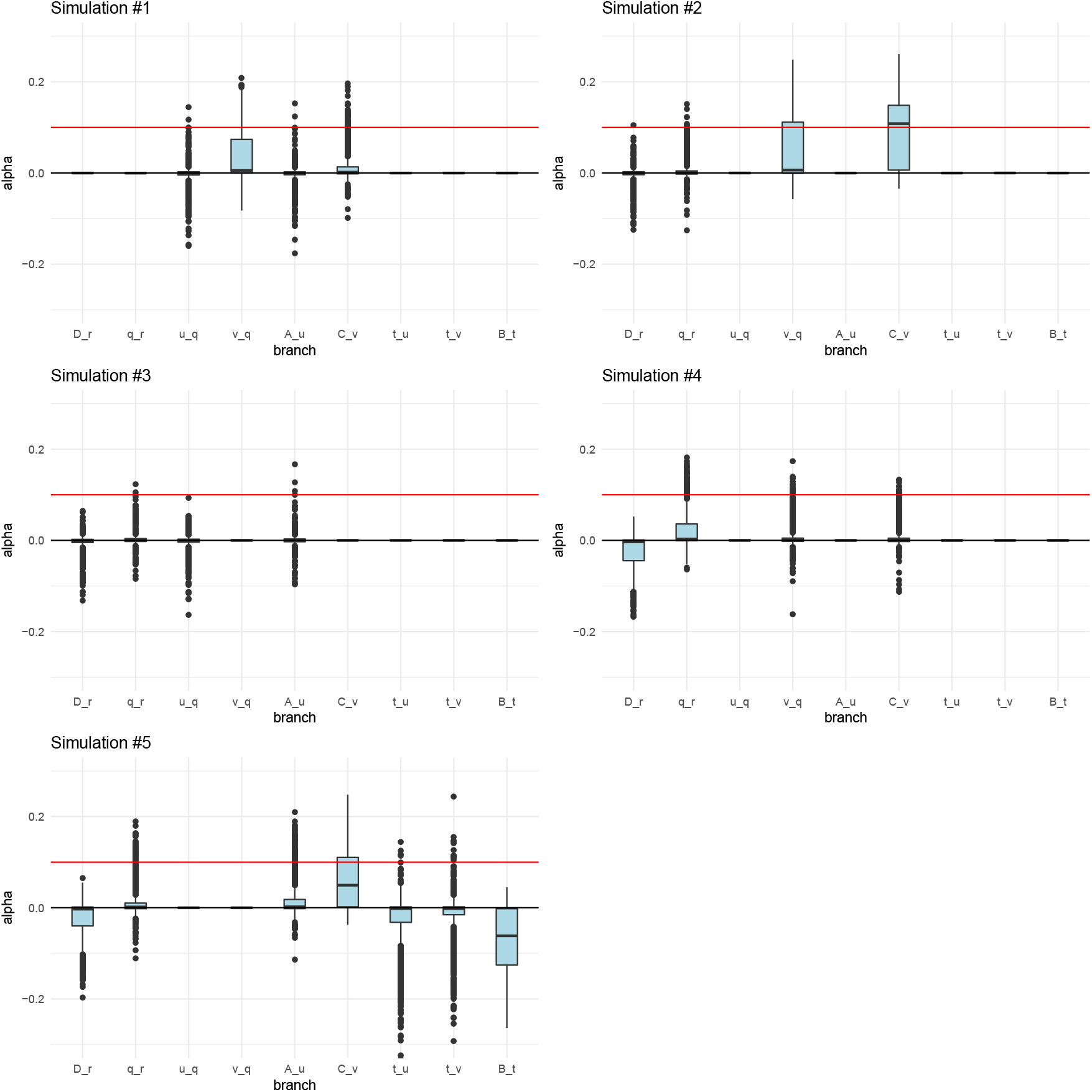
Posterior distributions of *α* parameters for five 400-SNPs simulations of a 4-leaf admixture graph with large branch drift parameters (0.05) and *α* = 0.1 (red line) simulated in a terminal branch (C-v). The lower, middle and upper hinges denote the 25th, 50th and 75th percentiles, respectively. The upper whisker extends to the highest value that is within 1.5 * IQR of the upper hinge, where IQR is the inter-quartile range. The lower whisker extends to the lowest value within 1.5 * IQR of the lower hinge. Data beyond the whiskers are plotted as points.

**Figure S17:**
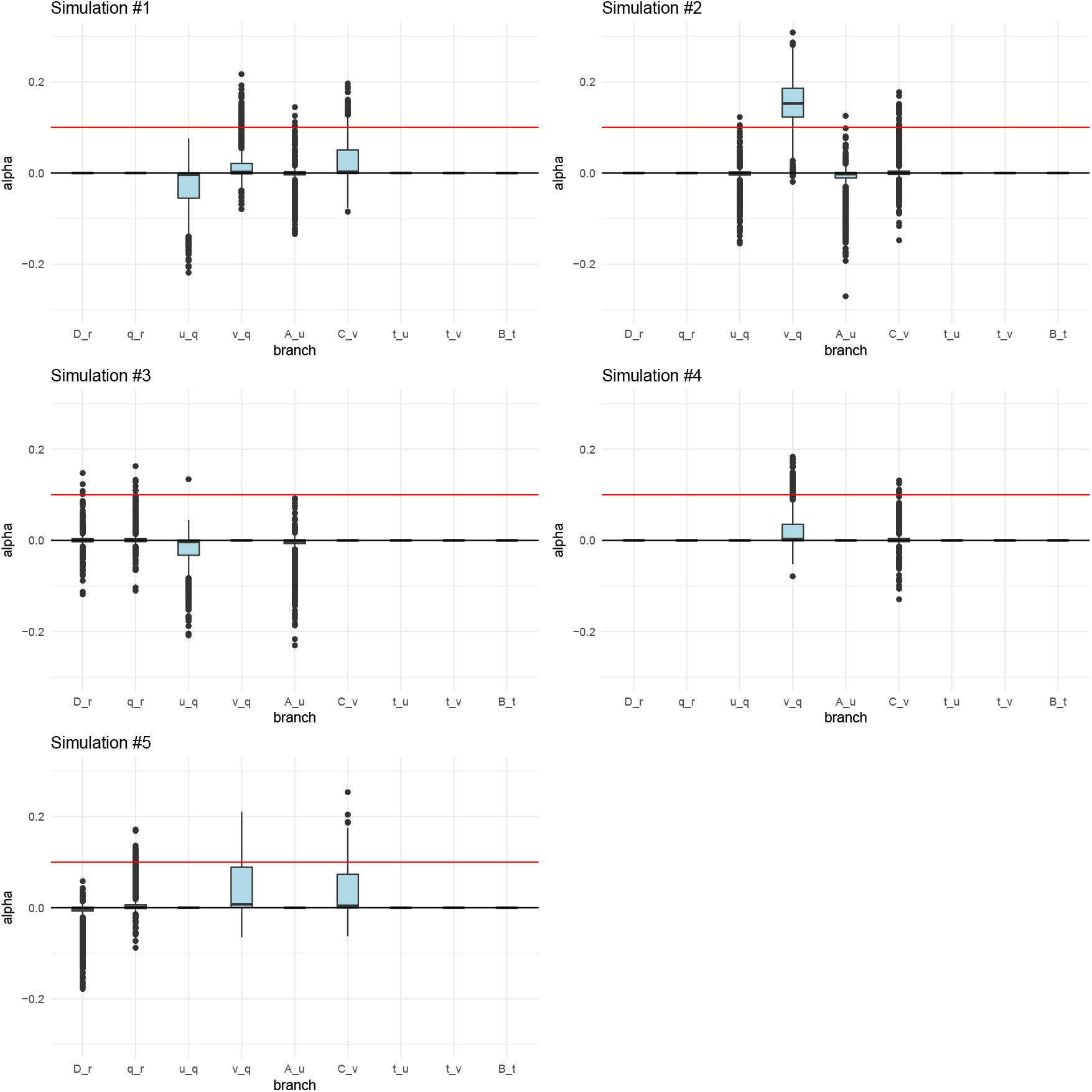
Posterior distributions of *α* parameters for five 400-SNPs simulations of a 4-leaf admixture graph with large branch drift parameters (0.05) and *α* = 0.1 (red line) simulated in an interior branch (v-q). The lower, middle and upper hinges denote the 25th, 50th and 75th percentiles, respectively. The upper whisker extends to the highest value that is within 1.5 * IQR of the upper hinge, where IQR is the inter-quartile range. The lower whisker extends to the lowest value within 1.5 * IQR of the lower hinge. Data beyond the whiskers are plotted as points.

**Figure S18:**
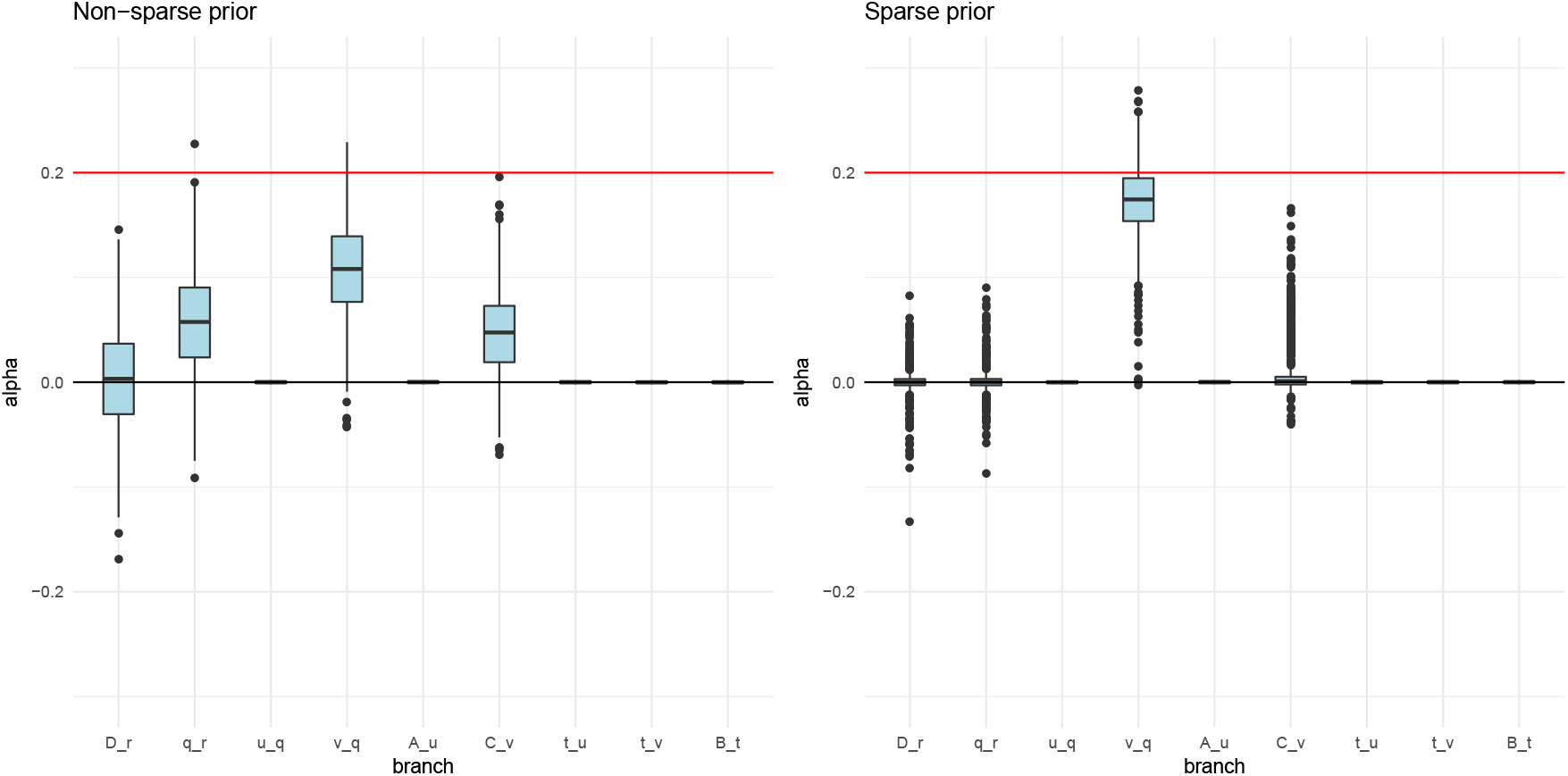
Posterior distributions of *α* parameters for two 400-SNPs simulations of a 4-leaf admixture graph with small branch drift parameters (0.02) and *α* = 0.2 (red line) simulated in an interior branch (v-q). The left panel shows a simulation whose parameters were estimated with a non-sparse prior (*κ* = 0) and the right panel shows a simulation whose parameters were estimated with a sparse prior, as defined in the main text.

**Figure S19:**
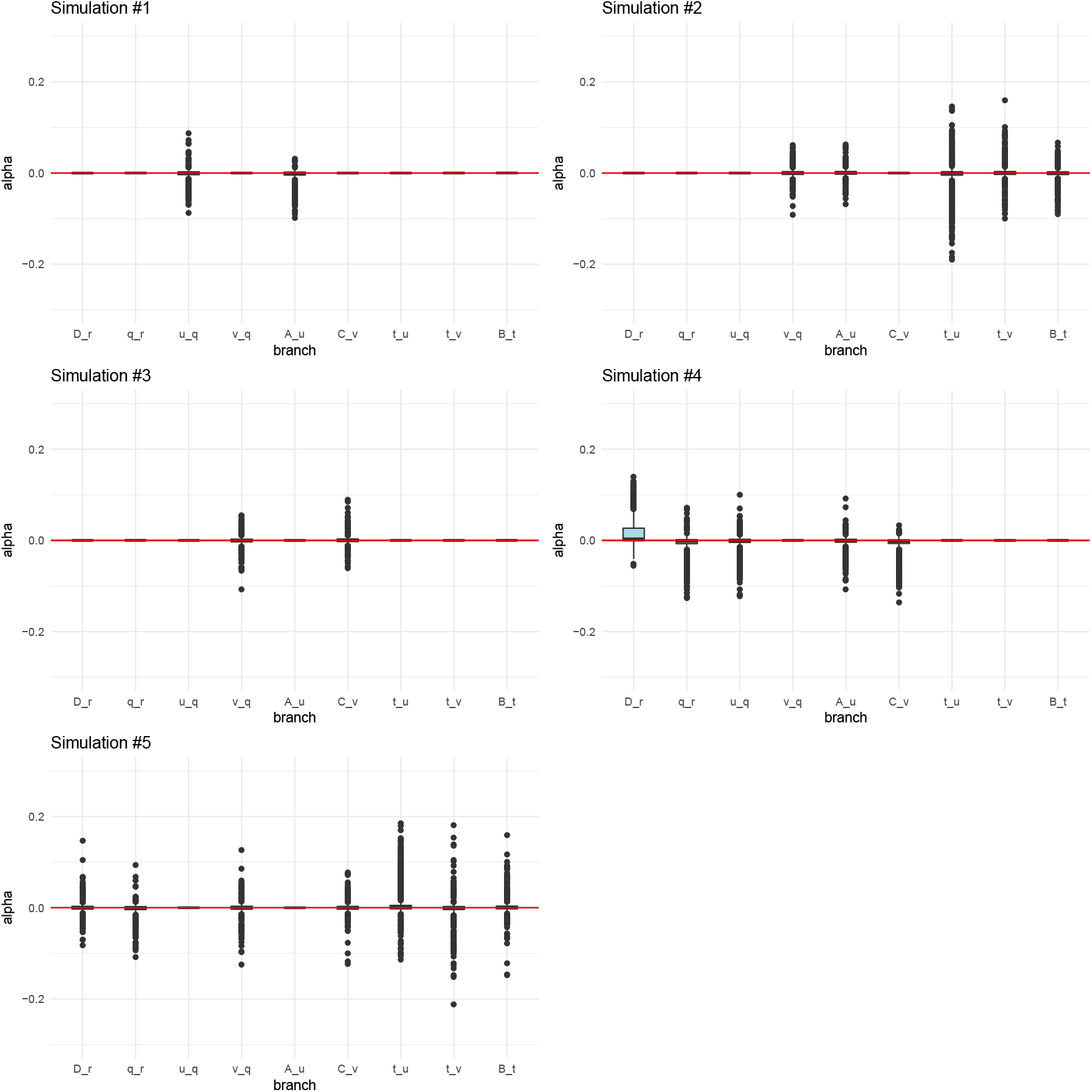
Posterior distributions of *α* parameters for five 400-SNPs neutral simulations of a 4-leaf admixture graph with drift parameters equal to 0.02 in each branch.

**Figure S20:**
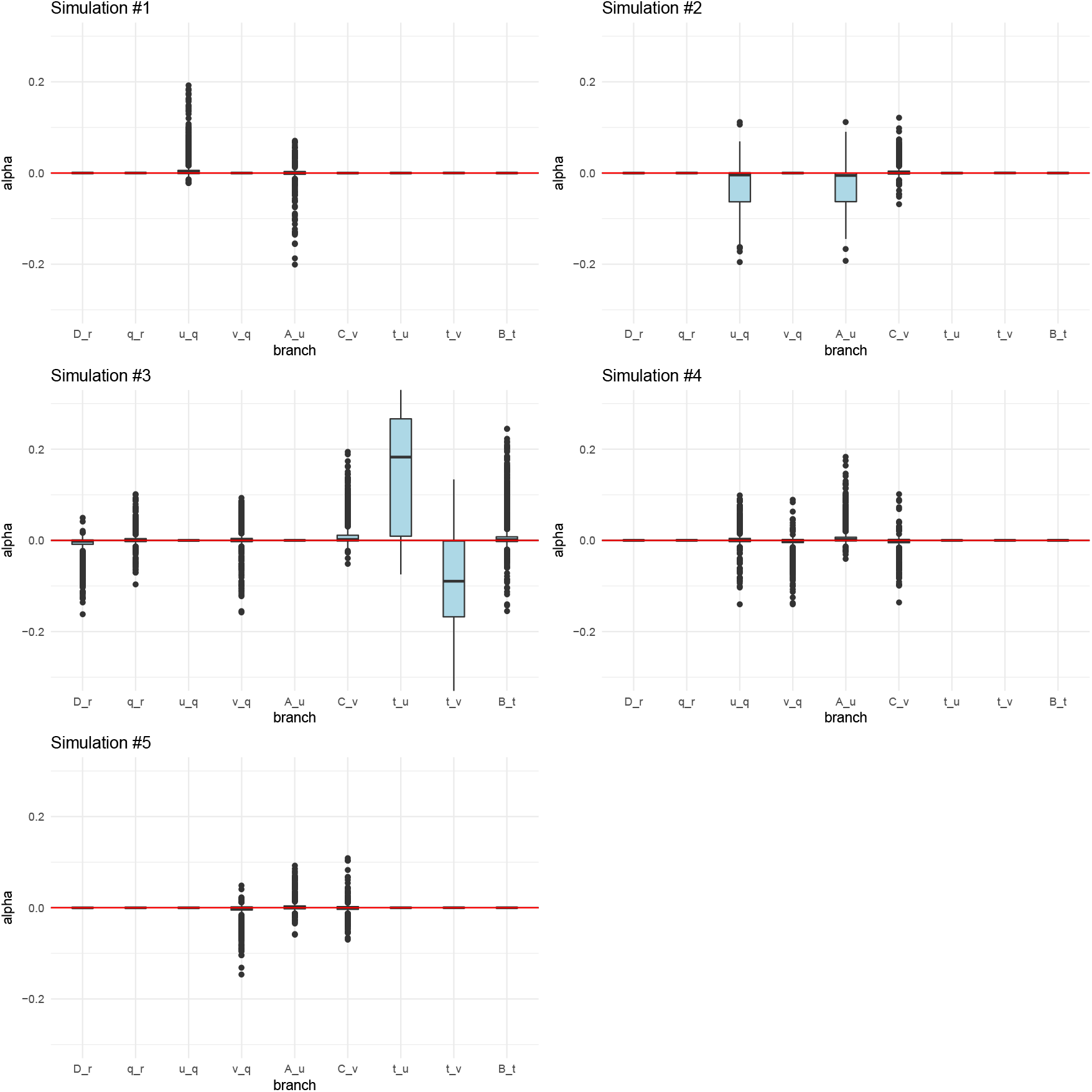
Posterior distributions of *α* parameters for five 400-SNPs neutral simulations of a 4-leaf admixture graph with drift parameters equal to 0.03 in each branch, but incorrectly specified to be equal to 0.02.

**Figure S21:**
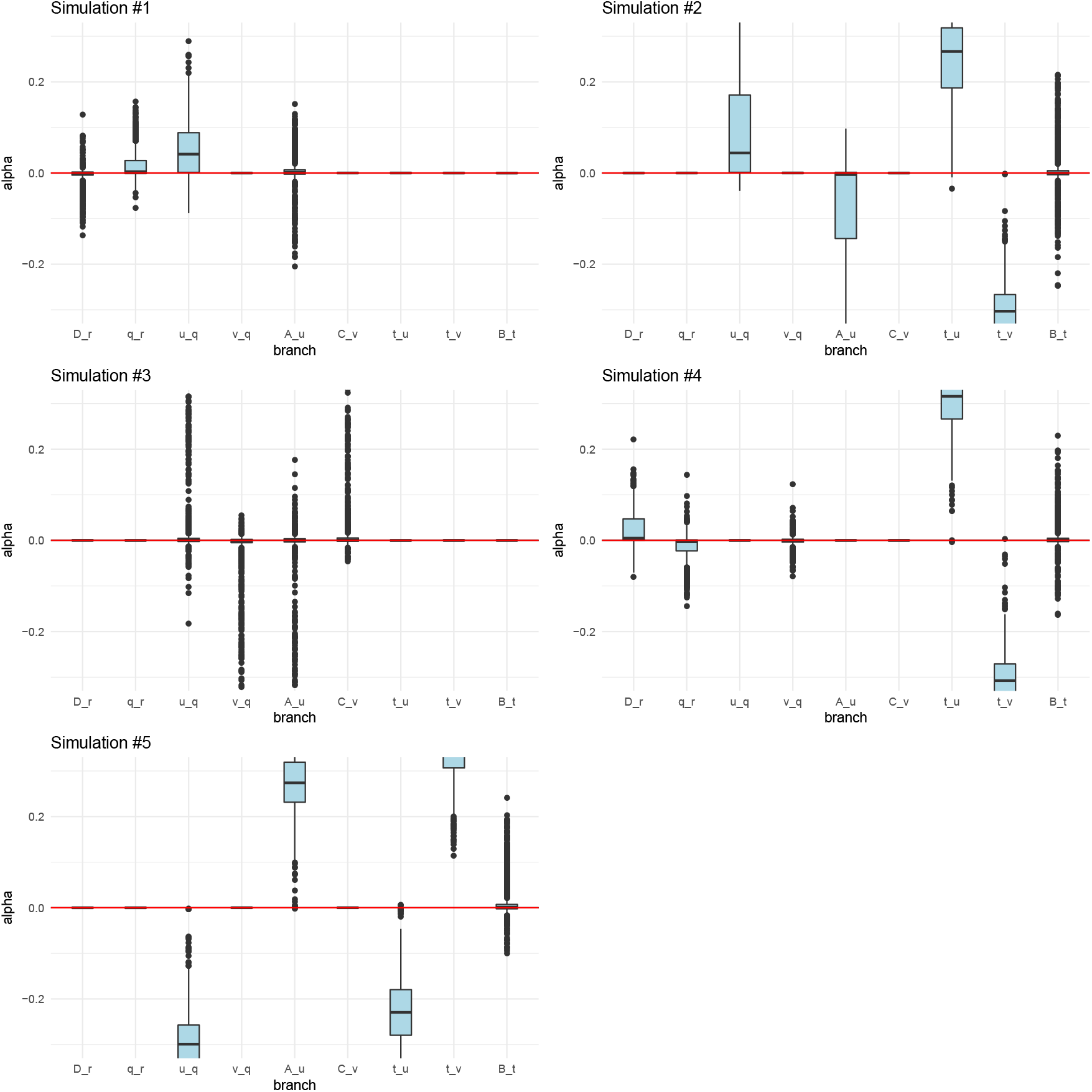
Posterior distributions of *α* parameters for five 400-SNPs neutral simulations of a 4-leaf admixture graph with drift parameters equal to 0.04 in each branch, but incorrectly specified to be equal to 0.02.

**Figure S22:**
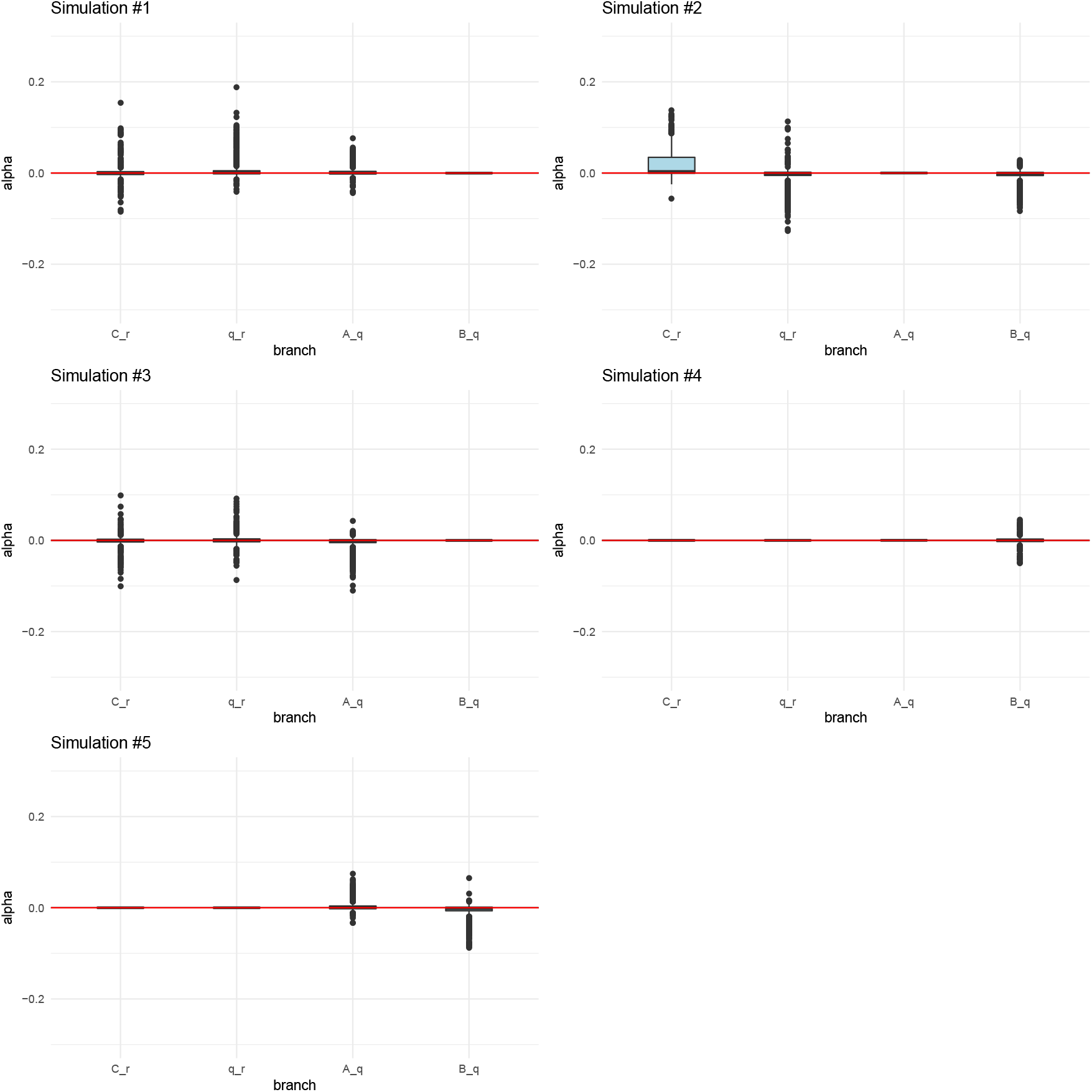
Posterior distributions of *α* parameters for five 400-SNPs neutral simulations of a 4-leaf admixture graph with drift parameters equal to 0.02 in each branch. We pretended one of the populations (A) was not sampled, and specified the topology to be a 3-leaf tree with drift parameters equal to 0.02 in each branch. Populations B, C and D were relabeled to be A, B and C, respectively.

**Figure S23:**
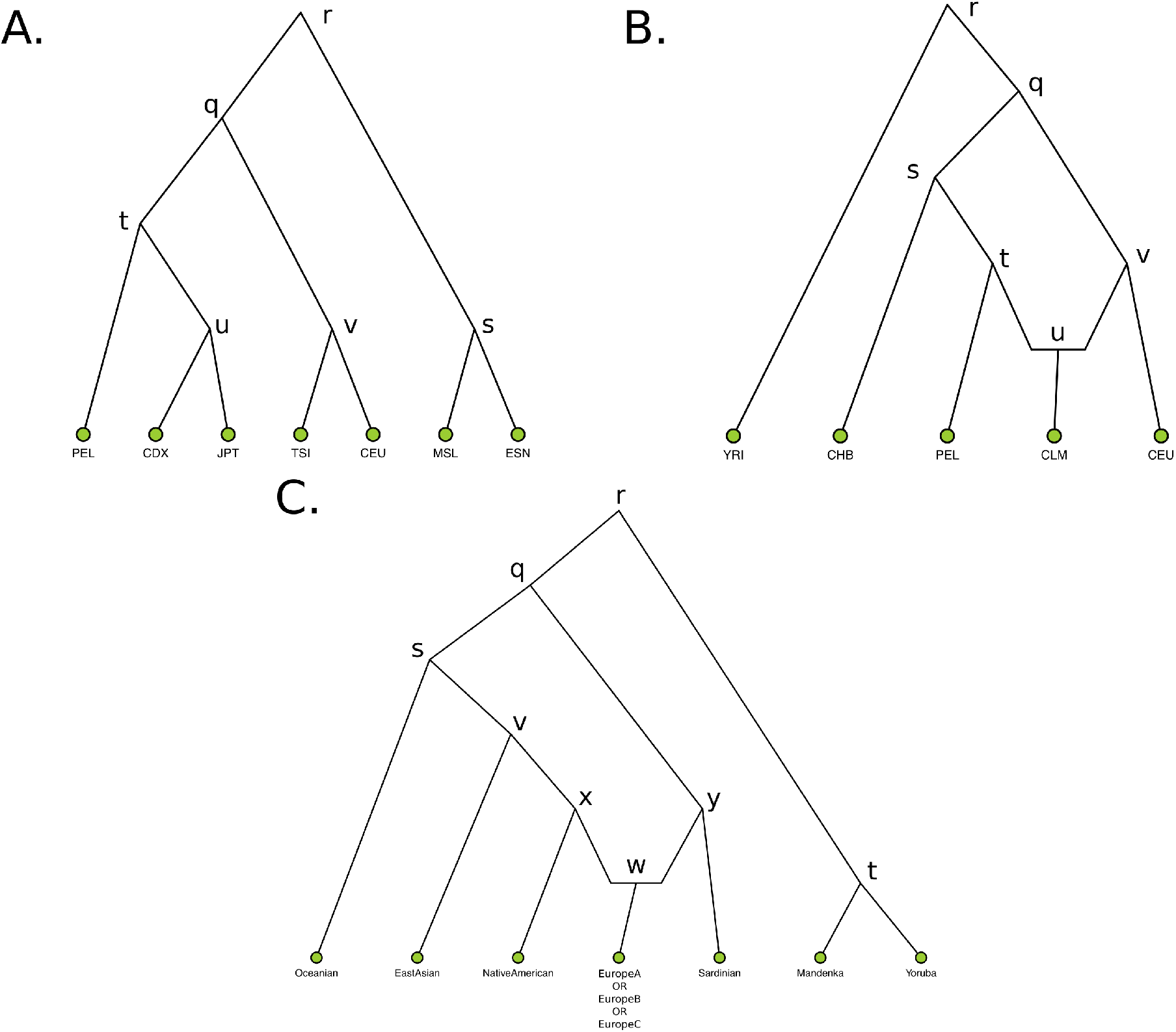
A) A 7-leaf population tree containing the following population panels from the 1000 Genomes Project: Nigerian Esan (ESN), Sierra Leone Mende (MSL), Northern Europeans from Utah (CEU), Southern Europeans from Tuscany (TSI), Dai Chinese (CDX), Japanese (JPT) and Peruvians (PEL). B) A 5-leaf admixture graph containing the following panels from the 1000 Genomes project: Yoruba (YRI), Colombians (CLM), CEU, CHB and PEL. C) A 7-leaf admixture graph containing panels (and combinations of panels) from the imputed Lazaridis et al. (2014) dataset. EastAsian = [Cambodian, Mongola, Xibo, Daur, Hezhen, Oroqen, Naxi, Yi, Japanese, Han_NChina, Lahu, Miao, She, Han, Tujia, Dai]. NativeAmerican = [Maya, Pima, Surui, Karitiana, Colombian]. Oceanian = [Papuan, Australian]. We tested three graphs with three different combinations of European populations, with different amounts of EEF ancestry: EuropeA (low EEF) = [Estonian, Lithuanian, Scottish, Icelandic, Norwegian, Orcadian, Czech, English]. EuropeB (medium EEF) = [Hungarian, Croatian, French, Basque, Spanish_North, French_South]. EuropeC (high EEF) = [Bulgarian, Bergamo, Tuscan, Albanian, Greek, Spanish].

**Figure S24:**
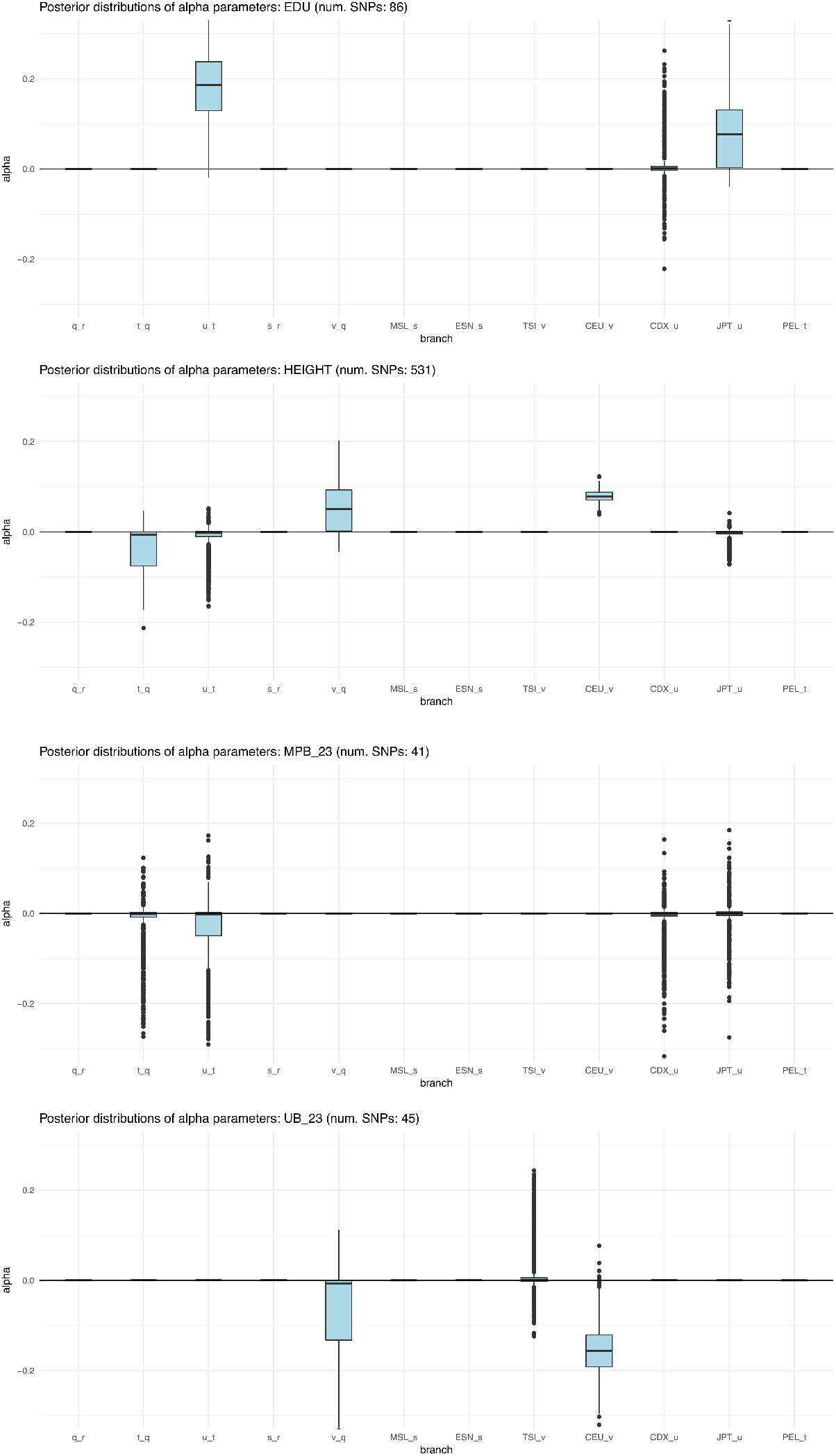
Box plots of posterior distributions for *α* parameters, for trait-associated variants with significant evidence for selection in a 7-leaf population tree using the 1000 Genomes data (Figure S23.A). Parameters with flat distributions were discarded a priori using the *Q_B_* statistic. The lower, middle and upper hinges denote the 25th, 50th and 75th percentiles, respectively. The upper whisker extends to the highest value that is within 1.5 * IQR of the upper hinge, where IQR is the inter-quartile range. The lower whisker extends to the lowest value within 1.5 * IQR of the lower hinge. Data beyond the whiskers are plotted as points. EDU = educational attainment. MPB_23 = self-reported male-pattern baldness from 23andMe. UB_23 = self-reported unibrow from 23andMe.

**Figure S25:**
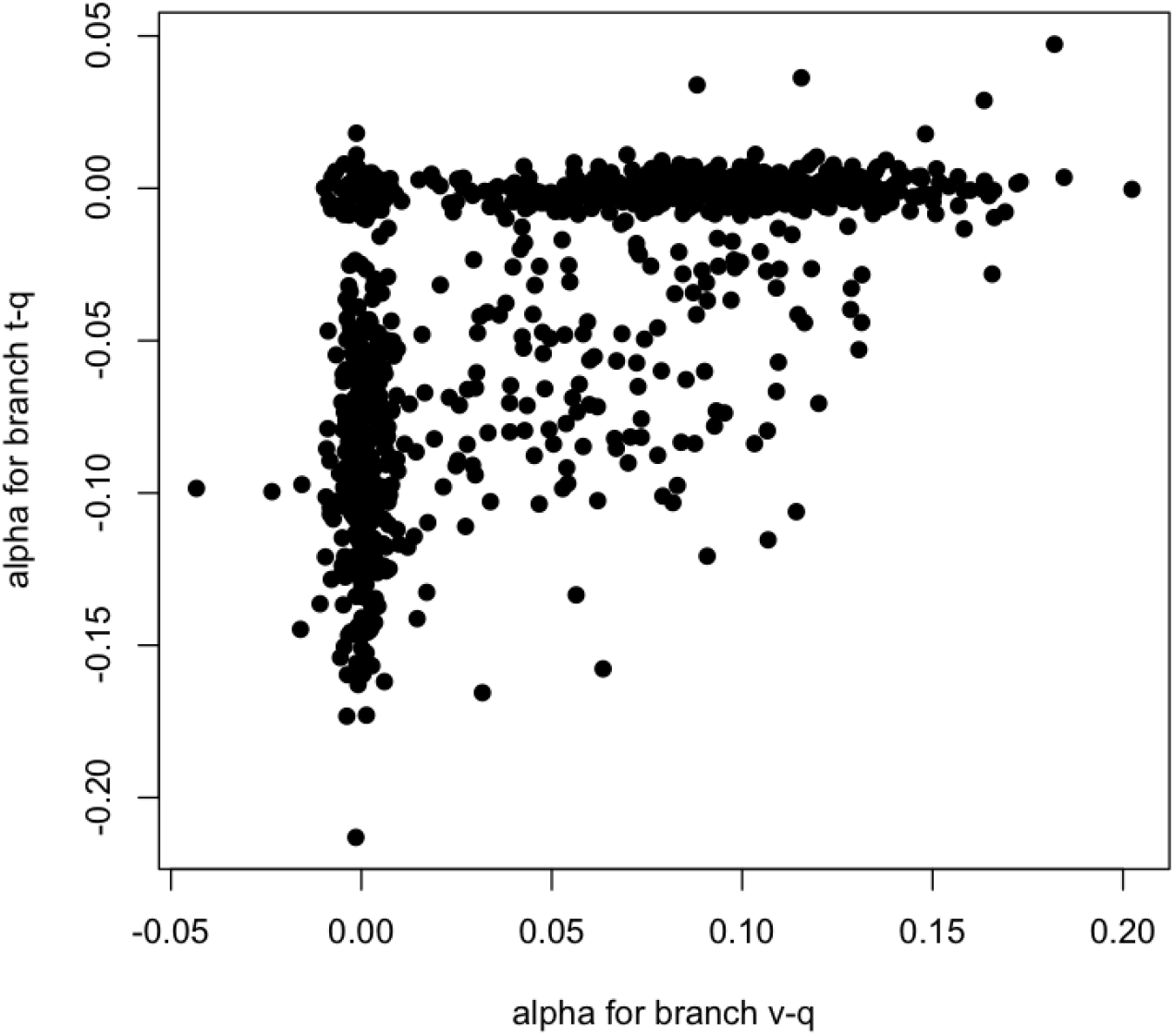
We plotted the *α* values of two branches for all posterior samples of the MCMC run for variants associated with height in the 7-leaf population tree composed of 1000 Genomes panels. For each MCMC sample, the x-axis corresponds to the *α* parameter in the ancestral European branch, while the y-axis corresponds to the *α* parameter in the ancestral East Asian / Native American branch.

**Figure S26:**
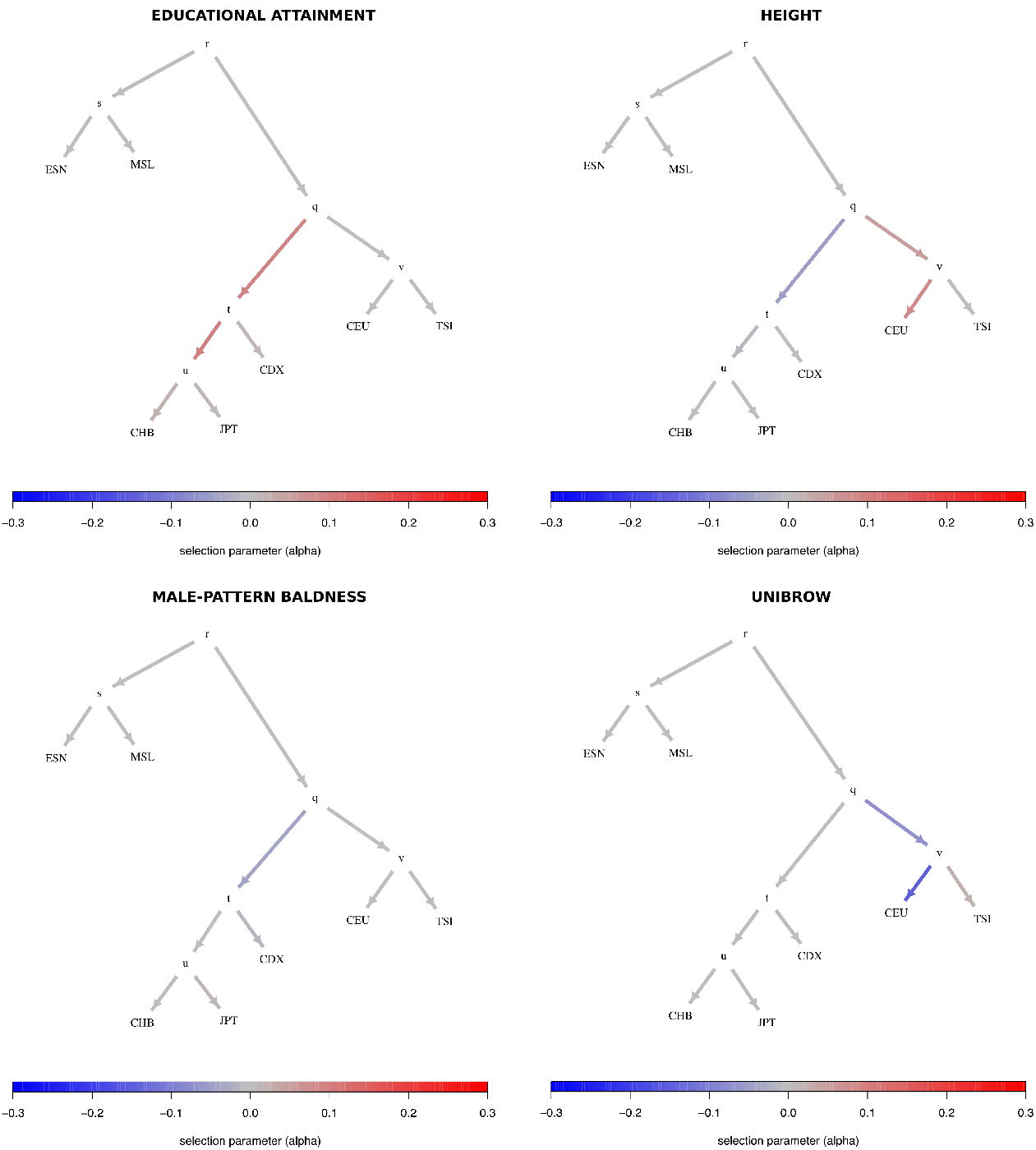
Poly-graphs for trait-associated variants that show significant evidence for polygenic adaptation in a 7-leaf tree built using 1000 Genomes allele frequency data. ESN = Nigerian Esan; MSL = Sierra Leone Mende; CEU = Northern Europeans from Utah; TSI = Southern Europeans from Tuscany; CDX = Dai Chinese; JPT = Japanese; CHB = Han Chinese from Beijing.

**Figure S27:**
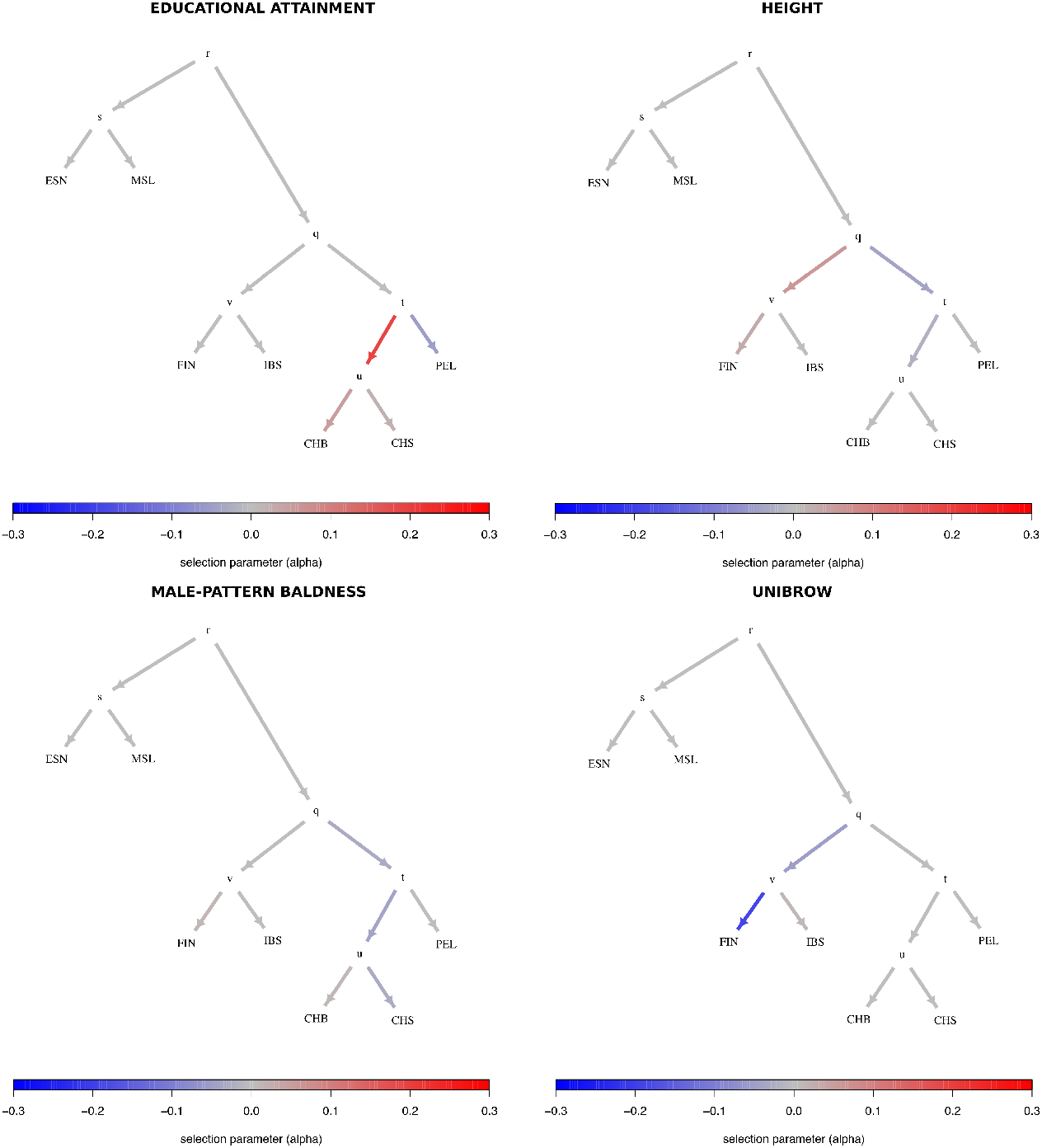
Poly-graphs for trait-associated variants that show significant evidence for polygenic adaptation in a 7-leaf tree built using 1000 Genomes allele frequency data. ESN = Nigerian Esan; MSL = Sierra Leone Mende; CHB = Han Chinese from Beijing; CHS = Southern Han Chinese; FIN = Finnish; IBS = Iberians from Spain; PEL = Peruvians from Lima.

**Figure S28:**
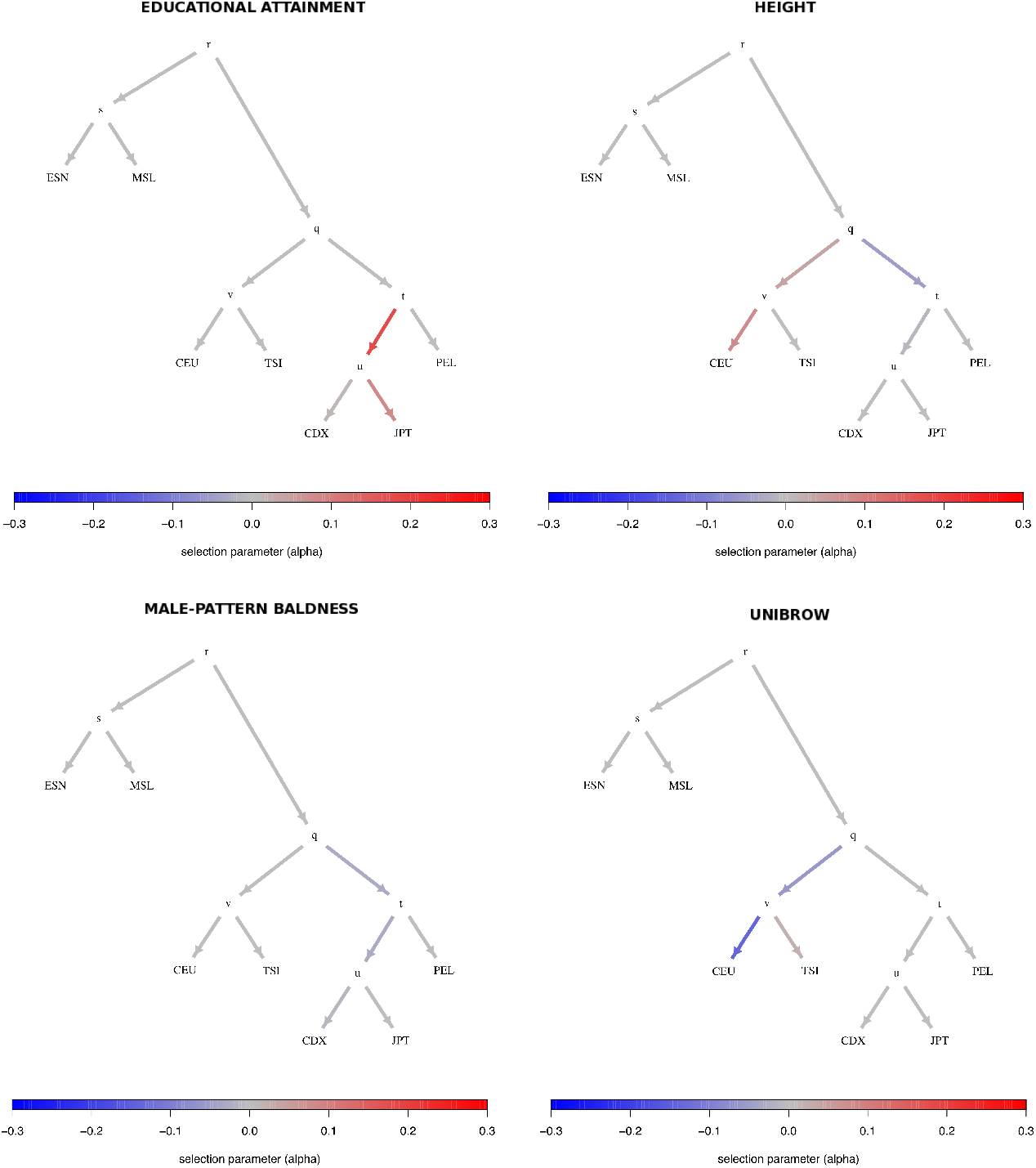
Poly-graphs for trait-associated variants that show significant evidence for polygenic adaptation in the 7-leaf tree built using 1000 Genomes allele frequency data, using a Beta(2,2) root allele frequency prior in the MCMC. ESN = Nigerian Esan; MSL = Sierra Leone Mende; CEU = Northern Europeans from Utah; TSI = Southern Europeans from Tuscany; CDX = Dai Chinese; JPT = Japanese; PEL = Peruvians.

**Figure S29:**
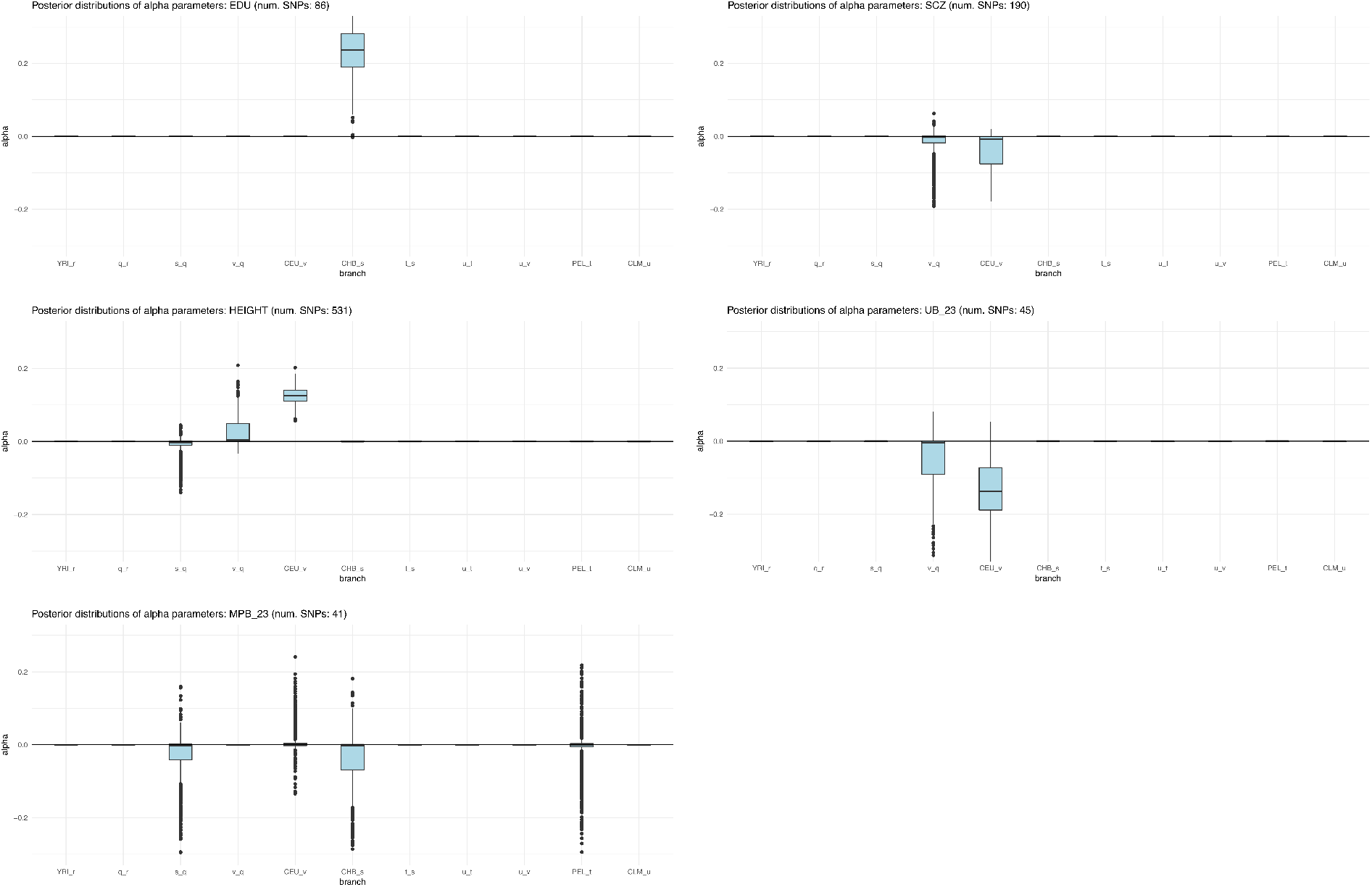
Box plots of posterior distributions for *α* parameters, for trait-associated variants with significant evidence for selection in a 5-leaf admixture graph using the 1000 Genomes data (Figure S23.B). Parameters with flat distributions were discarded a priori using the *Q_B_* statistic. The lower, middle and upper hinges denote the 25th, 50th and 75th percentiles, respectively. The upper whisker extends to the highest value that is within 1.5 * IQR of the upper hinge, where IQR is the inter-quartile range. The lower whisker extends to the lowest value within 1.5 * IQR of the lower hinge. Data beyond the whiskers are plotted as points. EDU = educational attainment. MPB_23 = self-reported male-pattern baldness from 23andMe. UB_23 = self-reported unibrow from 23andMe. SCZ = schizophrenia.

**Figure S30:**
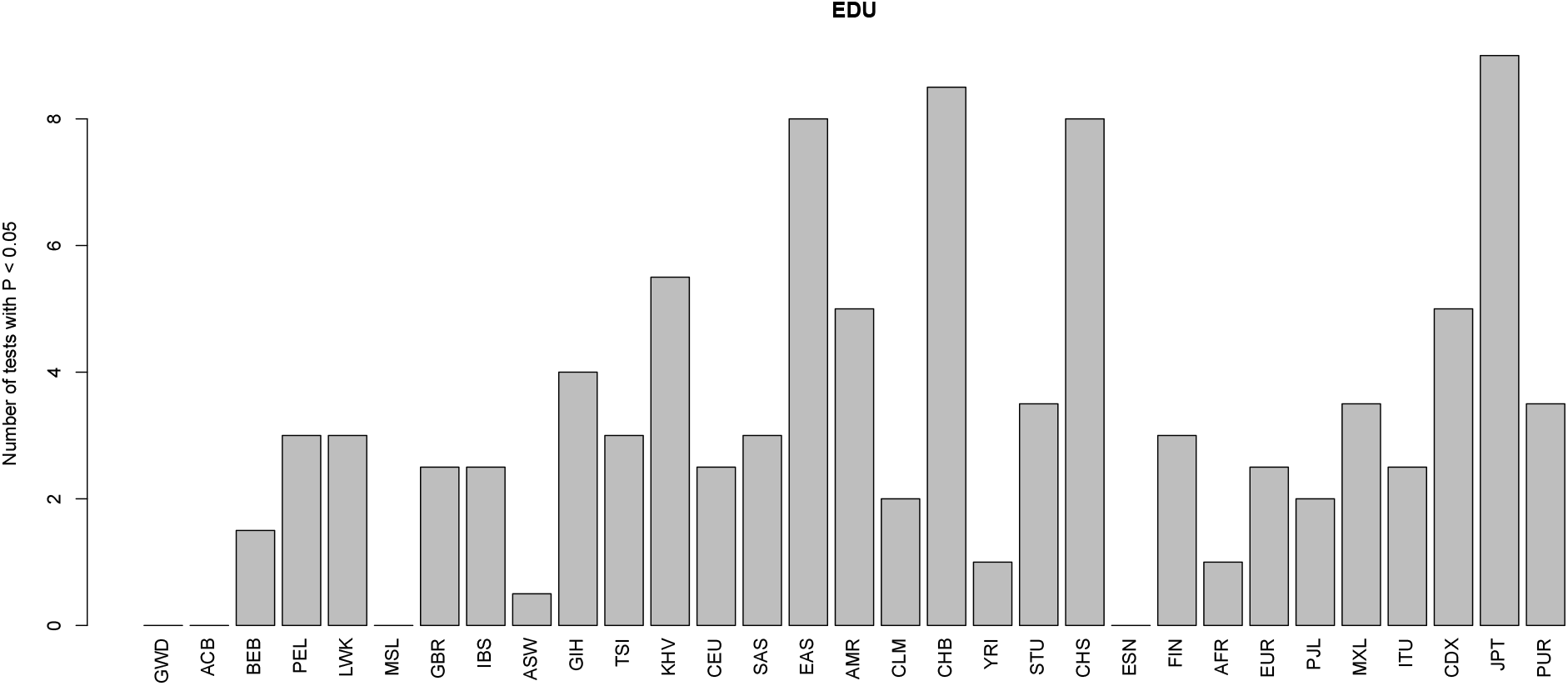
This barplot shows, for each panel, the number of two-tailed binomial tests for systematic allele frequency differences in the sign of the effect size estimate of variants associated with educational attainment with P-values < 0.05, which involve that panel as a member of the pair. We tested all panels and continental super-panels from the 1000 Genomes dataset.

**Figure S31:**
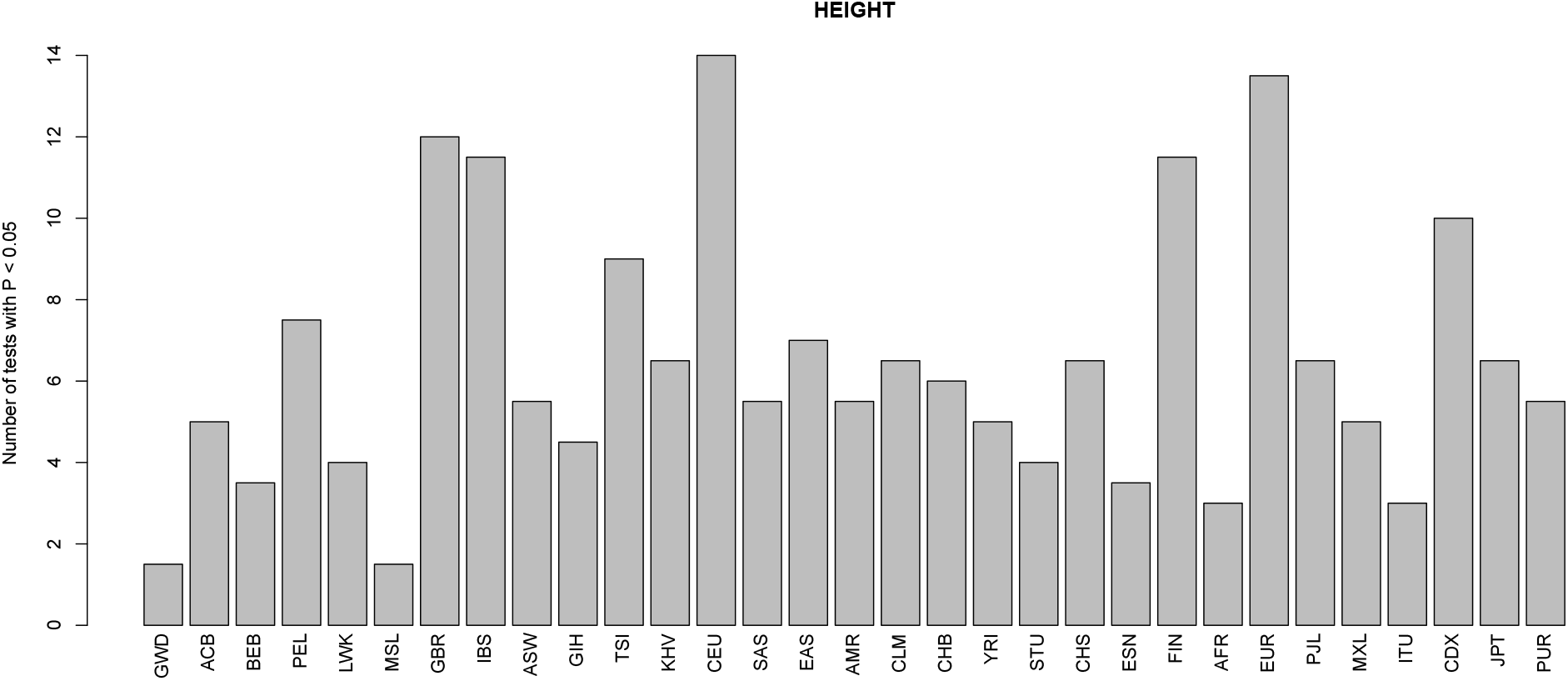
This barplot shows, for each panel, the number of two-tailed binomial tests for systematic allele frequency differences in the sign of the effect size estimate of variants associated with height with P-values < 0.05, which involve that panel as a member of the pair. We tested all panels and continental super-panels from the 1000 Genomes dataset.

**Figure S32:**
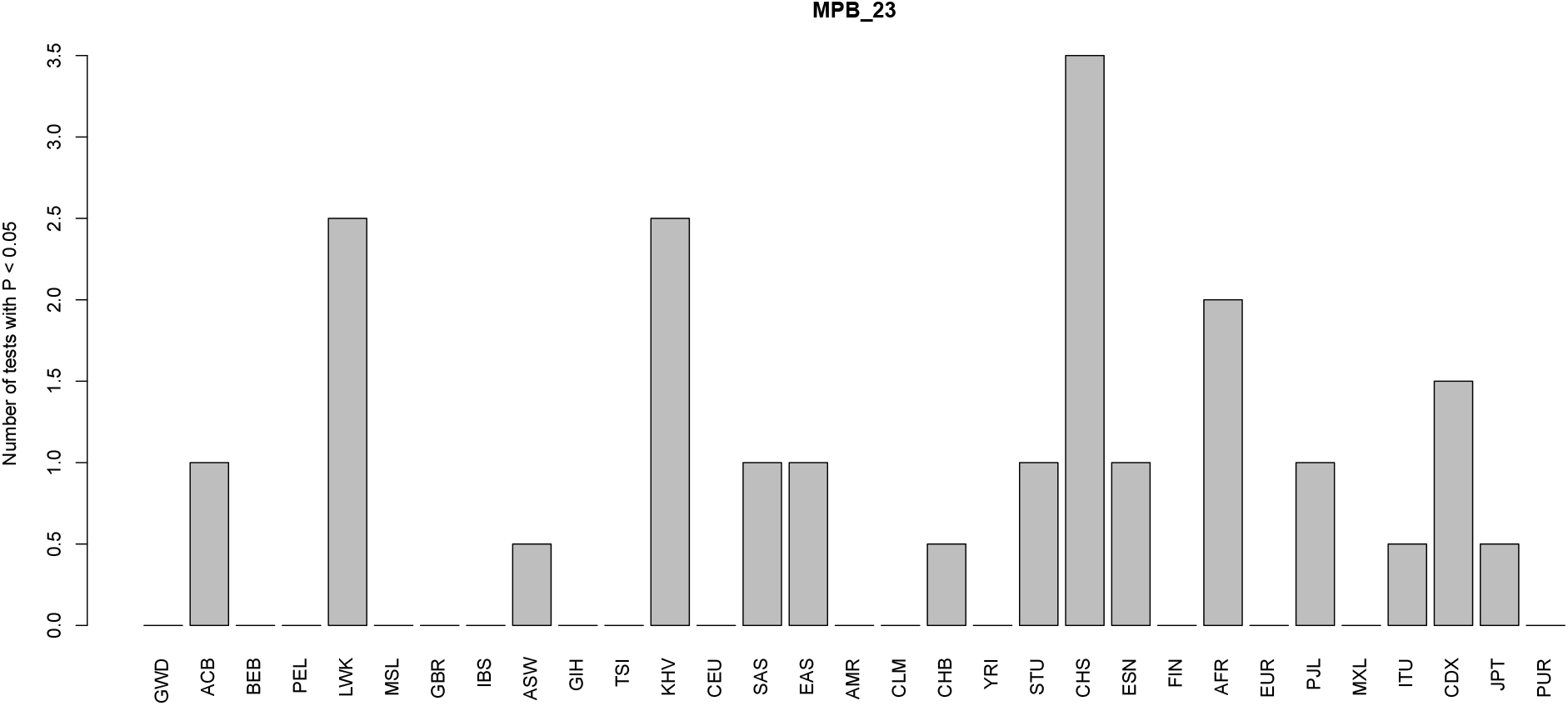
This barplot shows, for each panel, the number of two-tailed binomial tests for systematic allele frequency differences in the sign of the effect size estimate of variants associated with self-reported male-pattern baldness with P-values < 0.05, which involve that panel as a member of the pair. We tested all panels and continental super-panels from the 1000 Genomes dataset.

**Figure S33:**
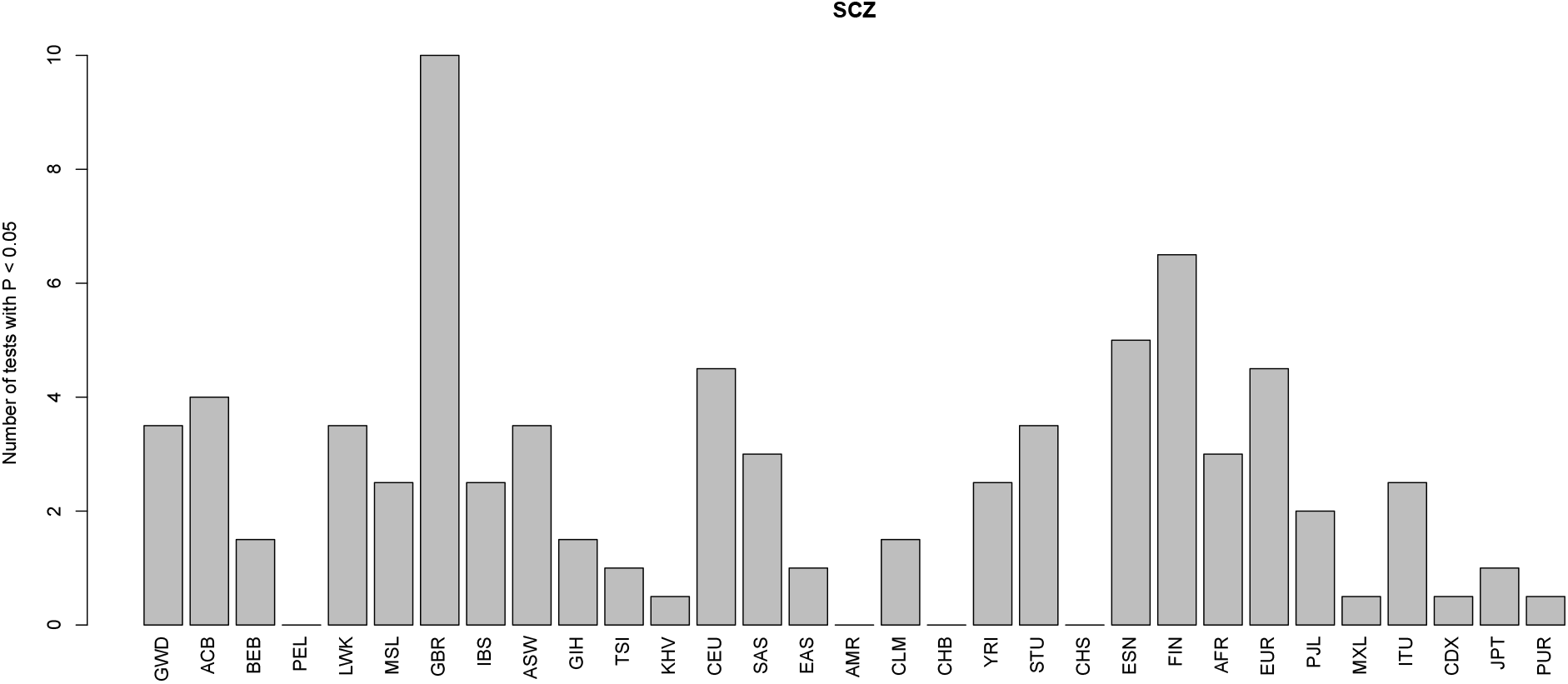
This barplot shows, for each panel, the number of two-tailed binomial tests for systematic allele frequency differences in the sign of the effect size estimate of variants associated with schizophrenia with P-values < 0.05, which involve that panel as a member of the pair. We tested all panels and continental super-panels from the 1000 Genomes dataset.

**Figure S34:**
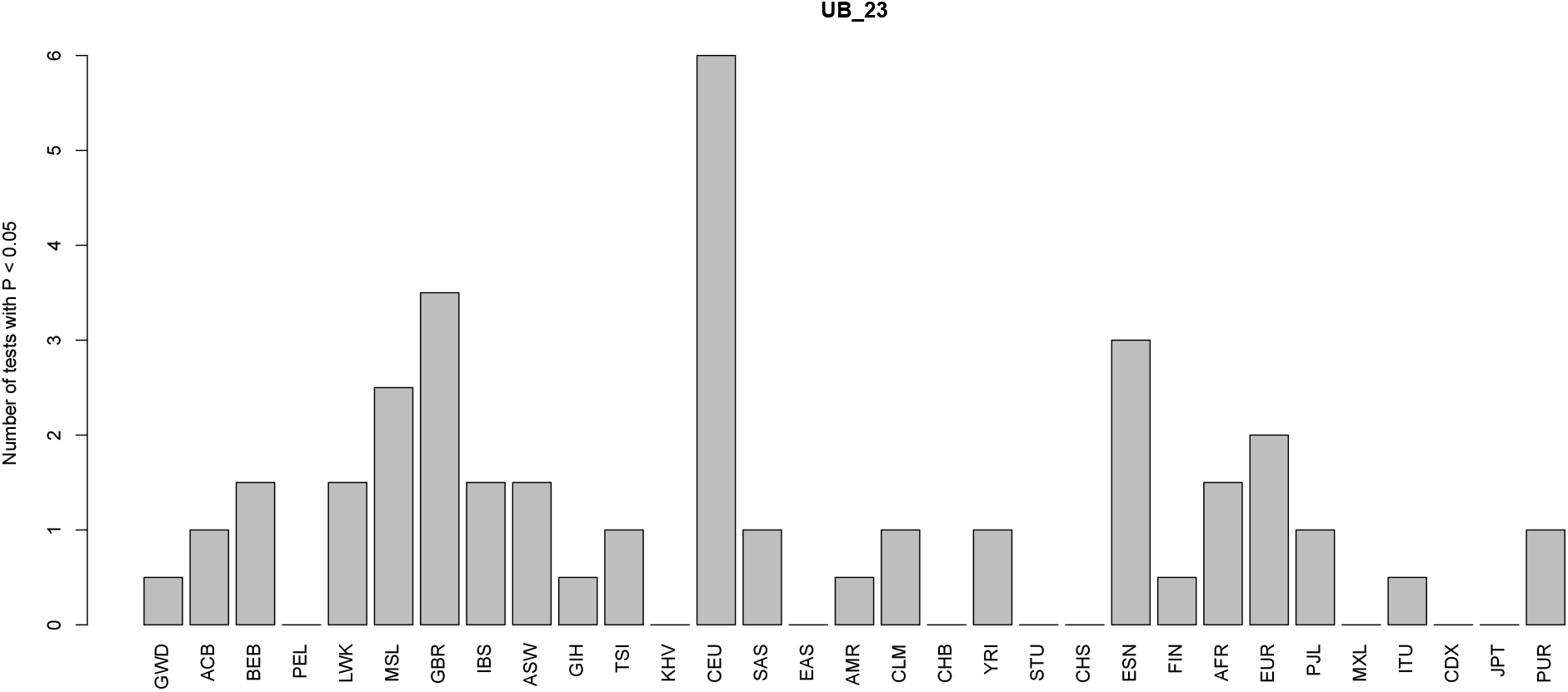
This barplot shows, for each panel, the number of two-tailed binomial tests for systematic allele frequency differences in the sign of the effect size estimate of variants associated with self-reported unibrow with P-values < 0.05, which involve that panel as a member of the pair. We tested all panels and continental super-panels from the 1000 Genomes dataset.

**Figure S35:**
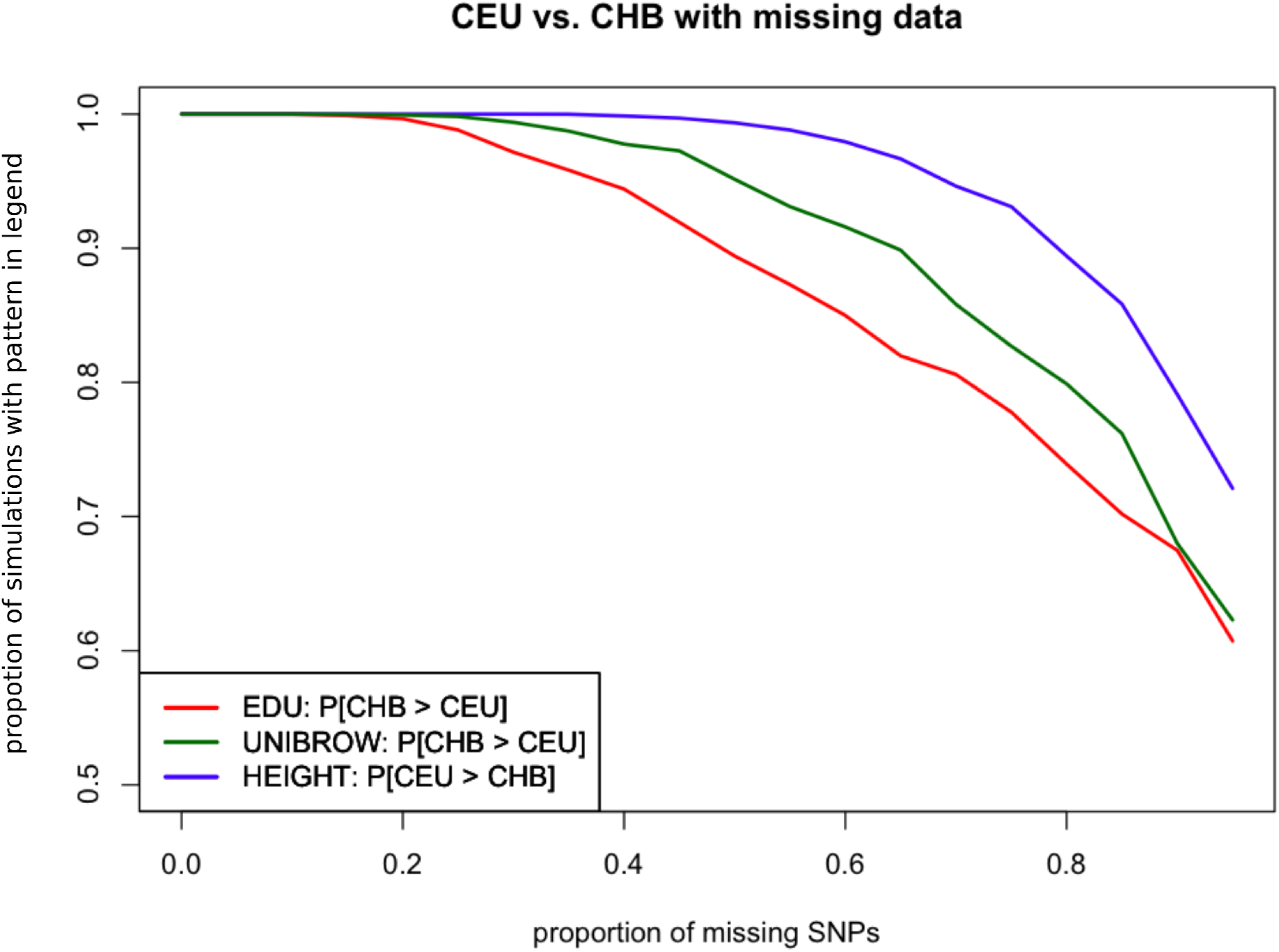
We simulated different proportions of missing trait-associated SNPs (x-axis). We then assessed - for each proportion - how often we could recreate the inequality relationship observed between the polygenic scores for CHB and CEU built using the trait-associated SNPs of the three traits with strongest evidence for polygenic adaptation: height, educational attainment and self-reported unibrow. The observed inequality relationship for educational attainment and unibrow is *Poly_CHB_* > *Poly_CEU_*, where *Poly_X_* is the polygenic score for panel X. The observed inequality relationship for height is the reverse: *Poly_CEU_* > *Poly_CHB_*. We used 10,000 simulations for each missing data scenario.

**Figure S36:**
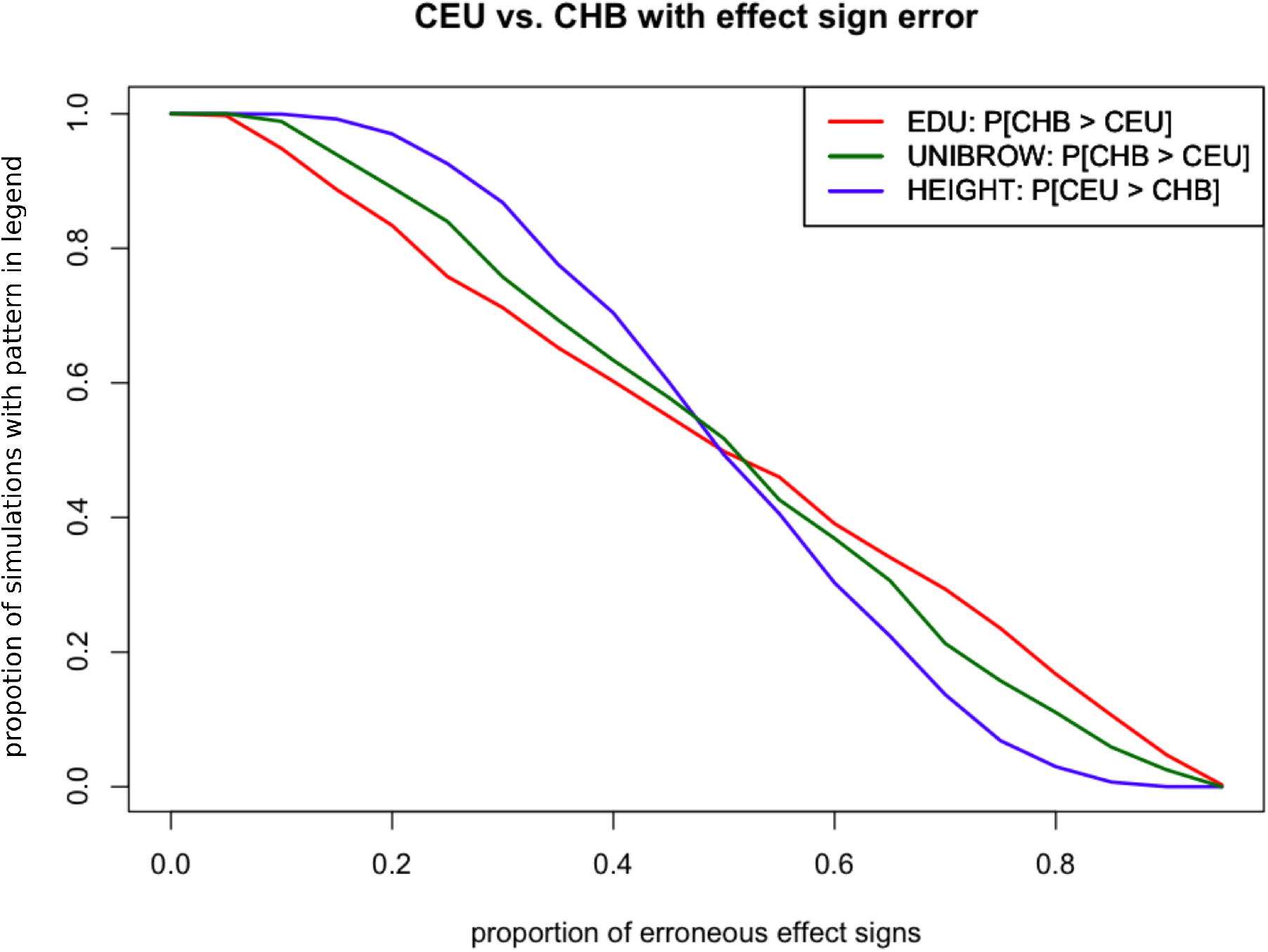
We simulated different proportions of incorrectly assigned signs for the effect size estimates of trait-associated SNPs. We then assessed - for each proportion - how often we could recreate the inequality relationship observed between the polygenic scores for CHB and CEU built using the trait-associated SNPs of the three traits with strongest evidence for polygenic adaptation: height, educational attainment and self-reported unibrow. The observed inequality relationship for educational attainment and unibrow is *Poly_CHB_* > *Poly_CEU_*, where *Poly_X_* is the polygenic score for panel X. The observed inequality relationship for height is the reverse: *Poly_CEU_* > *Poly_CHB_*. We used 10,000 simulations for each sign misassignment scenario.

**Figure S37:**
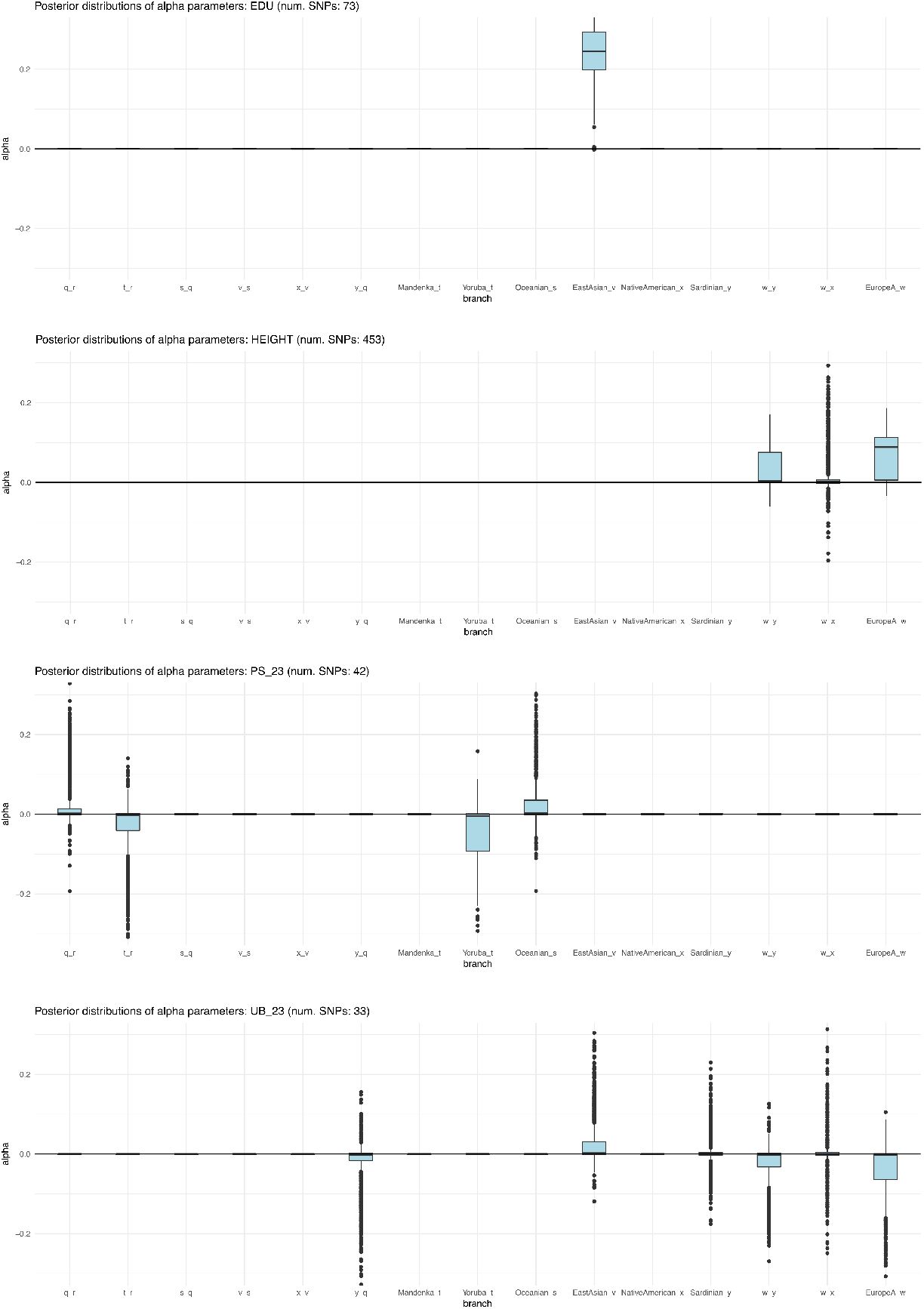
Box plots of posterior distribution for *α* parameters, for trait-associated variants with significant evidence for selection in a 7-leaf population graph, using the Lazaridis et al. (2014) dataset and including the set of European populations with low EEF ancestry (“EuropeA”). Parameters with flat distributions were discarded a priori using the *Q_B_* statistic. The lower, middle and upper hinges denote the 25th, 50th and 75th percentiles, respectively. The upper whisker extends to the highest value that is within 1.5 * IQR of the upper hinge, where IQR is the inter-quartile range. The lower whisker extends to the lowest value within 1.5 * IQR of the lower hinge. Data beyond the whiskers are plotted as points. EDU = educational attainment. PS_23 = self-reported photic sneeze reflex from 23andMe. UB_23 = self-reported unibrow from 23andMe.

**Figure S38:**
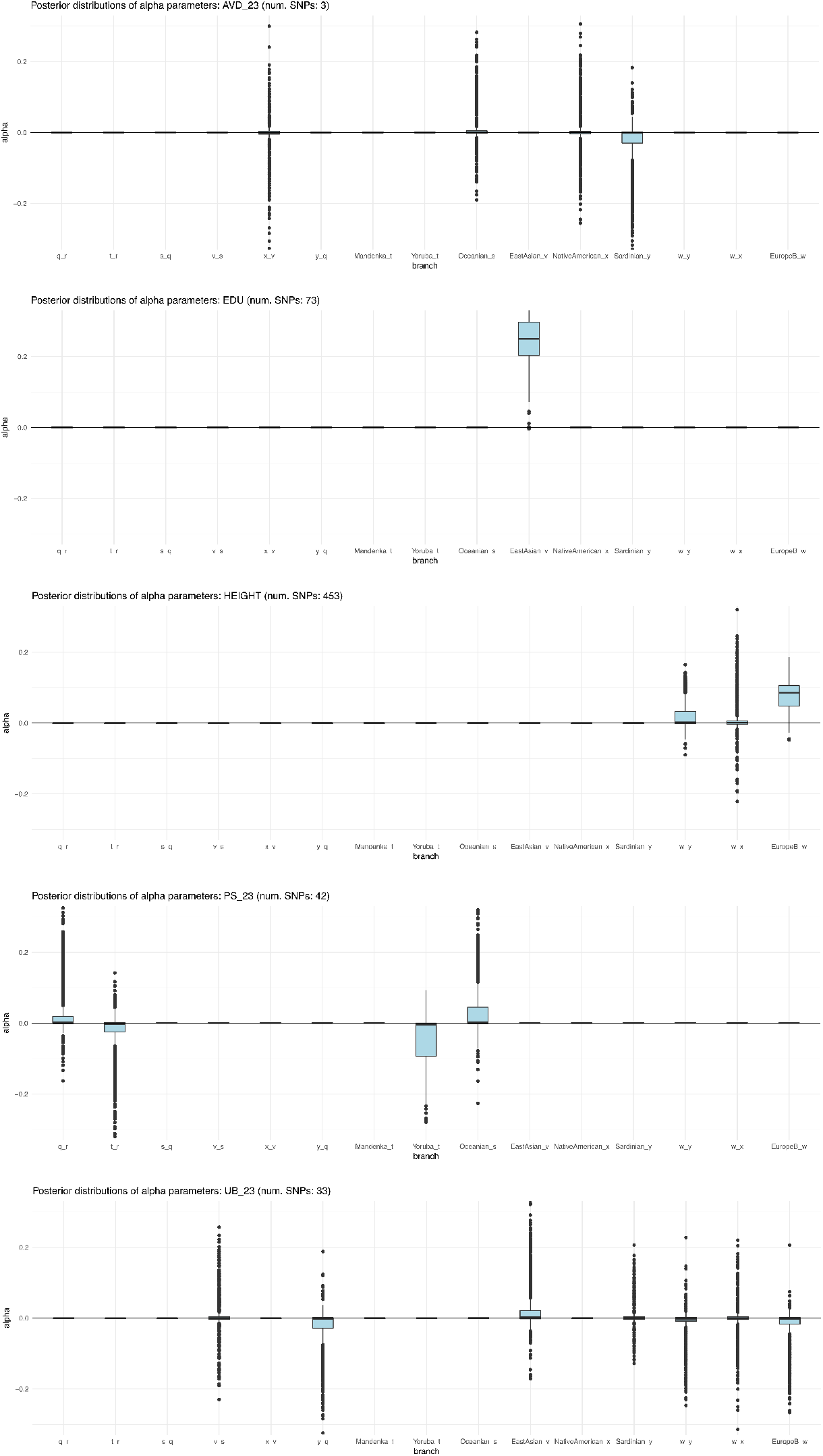
Box plots of posterior distribution for *α* parameters, for trait-associated variants with significant evidence for selection in a 7-leaf population graph, using the Lazaridis et al. (2014) dataset and including the set of European populations with medium EEF ancestry (“EuropeB”). Parameters with flat distributions were discarded a priori using the *Q_B_* statistic. The lower, middle and upper hinges denote the 25th, 50th and 75th percentiles, respectively. The upper whisker extends to the highest value that is within 1.5 * IQR of the upper hinge, where IQR is the inter-quartile range. The lower whisker extends to the lowest value within 1.5 * IQR of the lower hinge. Data beyond the whiskers are plotted as points. AVD_23 = self-reported age at voice drop. EDU = educational attainment. PS_23 = self-reported photic sneeze reflex from 23andMe. UB_23 = self-reported unibrow from 23andMe.

**Figure S39:**
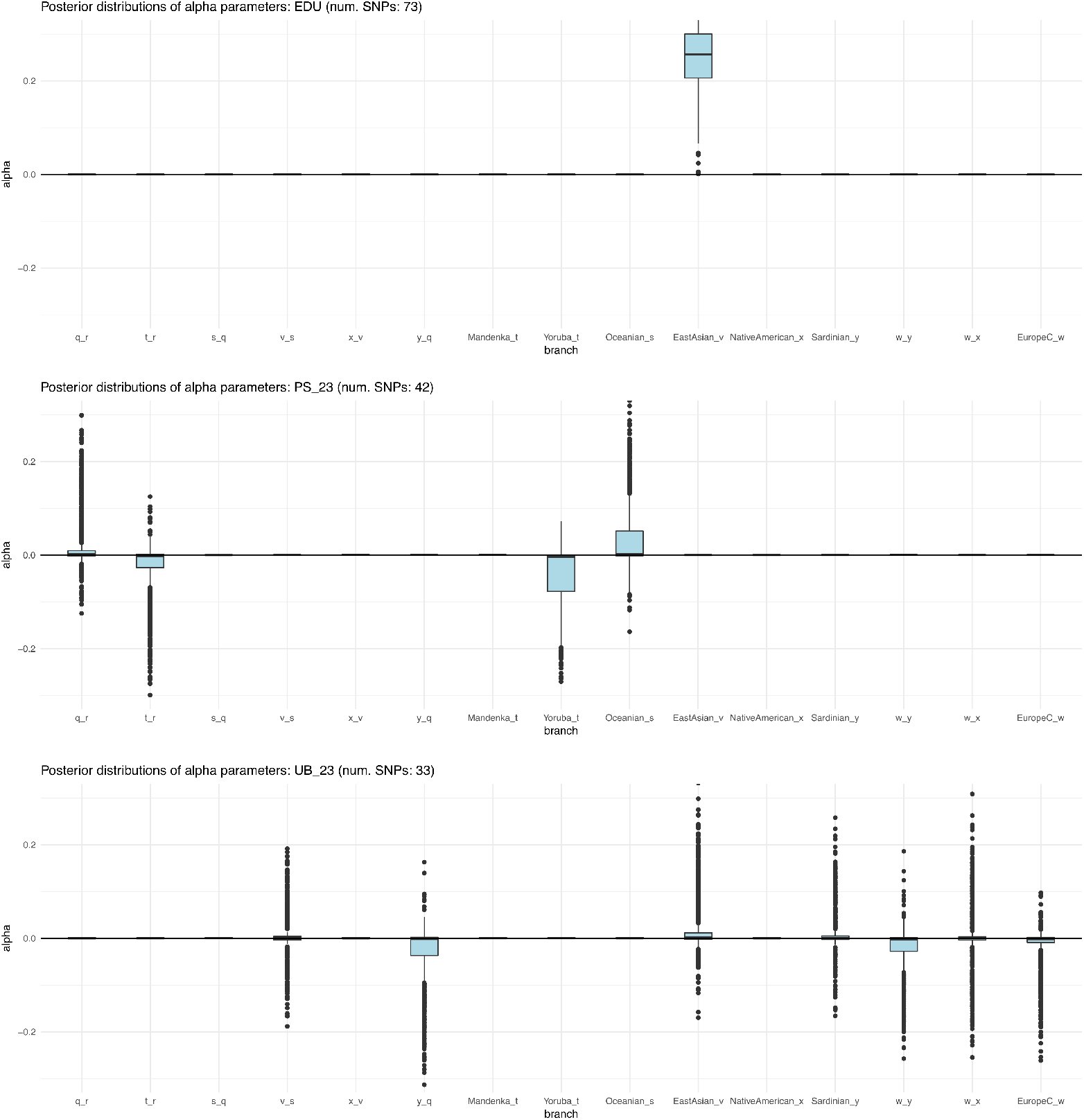
Box plots of posterior distribution for *α* parameters, for trait-associated variants with significant evidence for selection in a 7-leaf population graph, using the Lazaridis et al. (2014) dataset and including the set of European populations with high EEF ancestry (“EuropeC”). Parameters with flat distributions were discarded a priori using the *Q_B_* statistic. The lower, middle and upper hinges denote the 25th, 50th and 75th percentiles, respectively. The upper whisker extends to the highest value that is within 1.5 * IQR of the upper hinge, where IQR is the inter-quartile range. The lower whisker extends to the lowest value within 1.5 * IQR of the lower hinge. Data beyond the whiskers are plotted as points. EDU = educational attainment. PS_23 = self-reported photic sneeze reflex from 23andMe. UB_23 = self-reported unibrow from 23andMe.

**Figure S40:**
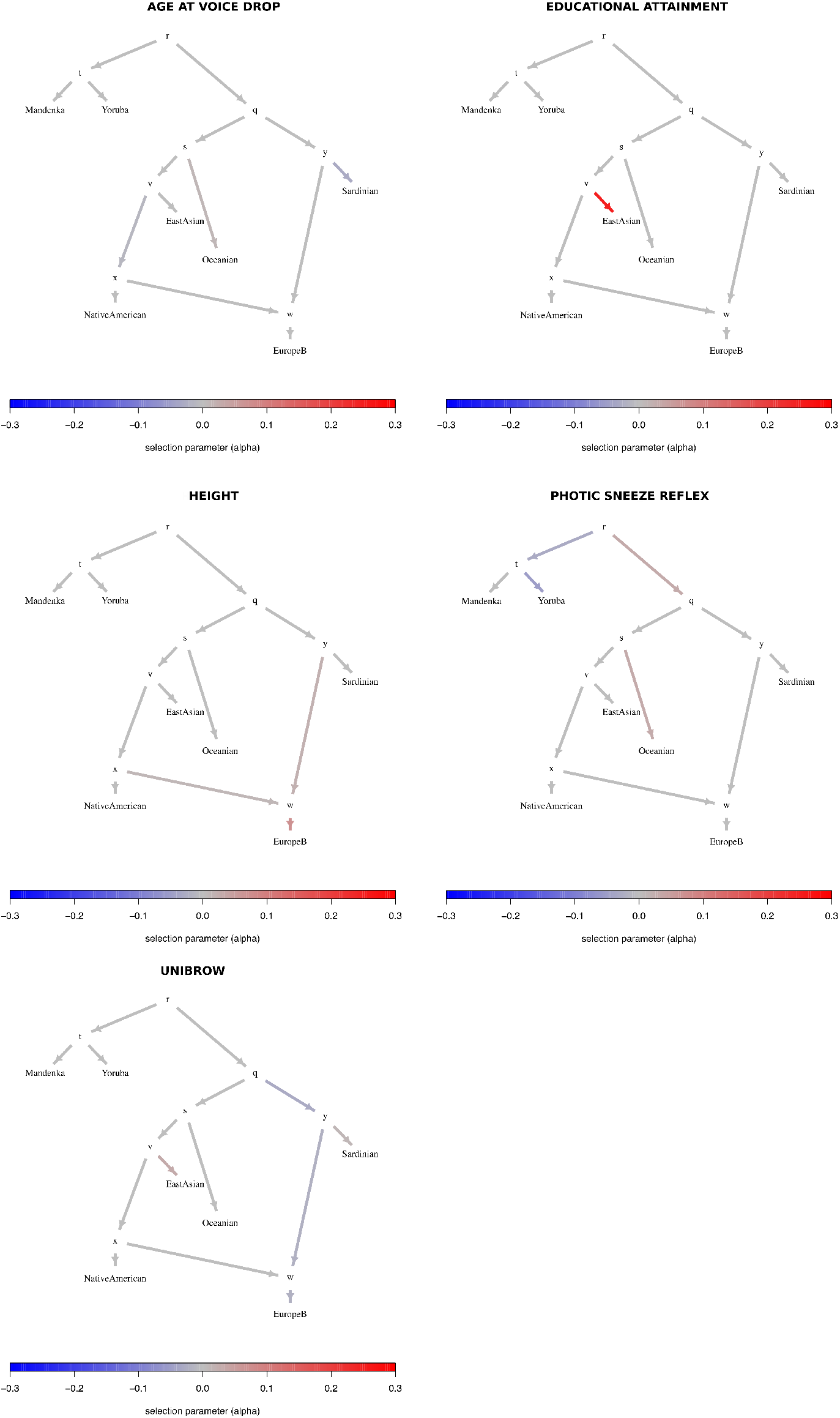
Graphs for trait-associated variants that show significant evidence for polygenic adaptation in the 7-leaf admixture graph built using the Lazaridis et al. (2014) dataset and including the set of European populations with medium EEF ancestry (“EuropeB”).

**Figure S41:**
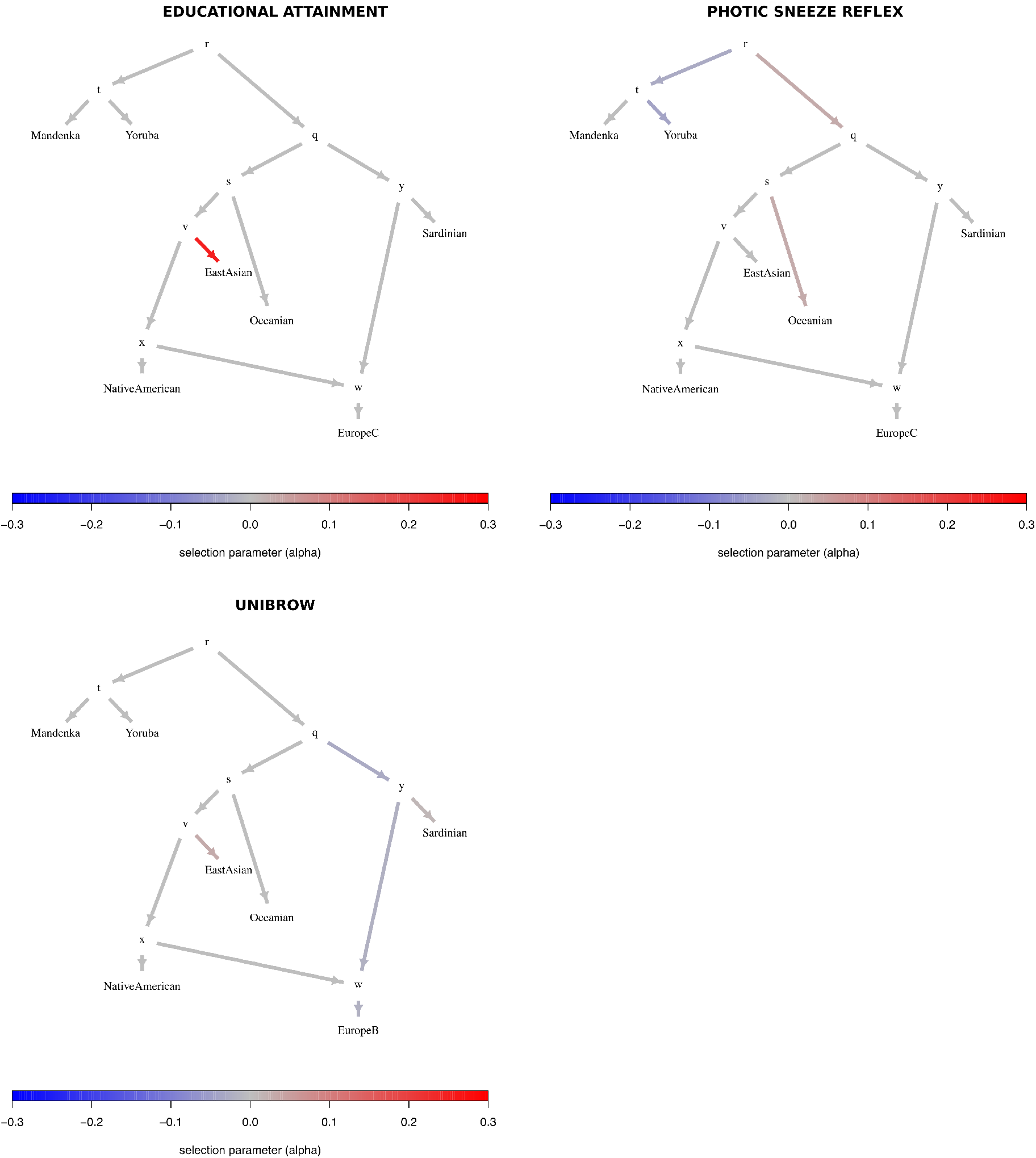
Graphs for trait-associated variants that show significant evidence for polygenic adaptation in the 7-leaf admixture graph built using the Lazaridis et al. (2014) dataset and including the set of European populations with high EEF ancestry (“EuropeC”).

**Figure S42:**
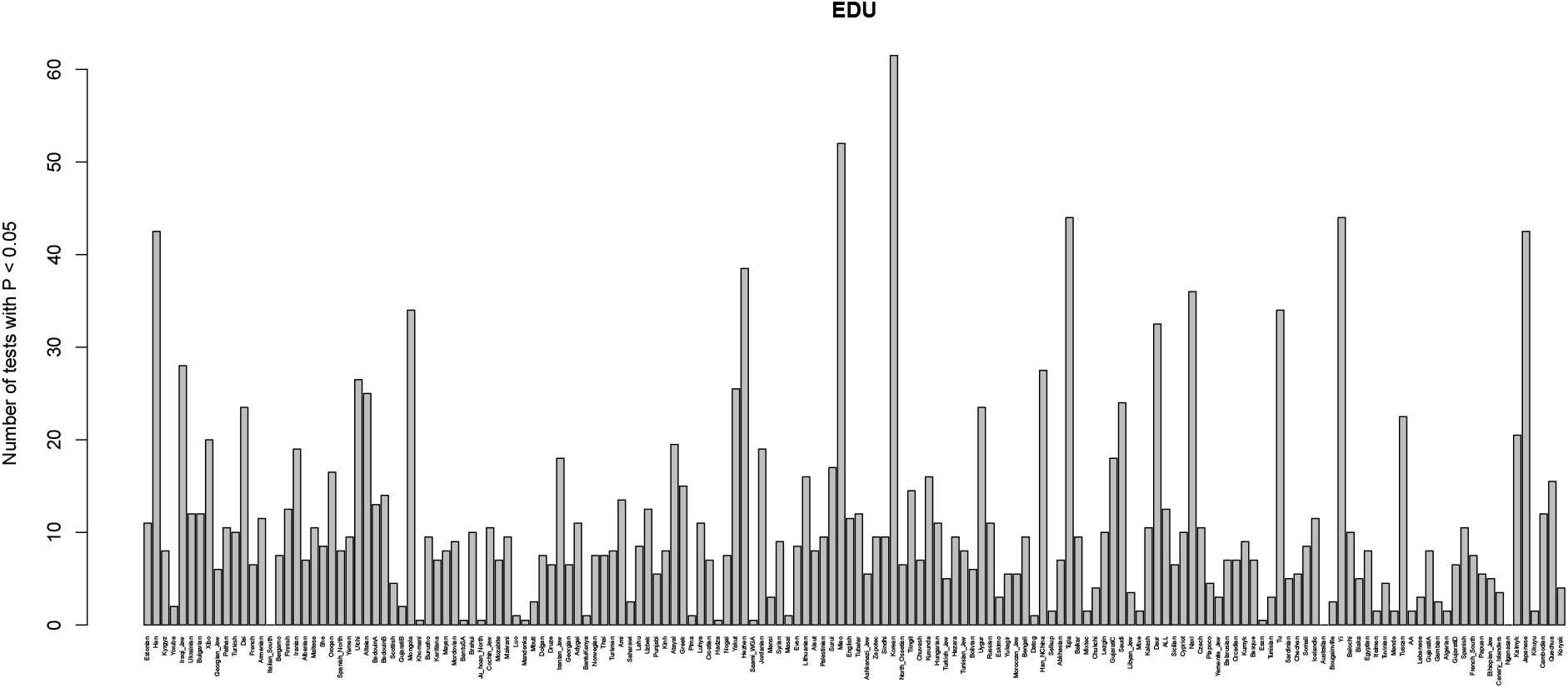
This barplot shows, for each panel, the number of two-tailed binomial tests for systematic allele frequency differences in the sign of the effect size estimate of variants associated with educational attainment with P-values < 0.05, which involve that panel as a member of the pair. We tested all panels from the Lazaridis et al. (2014) dataset.

**Figure S43:**
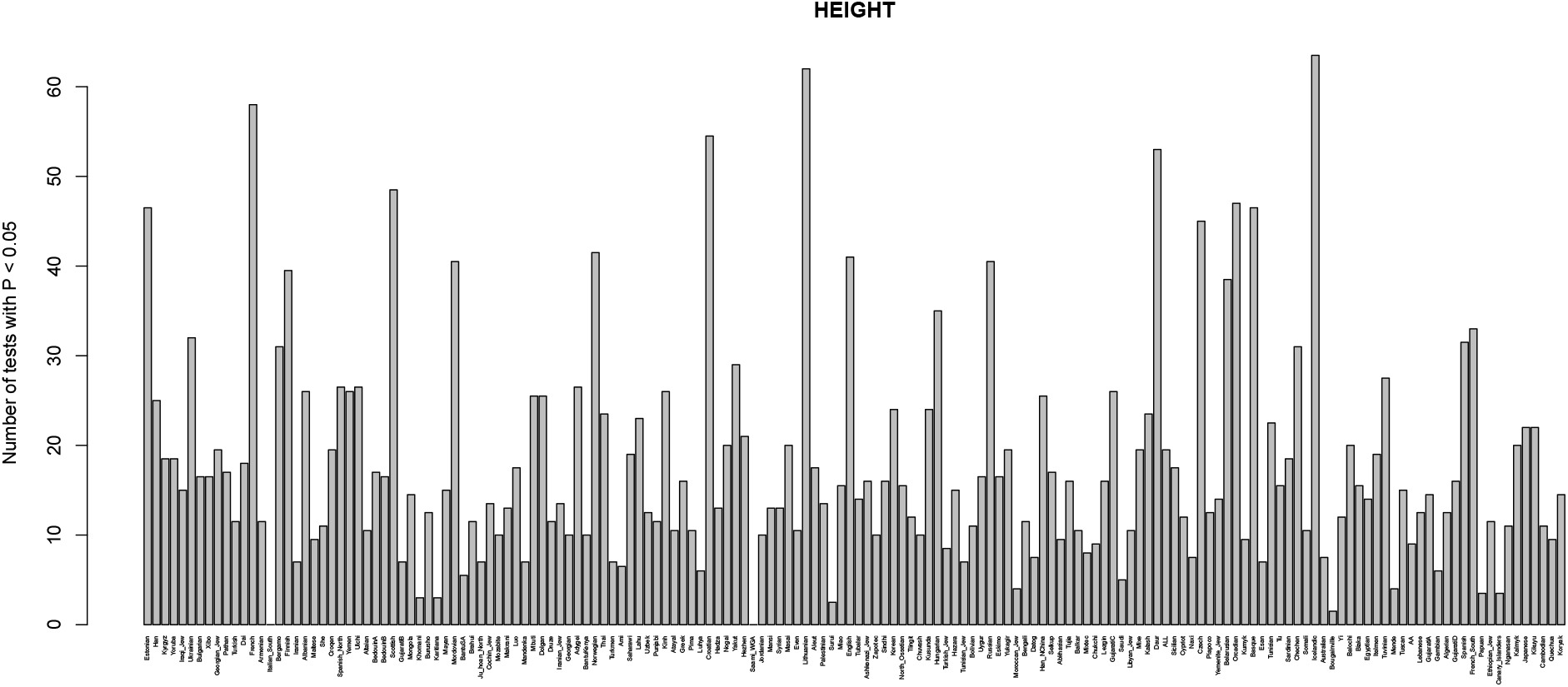
This barplot shows, for each panel, the number of two-tailed binomial tests for systematic allele frequency differences in the sign of the effect size estimate of variants associated with height with P-values < 0.05, which involve that panel as a member of the pair. We tested all panels from the Lazaridis et al. (2014) dataset.

**Figure S44:**
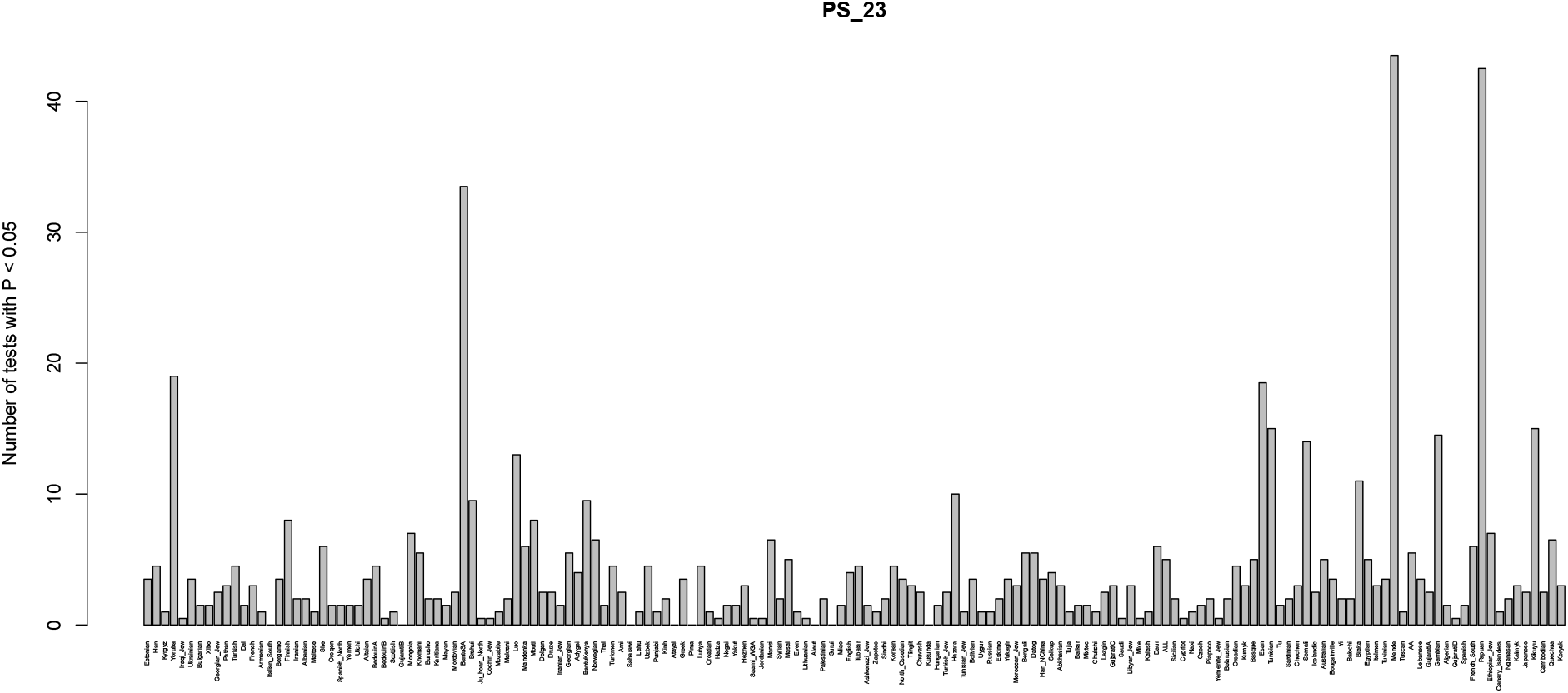
This barplot shows, for each panel, the number of two-tailed binomial tests for systematic allele frequency differences in the sign of the effect size estimate of variants associated with self-reported photic sneeze reflex with P-values < 0.05, which involve that panel as a member of the pair. We tested all panels from the Lazaridis et al. (2014) dataset.

**Figure S45:**
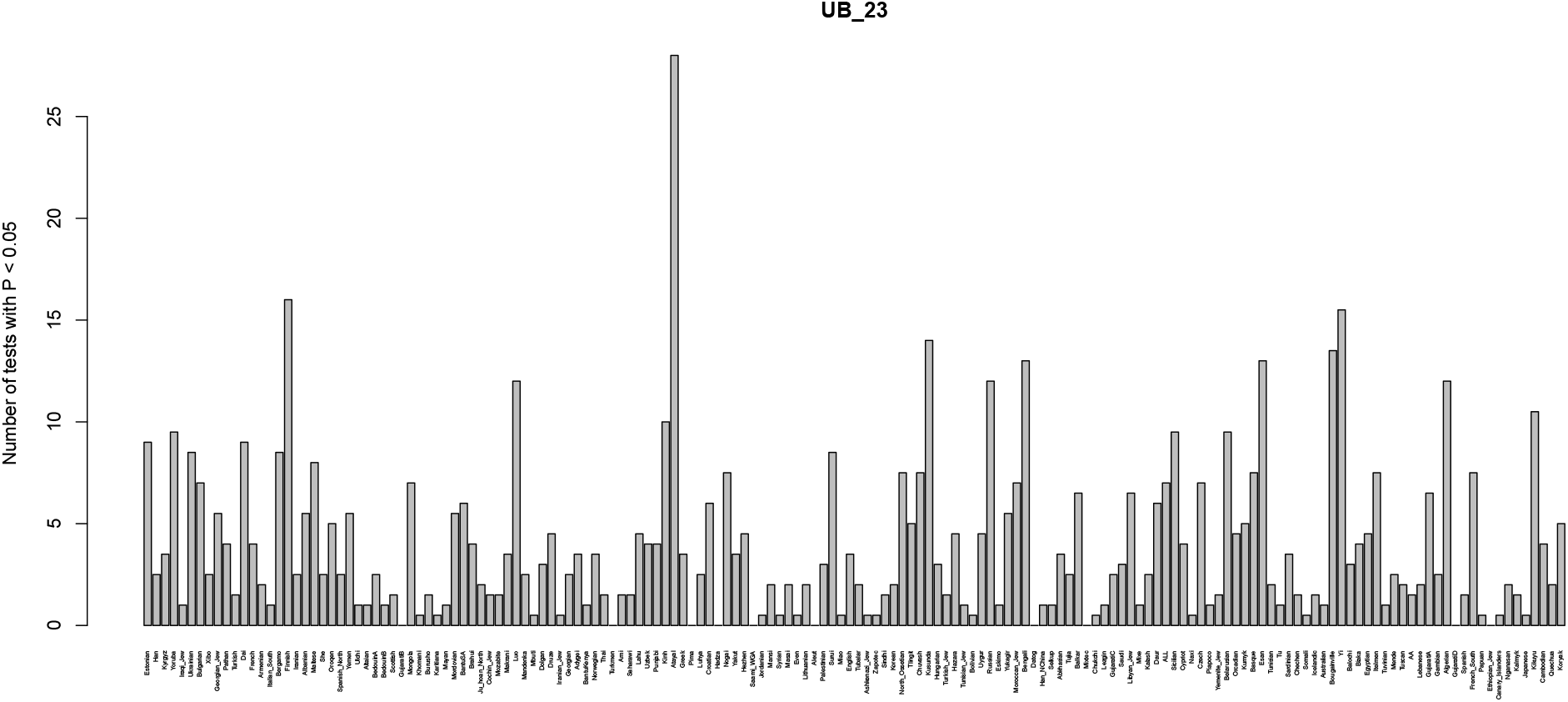
This barplot shows, for each panel, the number of two-tailed binomial tests for systematic allele frequency differences in the sign of the effect size estimate of variants associated with self-reported unibrow with P-values < 0.05, which involve that panel as a member of the pair. We tested all panels from the Lazaridis et al. (2014) dataset.

**Figure S46:**
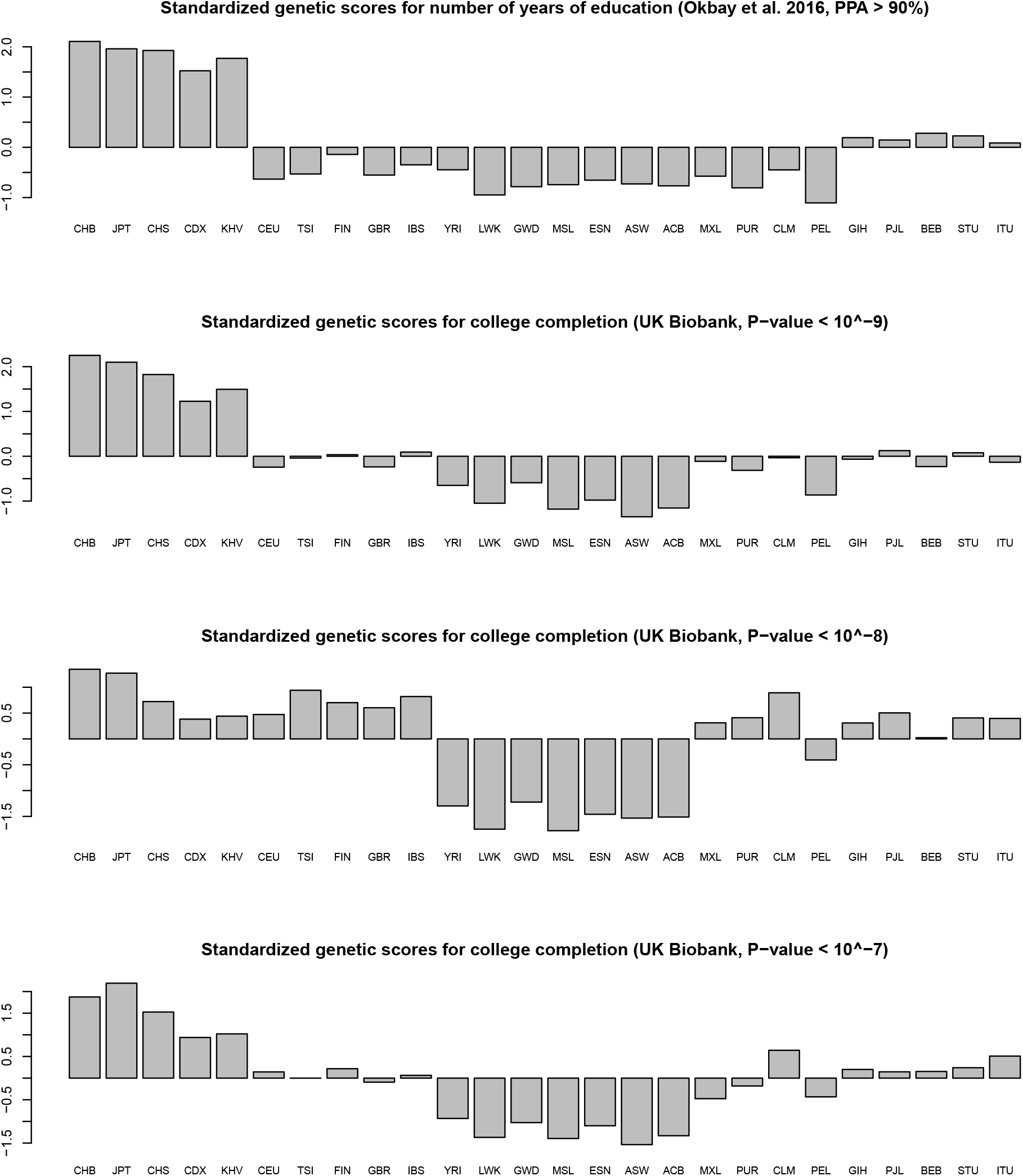
We compared the standardized population genetic scores using the ref. [46] GWAS data for number of years of education - looking at LD blocks with a posterior probability of association (PPA) > 90% (top panel), with the standardized population genetic scores using the Neale lab GWAS summary statistics obtained from the UK Biobank data, for completion of college / university degree (lower panels). We selected the SNP with the lowest P-value per LD block, and then used different P-value cutoffs to build the score: 10^−9^, 10^−8^, 10^−7^.

**Figure S47:**
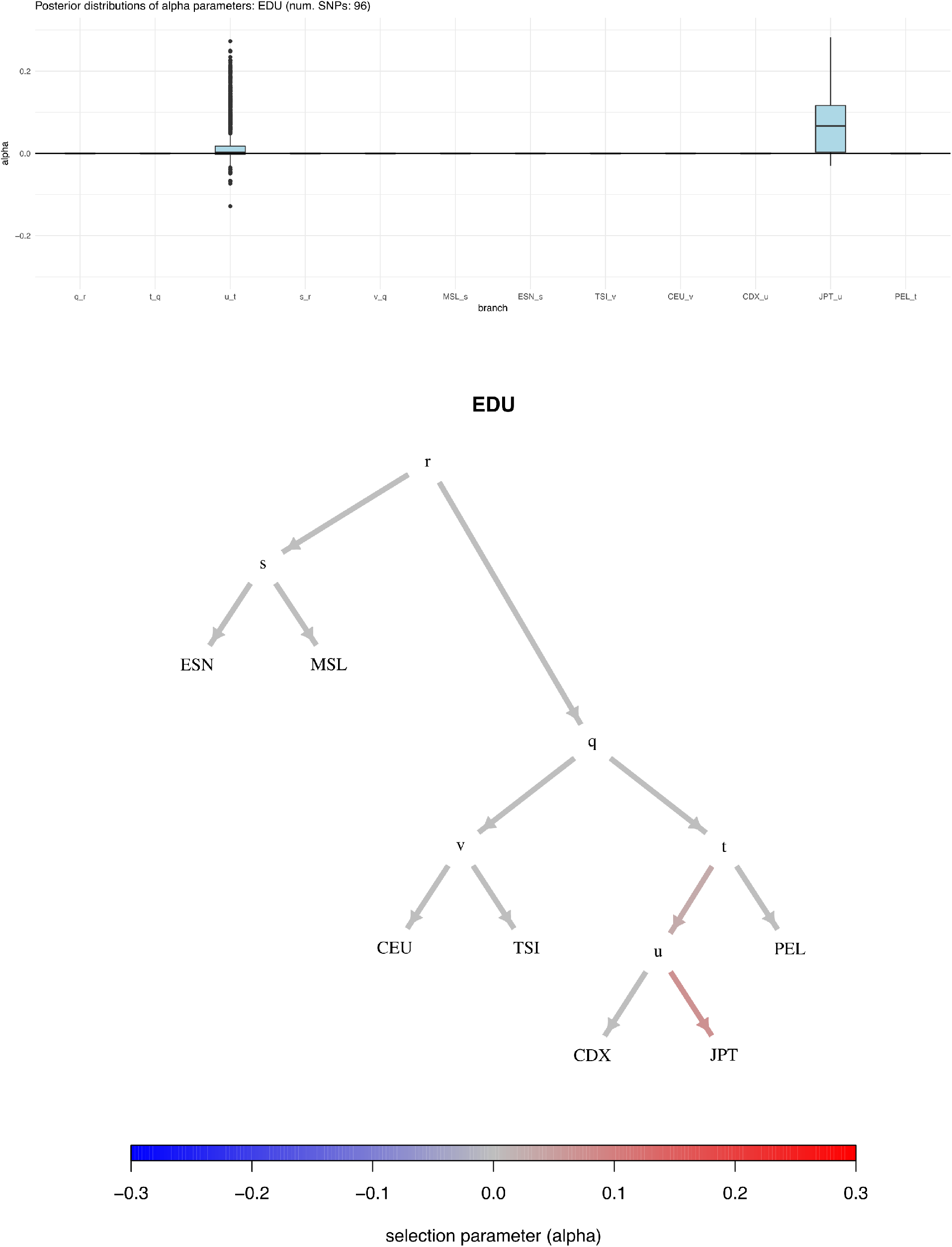
Boxplot and poly-graph of estimated *α* parameters for completion of college / university degree, using the Neale lab GWAS summary statistics obtained from the UK Biobank data. The population graph is the 7-leaf tree of the following 1000 Genomes panels: ESN, MSL, CEU, TSI, CDX, JPT and PEL.

**Figure S48:**
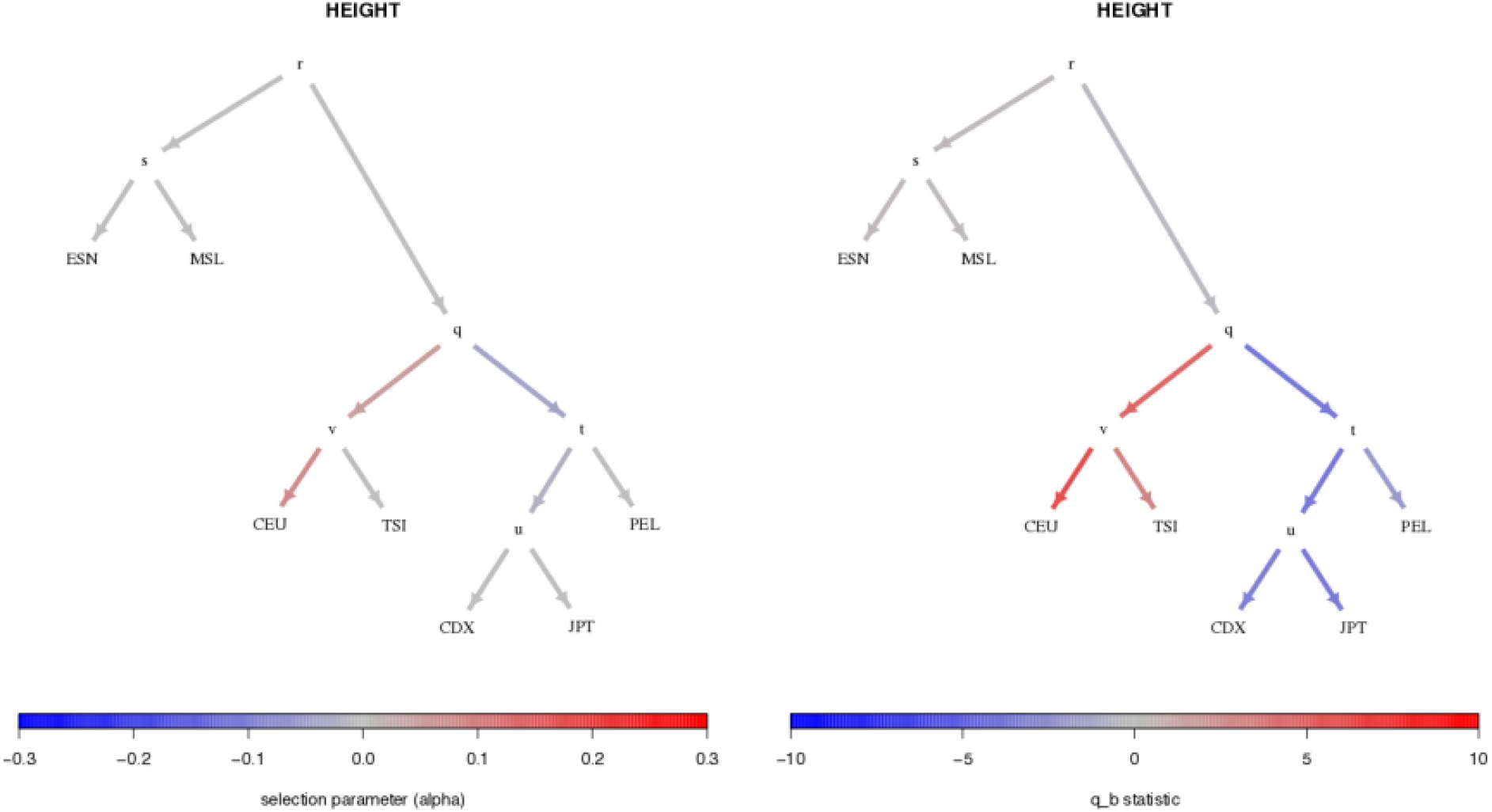
Side-by-side comparison of a graph built using the posterior estimates of the *alpha* parameters using variants associated with height (left), and a graph built using the *q_b_* statistics using the same variants (right). The color scale is arbitrary and specific to each graph.

**Figure S49:**
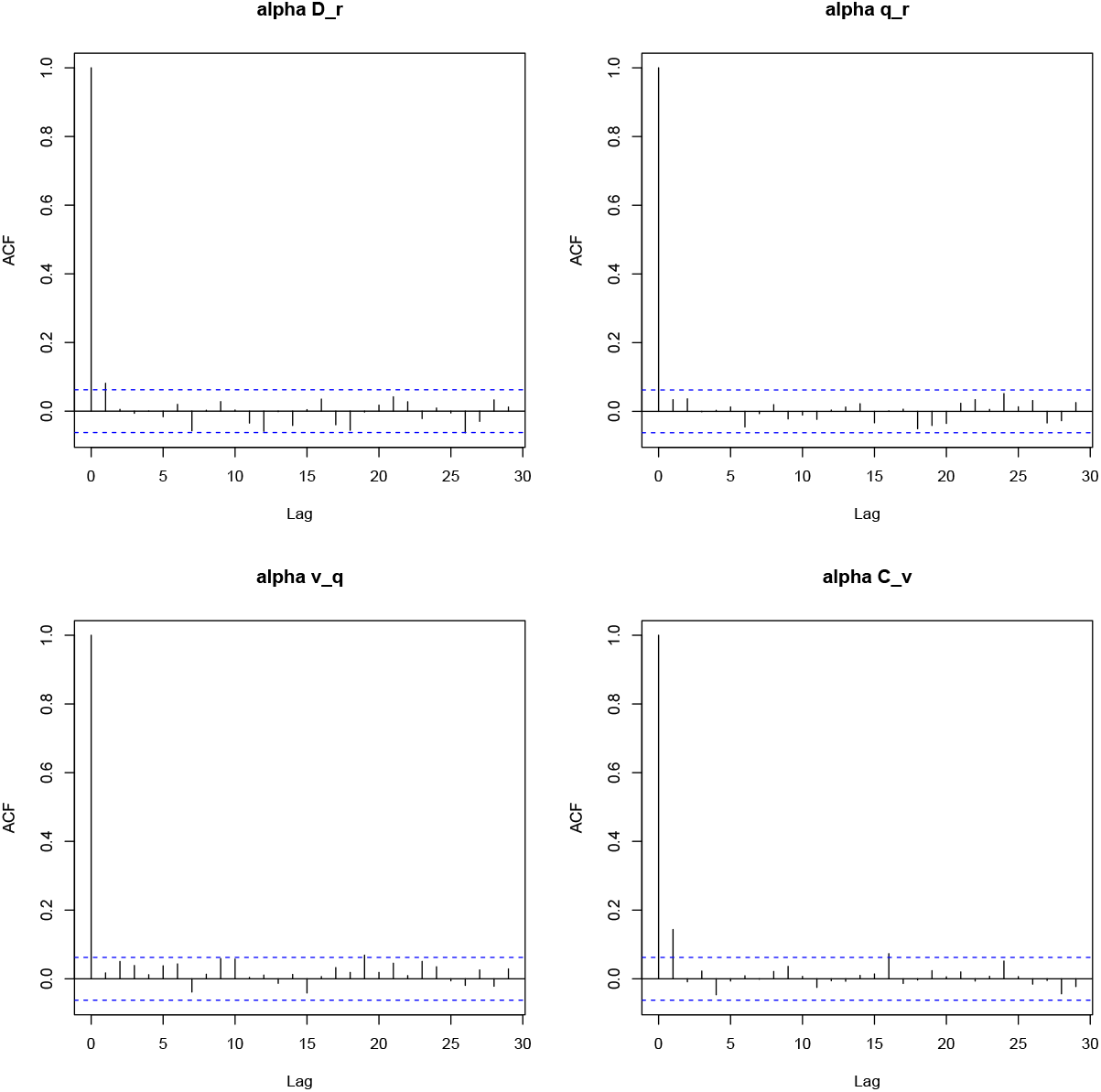
Auto-correlation plots for *α* parameters of simulation 1 of Figure S5. The rest of the *α* parameters did not pass the *Q_B_* cutoff and were therefore set to a fixed value of 0 throughout the chain.

**Figure S50:**
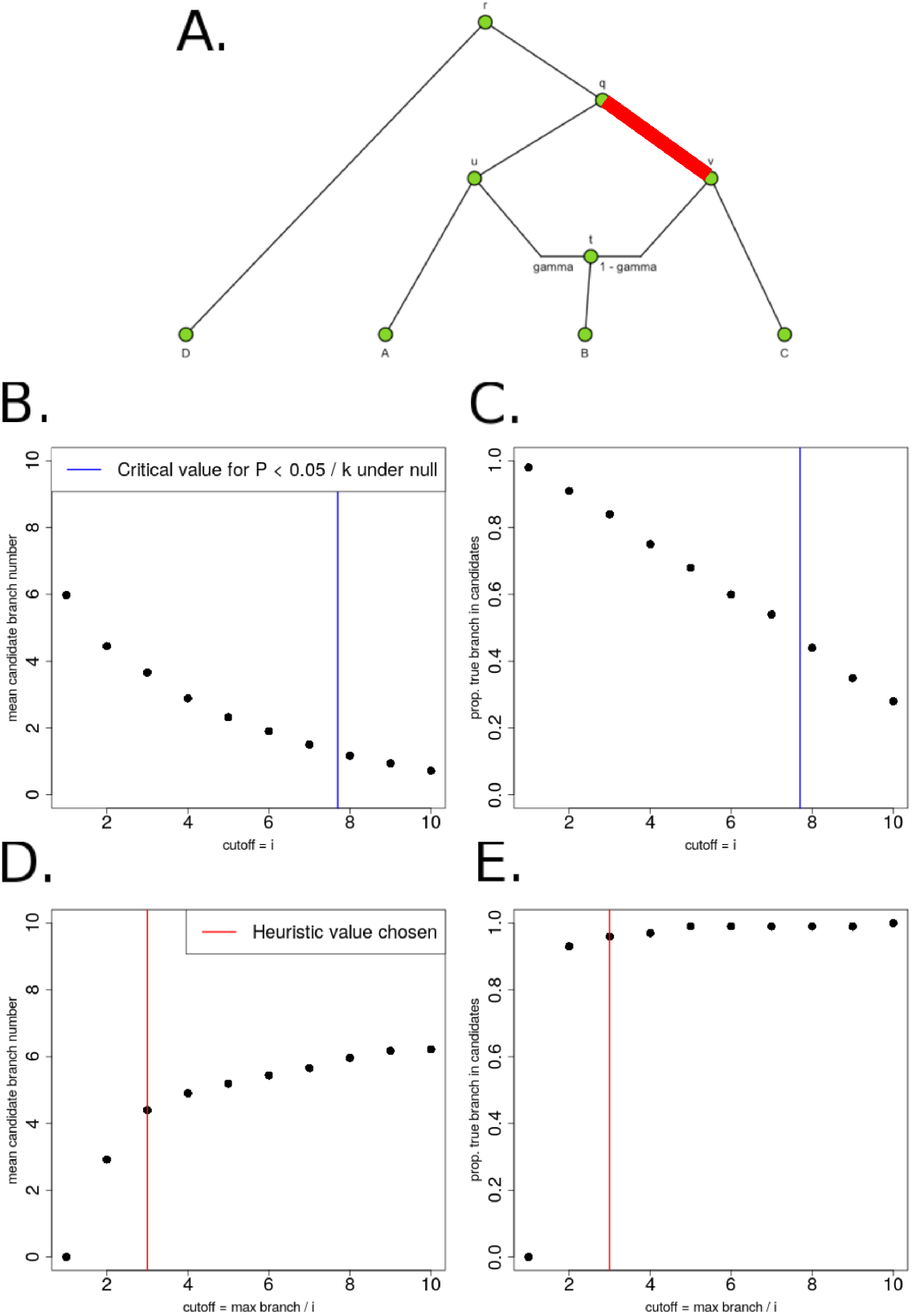
A. We produced 100 simulations of polygenic adaptation in an internal branch of a graph (v-q), each with 400 SNPs and *α* = 0.1. All branches had drift lengths equal to 0.02 B. We tested various cutoffs for *Q_B_* and plotted the average number of candidate branches that passed the cutoff, over all simulations. The blue line is the 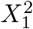 cutoff corresponding to P = 0.05/*k*, where *k* is the number of branches in the graph, under the null hypothesis of no selection. C. For each of the same cutoffs, we also plotted the proportion of simulations in which the true selected branch was among the candidate branches. D. Instead of testing constant values for the *Q_B_* cutoff, we chose the cutoff to be the maximum branch statistic in each simulation, divided by *i*, and tested various values of i. We plotted the average number of candidate branches that passed the cutoff, over all simulations. E. For the same cutoff method as in panel D, we plotted the proportion of simulations in which the true selected branch was among the candidate branches. For simulations and applications to real data, we chose a heuristic value (i=3, red line).

**Figure S51:**
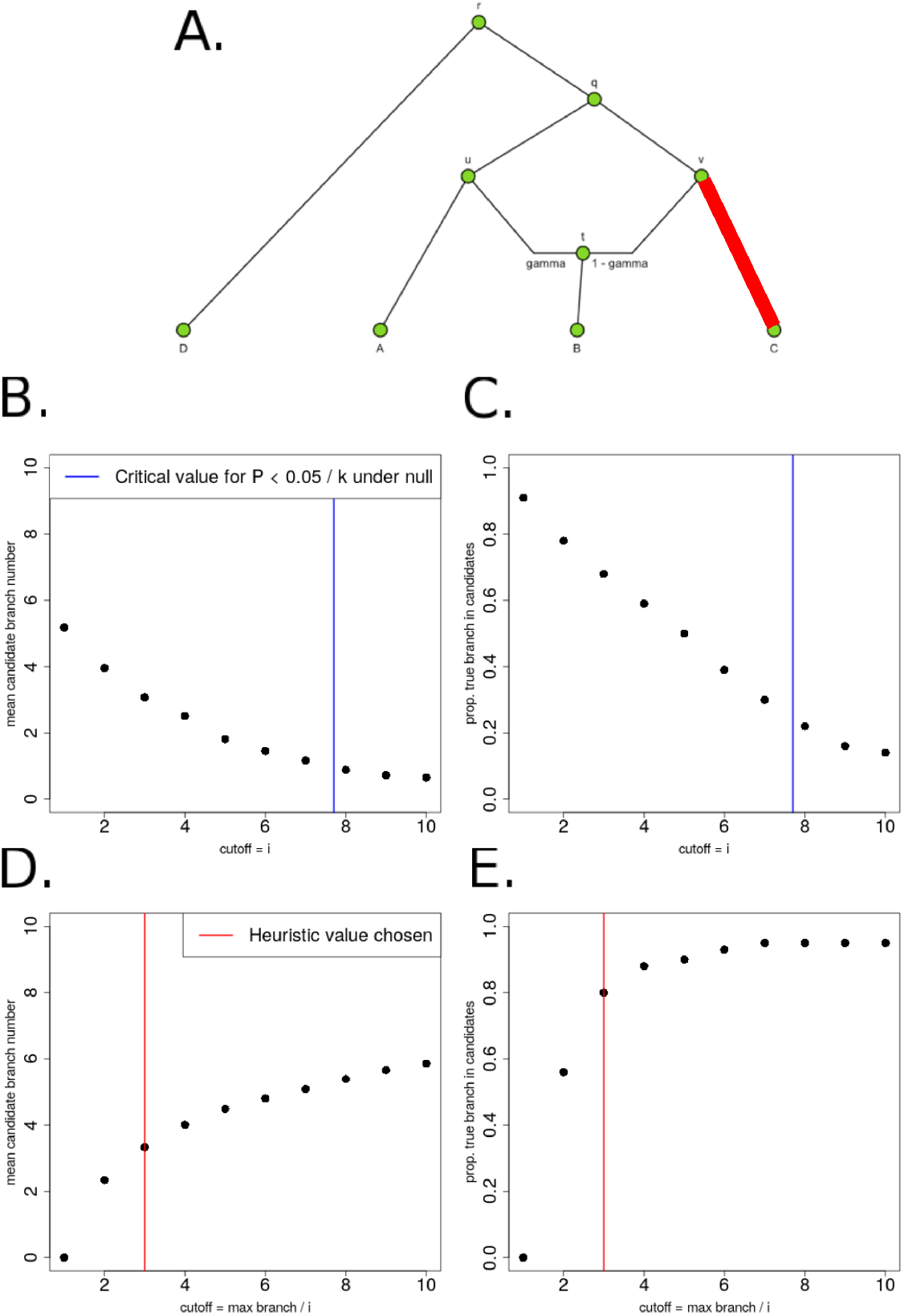
A. We produced 100 simulations of polygenic adaptation in a terminal branch of a graph (C-v), each with 400 SNPs and *α* = 0.1. All branches had drift lengths equal to 0.02 B. We tested various cutoffs for *Q_B_* and plotted the average number of candidate branches that passed the cutoff, over all simulations. The blue line is the 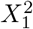 cutoff corresponding to P = 0.05/*k*, where *k* is the number of branches in the graph, under the null hypothesis of no selection. C. For each of the same cutoffs, we also plotted the proportion of simulations in which the true selected branch was among the candidate branches. D. Instead of testing constant values for the *Q_B_* cutoff, we chose the cutoff to be the maximum branch statistic in each simulation, divided by *i*, and tested various values of *i*. We plotted the average number of candidate branches that passed the cutoff, over all simulations. E. For the same cutoff method as in panel D, we plotted the proportion of simulations in which the true selected branch was among the candidate branches. For simulations and applications to real data, we chose a heuristic value (i=3, red line).

**Figure S52:**
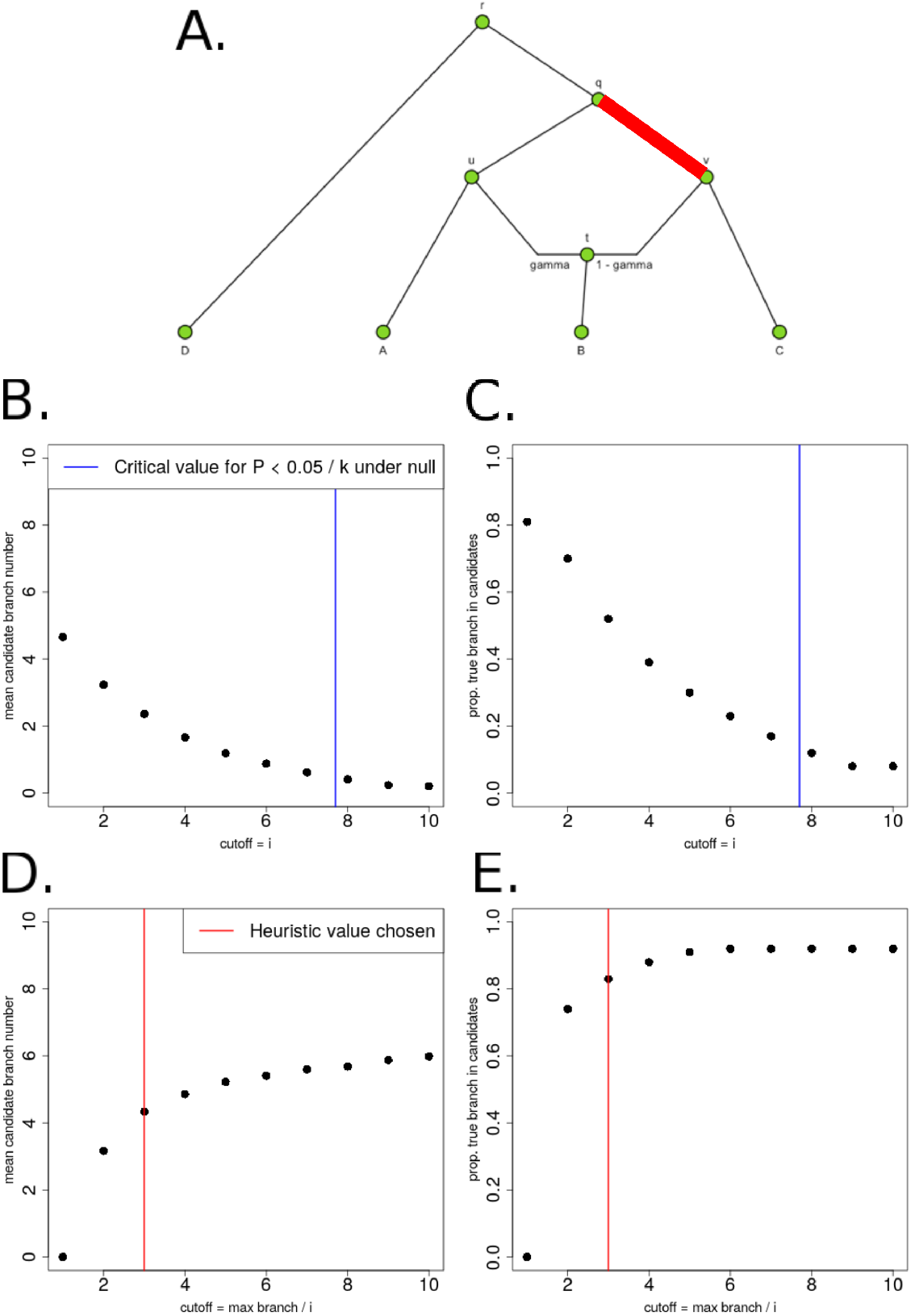
A. We produced 100 simulations of polygenic adaptation in an internal branch of a graph (v-q), each with 400 SNPs and *α* = 0.1. All branches had drift lengths equal to 0.05 B. We tested various cutoffs for *Q_B_* and plotted the average number of candidate branches that passed the cutoff, over all simulations. The blue line is the 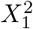 cutoff corresponding to P = 0.05/*k*, where *k* is the number of branches in the graph, under the null hypothesis of no selection. C. For each of the same cutoffs, we also plotted the proportion of simulations in which the true selected branch was among the candidate branches. D. Instead of testing constant values for the *Q_B_* cutoff, we chose the cutoff to be the maximum branch statistic in each simulation, divided by *i*, and tested various values of *i*. We plotted the average number of candidate branches that passed the cutoff, over all simulations. E. For the same cutoff method as in panel D, we plotted the proportion of simulations in which the true selected branch was among the candidate branches. For simulations and applications to real data, we chose a heuristic value (i=3, red line).

**Figure S53:**
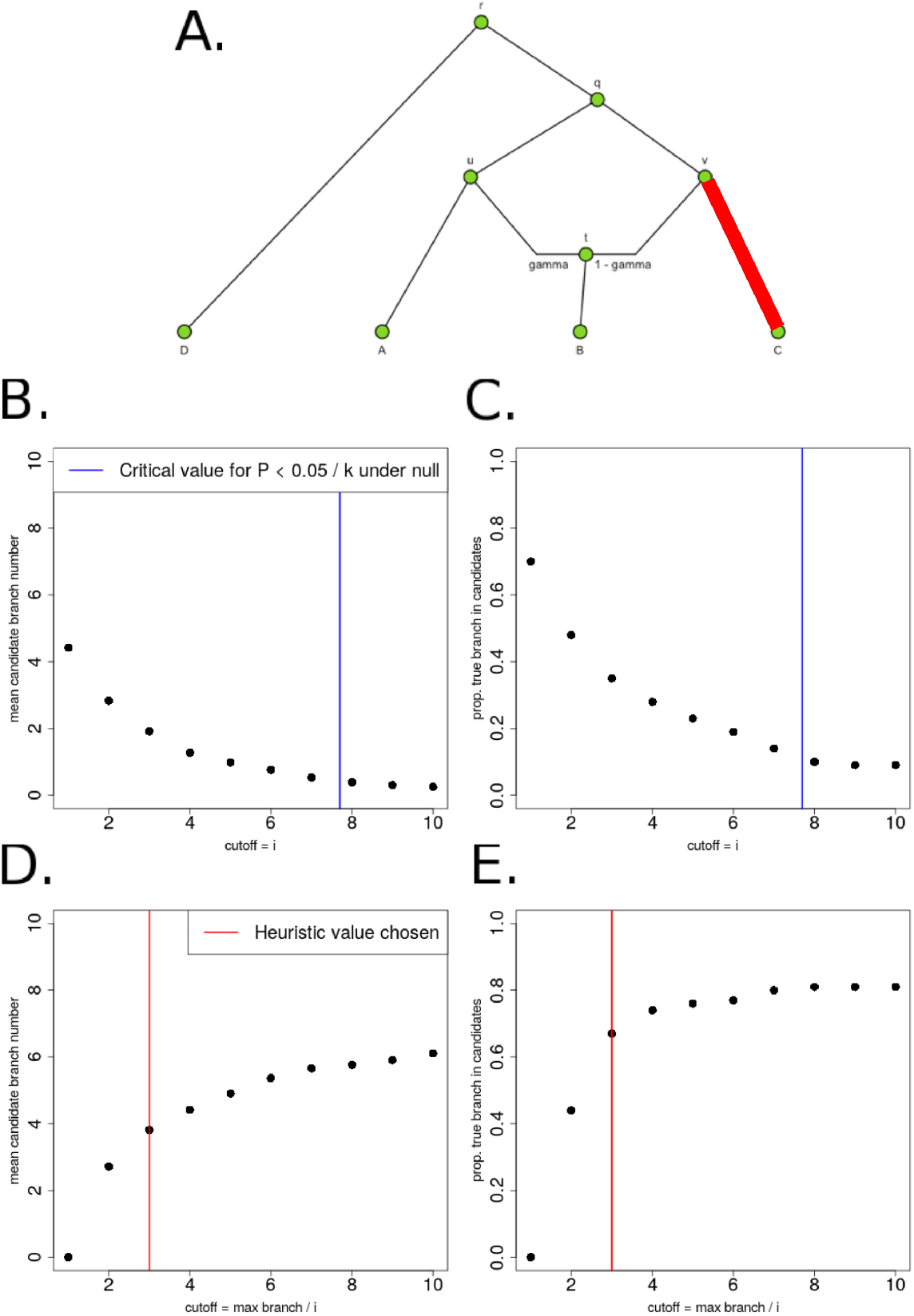
A. We produced 100 simulations of polygenic adaptation in a terminal branch of a graph (C-v), each with 400 SNPs and *α* = 0.1. All branches had drift lengths equal to 0.05 B. We tested various cutoffs for *Q_B_* and plotted the average number of candidate branches that passed the cutoff, over all simulations. The blue line is the 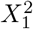 cutoff corresponding to P = 0.05/*k*, where *k* is the number of branches in the graph, under the null hypothesis of no selection. C. For each of the same cutoffs, we also plotted the proportion of simulations in which the true selected branch was among the candidate branches. D. Instead of testing constant values for the *Q_B_* cutoff, we chose the cutoff to be the maximum branch statistic in each simulation, divided by *i*, and tested various values of *i*. We plotted the average number of candidate branches that passed the cutoff, over all simulations. E. For the same cutoff method as in panel D, we plotted the proportion of simulations in which the true selected branch was among the candidate branches. For simulations and applications to real data, we chose a heuristic value (i=3, red line).

**Figure S54:**
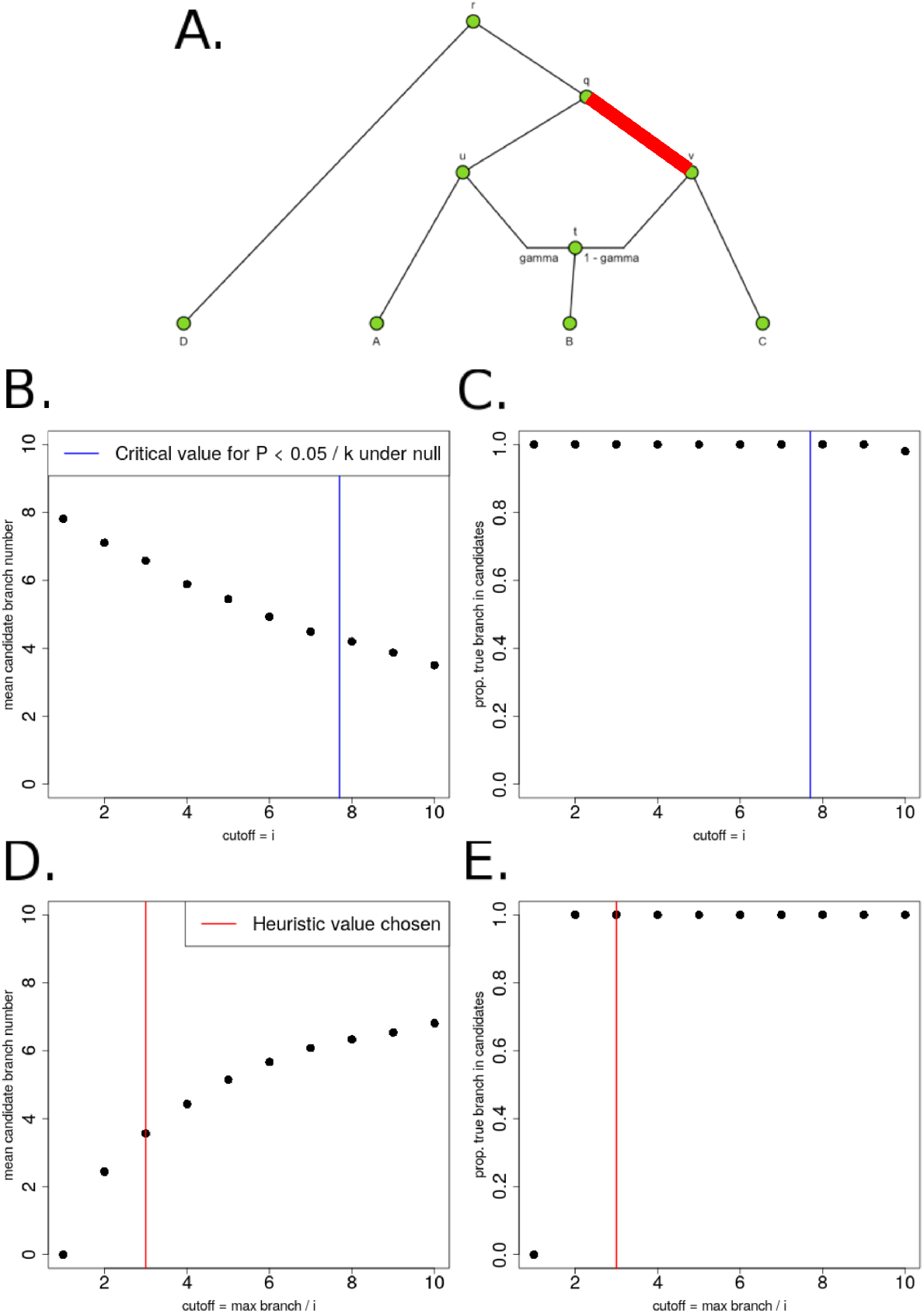
A. We produced 100 simulations of polygenic adaptation in an internal branch of a graph (v-q), each with 400 SNPs and *α* = 0.2. All branches had drift lengths equal to 0.02 B. We tested various cutoffs for *Q_B_* and plotted the average number of candidate branches that passed the cutoff, over all simulations. The blue line is the 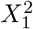 cutoff corresponding to P = 0.05/*k*, where *k* is the number of branches in the graph, under the null hypothesis of no selection. C. For each of the same cutoffs, we also plotted the proportion of simulations in which the true selected branch was among the candidate branches. D. Instead of testing constant values for the *Q_B_* cutoff, we chose the cutoff to be the maximum branch statistic in each simulation, divided by *i*, and tested various values of *i*. We plotted the average number of candidate branches that passed the cutoff, over all simulations. E. For the same cutoff method as in panel D, we plotted the proportion of simulations in which the true selected branch was among the candidate branches. For simulations and applications to real data, we chose a heuristic value (i=3, red line).

**Figure S55:**
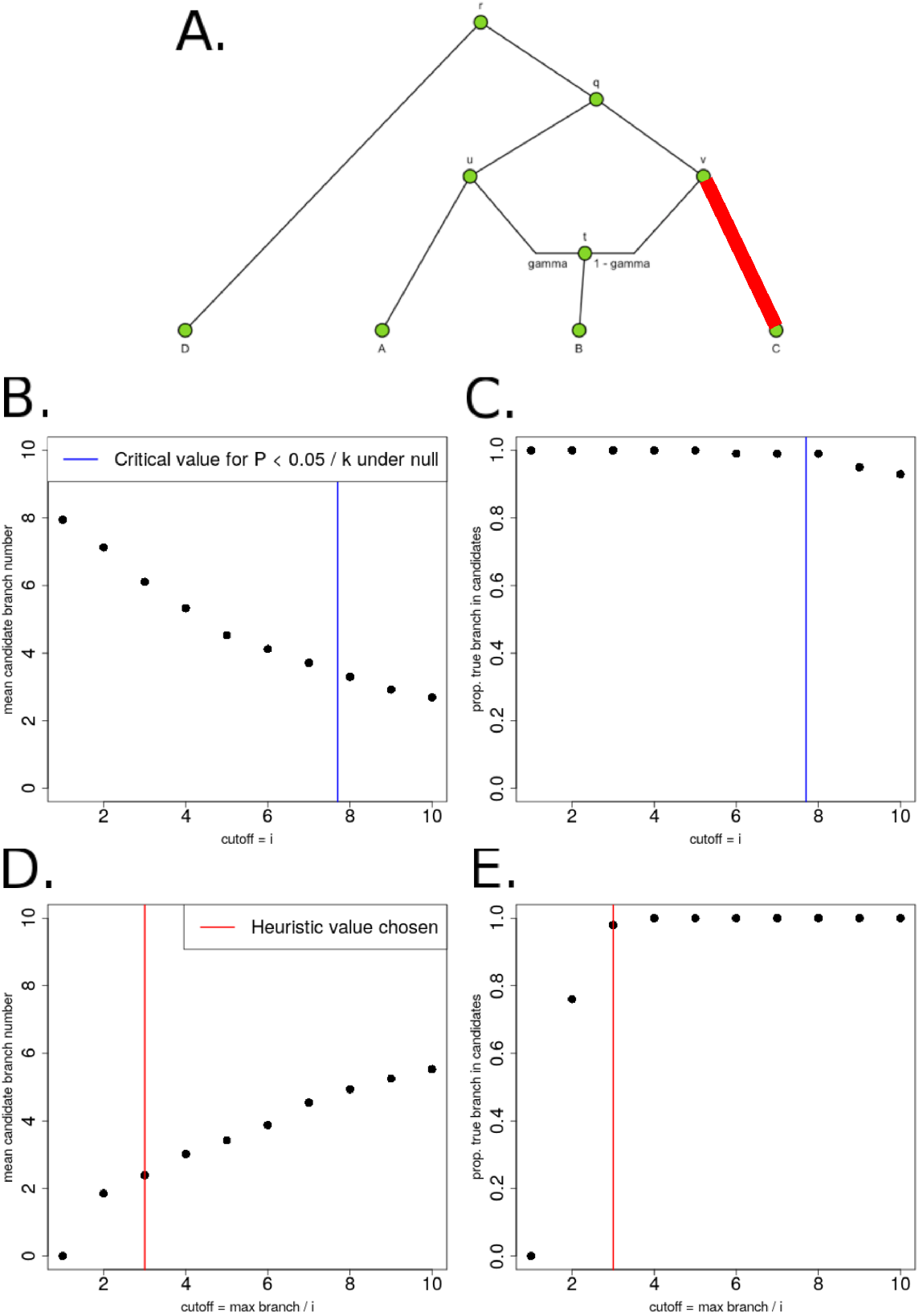
A. We produced 100 simulations of polygenic adaptation in a terminal branch of a graph (C-v), each with 400 SNPs and *α* = 0.2. All branches had drift lengths equal to 0.02 B. We tested various cutoffs for *Q_B_* and plotted the average number of candidate branches that passed the cutoff, over all simulations. The blue line is the 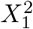 cutoff corresponding to P = 0.05/*k*, where *k* is the number of branches in the graph, under the null hypothesis of no selection. C. For each of the same cutoffs, we also plotted the proportion of simulations in which the true selected branch was among the candidate branches. D. Instead of testing constant values for the *Q_B_* cutoff, we chose the cutoff to be the maximum branch statistic in each simulation, divided by *i*, and tested various values of *i*. We plotted the average number of candidate branches that passed the cutoff, over all simulations. E. For the same cutoff method as in panel D, we plotted the proportion of simulations in which the true selected branch was among the candidate branches. For simulations and applications to real data, we chose a heuristic value (i=3, red line).

**Figure S56:**
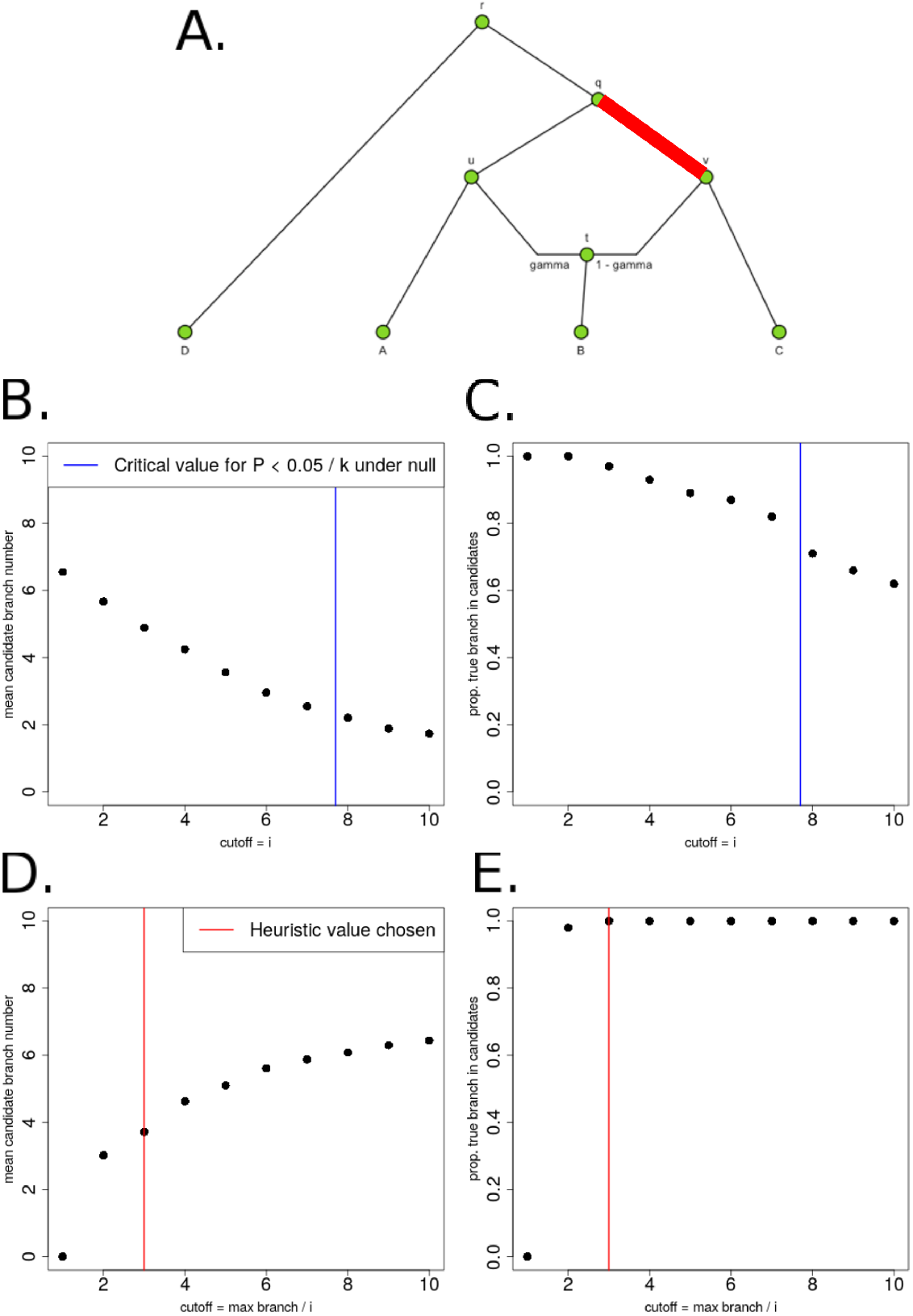
A. We produced 100 simulations of polygenic adaptation in an internal branch of a graph (v-q), each with 400 SNPs and *α* = 0.2. All branches had drift lengths equal to 0.05 B. We tested various cutoffs for *Q_B_* and plotted the average number of candidate branches that passed the cutoff, over all simulations. The blue line is the 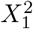 cutoff corresponding to P = 0.05/*k*, where *k* is the number of branches in the graph, under the null hypothesis of no selection. C. For each of the same cutoffs, we also plotted the proportion of simulations in which the true selected branch was among the candidate branches. D. Instead of testing constant values for the *Q_B_* cutoff, we chose the cutoff to be the maximum branch statistic in each simulation, divided by *i*, and tested various values of *i*. We plotted the average number of candidate branches that passed the cutoff, over all simulations. E. For the same cutoff method as in panel D, we plotted the proportion of simulations in which the true selected branch was among the candidate branches. For simulations and applications to real data, we chose a heuristic value (i=3, red line).

**Figure S57:**
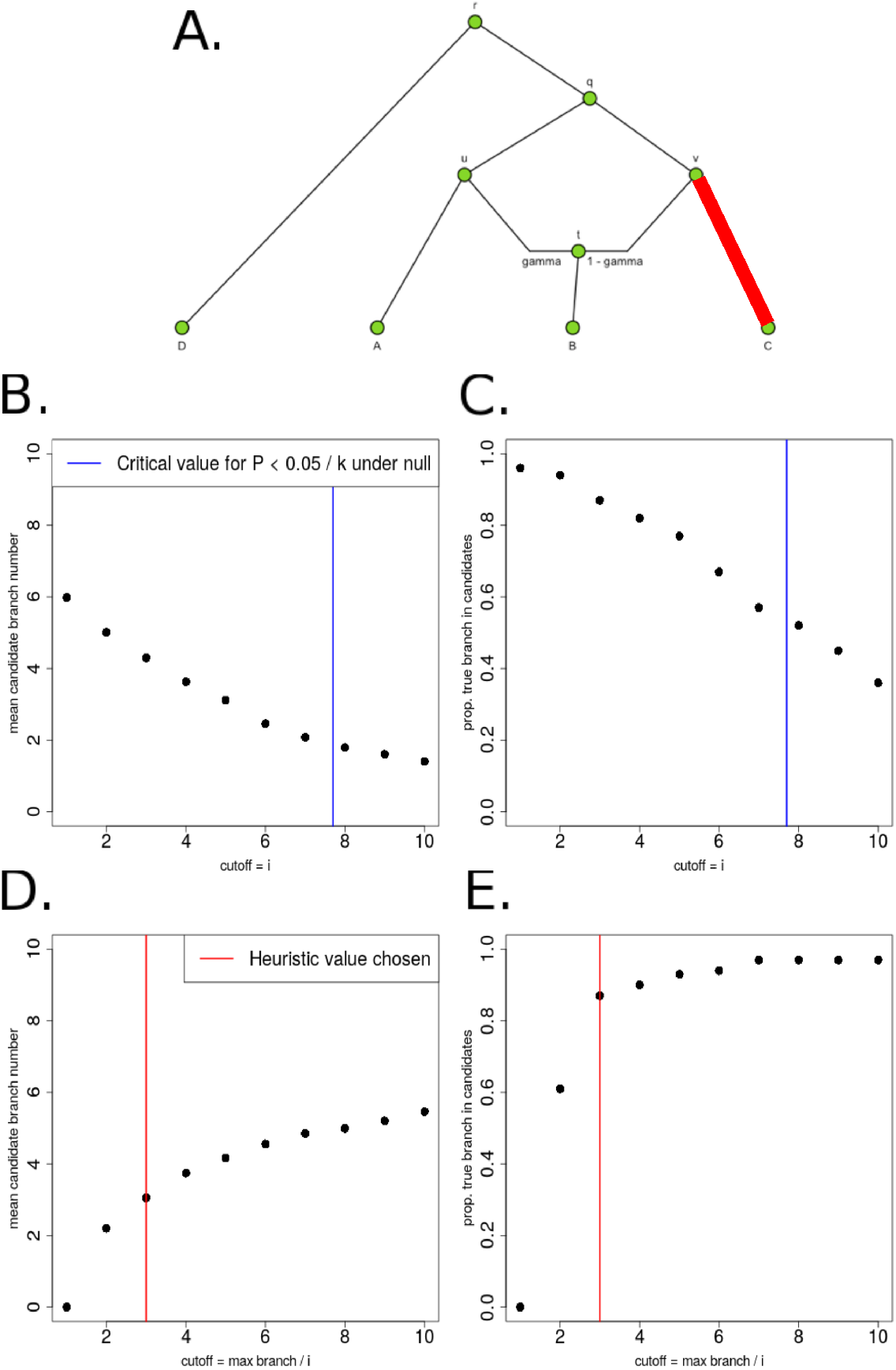
A. We produced 100 simulations of polygenic adaptation in a terminal branch of a graph (C-v), each with 400 SNPs and *α* = 0.2. All branches had drift lengths equal to 0.05 B. We tested various cutoffs for *Q_B_* and plotted the average number of candidate branches that passed the cutoff, over all simulations. The blue line is the 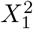 cutoff corresponding to P = 0.05/*k*, where *k* is the number of branches in the graph, under the null hypothesis of no selection. C. For each of the same cutoffs, we also plotted the proportion of simulations in which the true selected branch was among the candidate branches. D. Instead of testing constant values for the *Q_B_* cutoff, we chose the cutoff to be the maximum branch statistic in each simulation, divided by *i*, and tested various values of *i*. We plotted the average number of candidate branches that passed the cutoff, over all simulations. E. For the same cutoff method as in panel D, we plotted the proportion of simulations in which the true selected branch was among the candidate branches. For simulations and applications to real data, we chose a heuristic value (i=3, red line).

**Figure S58:**
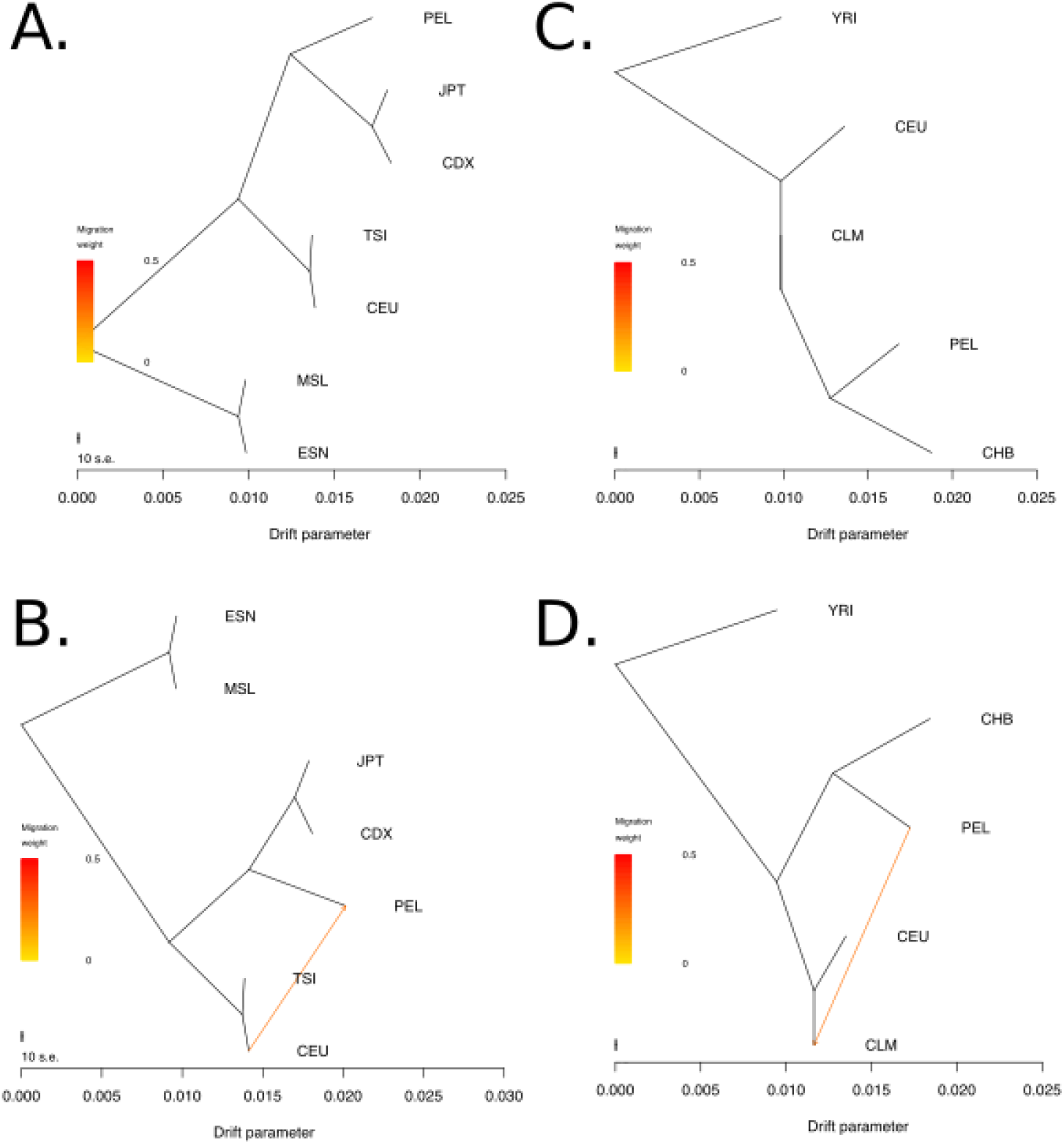
A. Topology inferred by *TreeMix* when forcing no admixture events and inputting the following populations from the 1000 Genomes Project: MSL, ESN, CDX, JPT, CEU, TSI and PEL. B. Topology inferred by *TreeMix* when inputting the same populations as in panel A, but forcing 1 migration event. C. Topology inferred by *TreeMix* when forcing no admixture events and inputting the following populations from the 1000 Genomes Project: YRI, CEU, CHB, PEL, CLM. D. Topology inferred by *TreeMix* when inputting the same populations as in panel C, but forcing 1 migration event. The trees were rooted such that the African populations were an outgroup to the rest of the populations. We note that the drift parameters outputted by *TreeMix* - unlike those outputted by *MixMapper* - are a product of the drift parameter that we use in PolyGraph and the average over the inferred heterozygosities at each branchs ancestral node, computed over all SNPs used to produce the *TreeMix* graph.

